# One-dimensional CNNs for Near-Infrared Prediction of Protein and Moisture in Cereal Grains: The Effects of Architecture and Input Preparation

**DOI:** 10.64898/2026.09.12.751160

**Authors:** Ganqi Deng, Weilai Chi, Peiqiang Yu, Fang-Xiang Wu

## Abstract

Near-infrared (NIR) spectroscopy is widely used for the rapid, nondestructive determination of constituents such as protein and moisture in cereal grains, but model comparisons in this field are often confounded by differences in the inputs received by each model. We benchmark a compact one-dimensional CNN derived by one-factor-at-a-time ablation (CNN-Baseline) and a randomly searched CNN (CNN-RS) against PLSR, SVR, XGBoost on six cereal-grain datasets (n=500-5046) under three input conditions: raw spectra, optimal preprocessing, and preprocessing plus wavelength selection. A single-filter with a kernel size of 11 of convolution and batch normalization proved sufficient. Given a unified raw input, CNN-RS was most accurate on five of the six datasets, while CNN-Baseline came within 0.003-0.017 in test *R*^2^ on the four larger datasets. PLSR led only on the smallest protein dataset (XDS, n=500) and was the most stable model. Preprocessing had almost no effect on CNN accuracy on the four larger datasets, but improved accuracy by 0.034-0.047 (CNN-Baseline) and 0.017-0.055 (CNN-RS) on the two smallest, and brought the networks to their optimum in a median of 49% fewer epochs. Therefore, replacing explicit preprocessing with learned filters was only justified when sufficient training data were available. Once each model received its own preprocessing and wavelength subset, SVR became most accurate on four of six datasets and XGBoost rose from the weakest model to within 0.01-0.03 of the best model on four of the six datasets, suggesting that the principal advantage of CNNs lied in robustness to raw input data rather than in attainable accuracy. Overall, model selection for NIR calibration of cereal grains depends more on sample size and the input preparation than on network architecture.

## 1. Introduction

Cereal grains serve as the primary dietary source of energy, protein, fiber, and essential micronutrients worldwide, which are also rich in bioactive phytochemicals that play an important role in heath maintenance (Poutanen et al., 2022). For example, barley and oat have abundant *β*-glucan (Deng et al., 2025, 2023a,b). Studies show that *β*-glucan is potentially associated with prevention of cardiovascular disease (Glenn et al., 2023; Llanaj et al., 2022; Wu et al., 2019), cancer prevention (Li et al., 2022; Paudel et al., 2021), and reducing blood cholesterol and coronary heart disease (CHD) incidence reduction (Hu et al., 2022; Li et al., 2024; Tosh and Bordenave, 2020). However, the full nutritional value of grains remains largely untapped due to analytical bottlenecks. Traditional wet chemistry methods of nutritional analysis are slow, costly, labor-intensive, destructive, and unfriendly-environmental (Deng et al., 2020; Shi and Yu, 2017). This hinders the process of providing comprehensive and rapid assessment information for decision-making across the food chain, from breeding of more nutritious crop varieties to real-time quality evaluation in food processing. Currently, around 41% of grains are used for human consumption and more than 35% are used for animal feed (Poutanen et al., 2022). The quality assessment of grains is not only crucial for human but also vital for animal health, because animal’s heath is closely linked to human food chain. In livestock production, the nutritional composition of feed grains is one of the factors determining animal growth, immune system and overall welfare (Garutti et al., 2022). Poor feed quality could lead to reduced productivity and increased susceptibility to disease. Therefore, robust systems for rapid assessment of grain nutritional contents can support the development of high-quality animal feed, promote healthier livestock, and contribute to a safer, more sustainable food supply for humans. Near-infrared (NIR) spectroscopy has become one of the most widely used techniques for the rapid, non-destructive analysis of the chemical composition of food and agriculture products (Roberts et al., 2004). For grains such as wheat, barley, and maize, protein and moisture contents are the key parameters for quality control and trade. However, routine determination using classical wet chemical methods is slow, labor-intensive, and destructive. NIR spectroscopy can detect the overtone and combination absorptions of C-H, N-H and O-H bands, enabling the estimation of these nutritional components within seconds based on a single spectrum (Manley, 2014). This makes it a standard tool for grain reception, plant breeding, and process control. Because NIR spectroscopy exhibits high collinearity and is susceptible to light scattering and baseline shifts, its quantitative calibration also relies on chemometric modeling. Before the partial least squares regression (PLSR) model became widely used, principal component regression (PCR) was the standard method for spectral analysis (Malinowski, E. R., 2002). This method involved first performing a principal component analysis (PCA) decomposition on the X of spectral data to extract the first few principal components (PCs) that contributed most. These principal components are orthogonal to one another (uncorrelated) and successfully reduce the dimensionality of several thousand spectral peaks to a few uncorrelated variables. Then, these principal components are used as new independent variables in a multiple linear regression analysis with the target chemical values Y (such as protein and moisture content). However, when extracting principal components, PCA considers only the spectral data X itself and seeks the direction with the greatest spectral variability (variance). The direction with the greatest variability in the spectrum may be caused by instrument noise, scattering, or temperature fluctuations, and is not necessarily related to the chemical composition Y to be predicted. Therefore, PCA may filter out features that are critical for predicting Y but have weak signals, deeming them “too small” to be significant. PLSR addresses this limitation by extracting latent variables using information from both **X** and *Y*, typically by maximizing their covariance (Geladi and Kowalski, 1986). It remains widely used for NIR calibration. To obtain a more accurate PLSR model, the raw spectra are typically subjected to mathematical preprocessing to suppress scatter and baseline effects (Rinnan et al., 2009), such as standard normal variate transform (Barnes et al., 1989), multiplicative scattering correction (Geladi et al., 1985), or Savitzky-Golay derivative (Savitzky and Golay, 1964). However, selecting the optimal preprocessing method relies heavily on experience and experiments. It depends on the dataset and analyst’s experience, and an inappropriate choice may remove useful information or introduce artifacts. In addition, standard linear PLSR cannot directly represent nonlinear relationships between spectra and reference values.

Therefore, nonlinear machine learning methods have been applied to near-infrared spectroscopy data, including support vector regression (SVR) (Smola and Schölkopf, 2004) and extreme gradient boosting (XGBoost) (Chen and Guestrin, 2016). Recently, deep learning such as one-dimensional convolutional neural networks (1D-CNNs) has also attracted attention in the field of food spectral analysis (Li et al., 2025; Liang et al., 2022; Luo et al., 2024; Mishra et al., 2022; Zhang et al., 2021). One of the CNN’s greatest advantages is that they can learn task-specific spectral representations directly from raw or minimally processed spectra, potentially reducing their dependence on manually selected preprocessing (Acquarelli et al., 2017; Cui and Fearn, 2018; Zhang et al., 2019).

Although CNNs have been applied to NIR calibration, there is still little standardization in practical design. The reported architectures vary greatly in depth and width, and often directly adapted from image classification networks, without a clear understanding of which components drive performance. The individual effects of architecture and training hyperparameters are rarely systematically examined through ablation studies, such as the number of convolutional layers and filters, kernel size, batch normalization or dropout, and the optimizer and learning rate. Furthermore, it is well known that deep learning models require large data sets (Zhang et al., 2019), while most studies either reported a single dataset or a set of uncorrelated datasets differing in instrument, composition, and validation design at the same time. Currently, it is only possible to infer across studies rather than measure directly within a single study if the advantages of deep learning depend on the size of calibration sample.

To address these gaps, this study develops and benchmarks one-dimensional convolutional neural networks (1D-CNNs) for predicting protein and moisture content in cereal grains using NIR spectra. The study makes three contributions. Firstly, a systematic, one-factor-at-a-time ablation study is implemented to investigate the effects of thirteen architecture and training hyperparameters. Based on the principle of parsimony, a minimal CNN architecture is identified and designed as CNN-Baseline. Additionally, a 1D-CNN has also been developed that can automatically search for the optimal hyperparameter configuration for a given dataset, named as CNN-RS. This makes it easier to transfer 1D-CNN across different grain datasets, eliminating the need to repeatedly perform manual grid searches for optimal parameters whenever the dataset changes. Secondly, all five models are benchmarked across six datasets under three input conditions (raw spectra, optimal preprocessing, and combination of preprocessing and wavelength selection), ensuring that each model is evaluated both under standardized input conditions and after undergoing the same degree of optimization. The preprocessing methods for the CNNs are selected independently rather than from chemometric models. Thirdly, it provides practical guidance on these three factors: which architecture is sufficient, the impact of data preprocessing on various models, and at what sample size the ranking between model changes.

## 2. Methods

### 2.1. Datasets

Models were developed and evaluated based on six near infrared (NIR) spectroscopic datasets extracted from the sensAIfood CRAW dataset (sensAIfood (IG19145), 2025), which hadn’t been used for a benchmarking study of this kind. The dataset covered three types of grains (wheat, barley and maize), two quality contents (protein and moisture, both expressed in %), and two spectrometers (NIRS5000 and XDS). The NIRS5000 spectrum comprised 700 wavelengths over 1100-2498 nm, while the XDS spectrum comprised 1050 wavelengths over 400-2498 nm.

Six datasets were wheat protein (NIRS5000, n=5046), barley protein (NIRS5000, n=2096), wheat protein (XDS, n=500), wheat moisture (NIRS5000, n=3305), barley moisture (NIRS5000, n=985), and maize moisture (NIRS5000, n=585). These datasets covered a wide range of compositional distributions and sample sizes (500-5046), which allowed for an assessment of the generalizability of each modeling method across grain types, compositions, tools, and dataset sizes. Table 1 summarized the basic information for each dataset.

**Table 1:** Overview of the six near-infrared datasets used in this study.

| Dataset | Spectrometer | Wavelengths | Spectral range (nm) | Samples ($n$) | Trait range (%) | Mean (%) | SD (%) |
| --- | --- | --- | --- | --- | --- | --- | --- |
| Wheat protein | NIRS5000 | 700 | 1100–2498 | 5046 | 6.44–20.25 | 11.96 | 1.37 |
| Barley protein | NIRS5000 | 700 | 1100–2498 | 2096 | 7.34–16.98 | 11.23 | 1.57 |
| Wheat protein | XDS | 1050 | 400–2498 | 500 | 9.00–14.03 | 11.27 | 0.98 |
| Wheat moisture | NIRS5000 | 700 | 1100–2498 | 3305 | 9.71–18.54 | 13.33 | 1.18 |
| Barley moisture | NIRS5000 | 700 | 1100–2498 | 985 | 9.90–17.43 | 13.02 | 1.64 |
| Maize moisture | NIRS5000 | 700 | 1100–2498 | 585 | 8.10–19.20 | 12.71 | 1.64 |

*Note:* SD = standard deviation (sample, $n - 1$ ). Trait values are protein or moisture content in percent (%). NIRS5000 and XDS denote the two spectrometers in the sensAlfood CRAW dataset; spectra contain 700 (NIRS5000) or 1050 (XDS) wavelengths.

### 2.2. Data partitioning

Each dataset was split into training, validation and test sets in a 70/15/15 ratio. This splitting was performed in two stages using the train-test-split function in scikit-learn, 15% of the samples were set aside as the test set first, then 17.6% of the remaining 85% were set aside as the validation set, resulting in a total of 15% of the original data. In both stages, a fixed random seed (42) was set and shuffling, so that every model was developed and evaluated on identical partitions. Figure 1 showed the splits of the representative wheat protein (NIRS5000) dataset, along with the corresponding spectral and compositional distributions.

**Figure 1:**
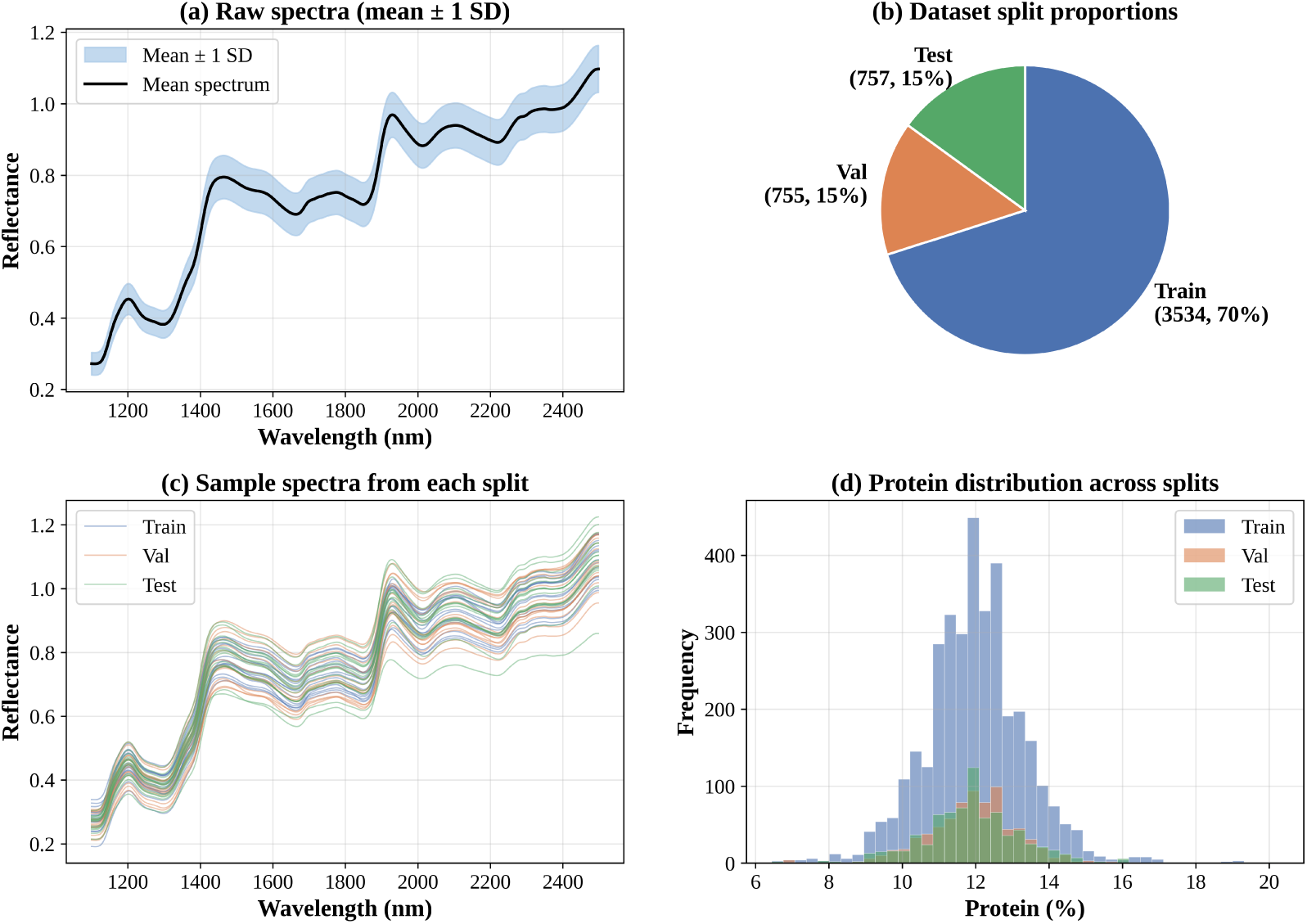
Spectral characteristics and data partitioning of the representative wheat protein (NIRS5000) dataset (*n* = 5046). **(a)** The average near-infrared reflectance spectrum (black line) and the mean *±*1 SD band (shaded area) illustrate the overall spectral profile and sample variability over the 1100–2498 nm range (700 wavelengths). **(b)** The training, validation, and test subsets (70/15/15) contain 3534, 755, and 757 samples, respectively. **(c)** Representative raw spectra randomly selected from each subset indicate that the three subsets share consistent spectral profiles. **(d)** The distributions of protein content (%) across the three subsets confirm that the random partition preserves comparable target distributions. The same 70/15/15 partitioning procedure—first separating the test set and then the validation set, using a random seed of 42—was applied to all six datasets; only this representative dataset is shown.

### 2.3. Input preparation

Three operations were performed between the raw spectra and the model input, including spectral preprocessing, feature scaling, and wavelength selection.

#### 2.3.1. Spectral preprocessing

Spectral preprocessing was performed row by row, applying the transformation to each spectrum using only its own values. Twelve strategies were compared: no preprocessing (raw spectra), baseline offset correction, second-order polynomial detrending, standard normal variate (SNV), multiplicative scatter correction (MSC), first and second order Savitzky-Golay derivatives, SNV followed by detrending (SNV-detrend), SNV followed by derivatives (SNV-FD, SNV-SD), and derivatives followed by SNV (FD-SNV, SD-SNV).

Baseline correction eliminated linear or nonlinear background slopes and vertical shifts caused by factors such as instrument drift or environmental fluctuations. Standard normal variate (SNV) and multiplicative scatter correction (MSC) mitigate physical light scattering causing from changes in particle size distribution and corrected residual baseline offsets. The Savitzky-Golay derivative, combined with a smoothing step, was applied to suppress random instrumental noise, highlighted subtle peak shapes, improved peak resolution, and preserved relative peak intensity information. In addition, detrending was employed to correct for baseline shifts and curves commonly found in densely packed or powdered samples. In each case, the criterion was the coefficient of determination for the validation set. These preprocessing methods could optimize spectra quality and remove non-chemical interference (Gauglitz and Moore, 2014; Manley, 2014; Moros et al., 2010).

Feature scaling was then performed on a column-by-column basis, normalizing each wavelength in the sample to have a mean of zero and a variance of one. A StandardScaler was fitted separately on the training partition, independently for each wavelength, and was applied consistently to the validation and test partitions. Each model under every configuration was scaled, including those that had not underwent spectral preprocessing.

#### 2.3.2. Wavelength selection

Wavelength selection was performed within the PLSR pipeline. Three types of subsets were compared: the full spectrum; an importance-based subset (retaining the top 5-25% of wavelengths ranked by the magnitude of the PLS regression coefficients); and a subset restricted to absorption regions associated with proteins or water. The subsets were selected using the validation set. PLSR ranked the wavelengths by the magnitude of the PLS regression coefficients, and more details were described in Section 2.4. SVR, XGBoost, CNN-Baseline and CNN-RS utilized the wavelength subset selected by PLSR on the validation set of the same dataset.

### 2.4. Partial least squares regression (PLSR)

PLSR (Geladi and Kowalski, 1986) was adopted as the chemometric reference method. For each dataset, the model was optimized in two sequential stages. First, the twelve spectral preprocessing strategies described in Section 2.3.1 were compared with the number of latent variables optimized for each strategy by minimizing validation RMSEP, up to a maximum of twenty. The strategy yielding the highest coefficient of determination (*R*^2^) on the validation set was retained. Second, regression coefficient analysis (RCA) based on the PLS algorithm was performed under that strategy to rank wavelengths by the importance of their regression coefficient, and wavelength subset yielding the highest *R*^2^ of validation set was selected for the final model.

### 2.5. Support vector regression (SVR) and extreme gradient boosting (XGBoost)

Two machine-learning regression methods were used for comparison, support vector regression (SVR) and extreme gradient boosting (XGBoost). In the main comparison, both models were trained on raw spectra with column-wise scaling only, without any additional spectral preprocessing. CNNs also followed the same protocol. In the SVR model, a radial basis function (RBF) kernel was utilized. Three hyperparameters were optimized via exhaustive grid search: the regularization parameter *C*, controlling the trade-off between model complexity and tolerance of training errors {0.1, 1, 10, 50, 100, 500, 1000, 5000}; the kernel coefficient *γ*_SVR_, determining the influence radius of a single training sample {scale, auto, 10^−4^, 10^−3^, 10^−2^, 10^−1^, 1, 10}; and the width *ε* of the insensitive tube, within which prediction errors are not penalized {0.01, 0.05, 0.10, 0.15, 0.20, 0.30}. The yielded in a total of 8 × 8 × 6 = 384 candidate combinations. As for XGBoost model, the randomized search of 150 sampled configurations was used instead of an exhaustive grid search because of the larger hyperparameter space. The candidate values that drawn from discrete grids were as follows: the number of boosting rounds {50, 100, 200, 300, 500, 800, 1000, 1500}; the learning rate {0.005, 0.01, 0.02, 0.05, 0.10, 0.15, 0.20, 0.30}; the maximum tree depth {3, 4, …, 10, 12, 15}; the minimum child weight {1, 2, 3, 5, 7, 10, 15}; the minimum split-loss reduction *γ*_XGB_ {0, 0.05, 0.10, 0.20, 0.50, 1, 2}; and from continuous intervals for the row and column subsampling ratios [0.6, 1.0]; the *L*_1_- and *L*_2_-regularization coefficients [0, 2] and [0.5, 20], respectively. Trees were grown using a histogram-based algorithm with a squared-error objective.

In both cases, the hyperparameters were selected by five-fold cross-validation on the training set to minimize the mean squared error. Tuning and evaluation were repeated 25 times with seeds 42-66. The seeds varied the cross-validation folds (SVR) and the stochastic components of the model (XGBoost), while the training/validation/test splits remained fixed. Performance was reported as the mean *±* standard deviation across the 25 runs. Since using the uniform raw inputs underestimated the accuracy attained by these methods, two additional configurations were evaluated: one in which each model received its own optimal preprocessing strategy, selected independently on the validation set from the twelve strategies of Section 2.3.1, and another in which that preprocessing combined with the wavelength subsets selected by PLSR in Section 2.3.2.

### 2.6. One-dimensional convolutional neural network (1D-CNN)

#### 2.6.1. Architecture optimization by ablation studies

To determine the optimal configurations for the deep learning framework, the 1D-CNN architecture and its associated training hyperparameters were identified through a systematic and sequential one-factor-at-a-time (OFAT) ablation study. This optimization process was conducted using a representative wheat protein (NIRS5000, *n* = 5046) dataset as the calibration baseline. Each factor was varied sequentially while other factors remained at their baseline values, and the candidate factors were compared based on the validation-set *R*^2^. The selected setting was then carried forward to the subsequent stage.

##### Convolutional block

Depth of sequential convolutional layers increased from 1 to 11. To maintain experimental control during the depth evaluation, each additional layer was configured with a fixed density of 16 filters, a kernel size of 5 and no pooling layers. Filter density in the first convolutional layer was evaluated over {1, 8, 16, 32, 64, 128, 256}. Kernel size was investigated using odd values {3, 5, 7, 9, 11, 13, 15, 19}.

##### Fully connected block

Layer depth and scaling profiles were tested from 1 to 5 layers, including (64), (128, 64), (128, 64, 32), (256, 128, 64, 32), (512, 256, 128, 64, 32), and (256, 128, 64, 32, 16). Within a fixed three-layer architecture, dimensional capacity was explicitly evaluated over five discrete size configurations: small (32, 16, 8), medium (64, 32, 16), large (128, 64, 32), extra-large (256, 128, 64), and the original baseline configuration (32, 18, 12).

##### Regularization

Dropout rate was tested over {0, 0.1, 0.2, 0.3, 0.4, 0.5}. The presence and absence of batch normalization were compared, and the *L*_2_ weight-decay was evaluated across {0, 0.001, 0.005, 0.01, 0.05}.

##### Optimization

The initial learning rate was evaluated over {5×10^−5^, 10^−4^, 5 × 10^−4^, 10^−3^, 2.5 × 10^−3^, 5 × 10^−3^} with the dynamic learning rate scheduler disabled to ensure that the effect of the initial step size was isolated. Adaptive moment estimation (Adam) optimizer was compared against traditional stochastic gradient descent (SGD) optimizer with a fixed momentum of 0.9. Mini-batch size was tested over {16, 32, 64, 128, 256, 512}, which required the data loading pipeline to be reconfigured for each setting.

##### Feature mapping

Six activation functions were systematically evaluated: Rectified Linear Unit (ReLU), Exponential Linear Unit (ELU), Leaky ReLU (with a negative slope parameter *α* = 0.1), Hyperbolic Tangent (Tanh), Scaled Exponential Linear Unit (SELU), and Gaussian Error Linear Unit (GELU). Five downsampling strategies for convolutional features were tested: a baseline without pooling; 1D max-pooling blocks with kernel size 2; 1D average-pooling blocks with kernel size 2; wider 1D max-pooling blocks with kernel size 4; and a wider 1D max pooling block with kernel size 4, and a multi-layer deep max pooling strategy involving two consecutive convolutional-pooling blocks with pooling size 2.

Candidate models from each individual ablation scan were evaluated and compared based on their performance on the validation set, specifically using the validation coefficient of determination (*R*^2^). The final network topology followed the principle of parsimony (Ockham’s Razor). This strategy successfully yielded a highly compact model while ensuring that any sacrifice in predictive power remained within a negligible range relative to the absolute best observed configuration.

#### 2.6.2. Comparison of CNN-Baseline and CNN-RS

The optimized network architecture derived from the ablation study was designated CNN-Baseline. The network was initialized with a single one-dimensional convolutional layer with one single filter of kernel size of 11, and “same” padding to preserve the spatial boundary dimensions of the input vector. This was followed by a batch normalization layer, an exponential linear unit (ELU) activation function, and a one-dimensional average-pooling layer with a pool size and stride of 2. The pooled feature map was then flattened and passed through three fully connected layers containing 32, 18, and 12 hidden units, respectively. Each fully connected layer had an independent batch normalization step and an ELU activation function. The output layer was a single linear output neuron, which was dedicated to the continuous regression prediction of the target component percentage (protein or moisture content, expressed as a percentage). To maintain information density within this compact design, the final model used neither dropout nor weight decay. When processing NIRS5000 inputs with 700 wavelengths, this lightweight network contained approximately 12,205 trainable parameters. For the XDS instrument with an input of 1050 wavelengths, the network scaled to 17,805 trainable parameters. The architecture was illustrated schematically in Fig. 2a, and its complete architecture and training configuration were provided in Table 2.

**Figure 2:**
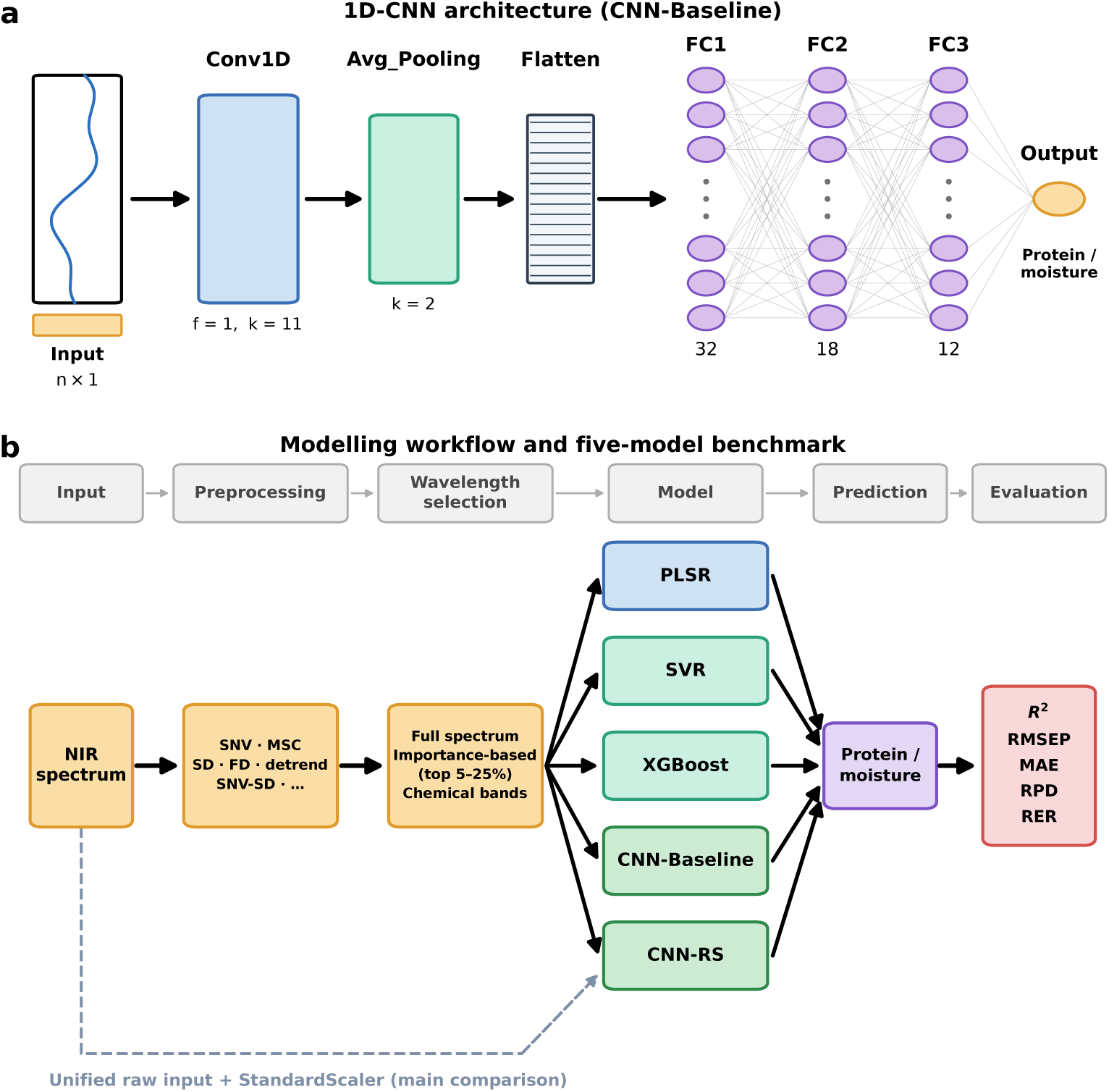
Overview of the modelling framework. **(a)** Final CNN-Baseline architecture. NIRS5000 and XDS inputs contained 700 and 1050 wavelengths, respectively. A single one-filter convolutional layer with a kernel size of 11 was followed by average pooling, flattening, three fully connected layers (32, 18, and 12 units), and one regression output. Neurons are shown schematically; complete specifications are provided in Table 2. **(b)** Benchmark workflow: all five models are evaluated under three input conditions, a unified raw input (dashed line), each model’s optimal preprocessing over the full spectrum, and this prerprocessing plus the PLSR selected wavelength subset shared by SVR, XGBoost and both CNNs, and evaluated using *R*^2^, RMSEP, MAE, RPD, and RER.

**Table 2:**
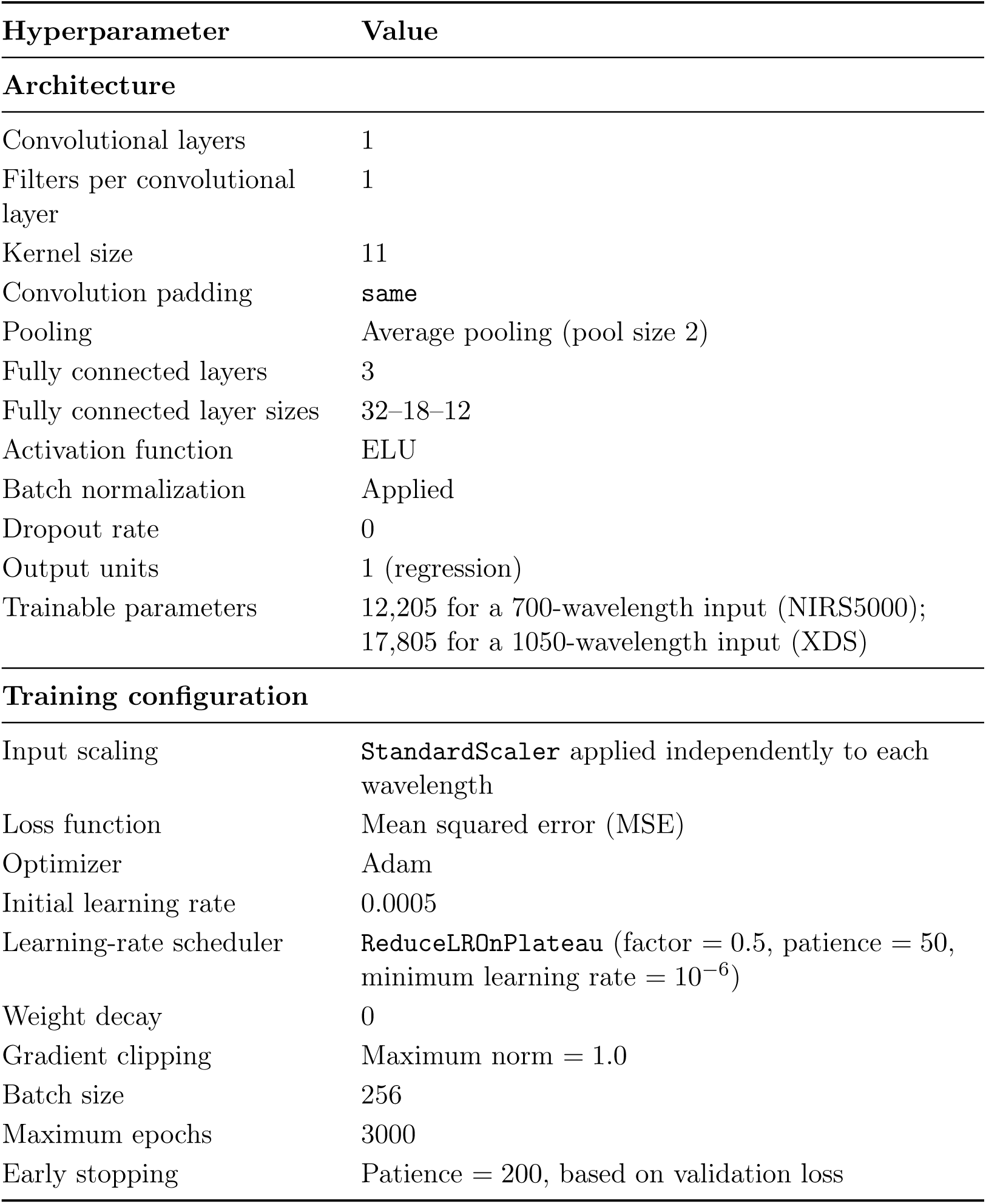
Final architecture and training configuration of the optimized one-dimensional CNN.

To complement the manually designed CNN-Baseline, the other 1D-CNN was obtained through automatic random grid search for optimal parameters and was named CNN-RS. Its purpose was twofold. First, it can serve as a benchmark to validate the simplicity-driven CNN-Baseline. If an extensive automated search across the same hyperparameter space failed to outperform the manually designed architecture, this provided strong evidence that additional architectural capacity was unnecessary. Second, from the practical standpoint, search-based models eliminated the need for repetitive manual grid searches whenever the datasets changed and can more easily be transferred across different grain types, compositions, and instruments.

CNN-RS was constructed by conducting 300 independent random trials on a thirteen-factor space. The configuration for each trial included the number of convolutional layers {1, 2, 3}, the number of filters per layer {1, 2, …, 32} for the first convolutional layer and {1, 2, …, 128} for subsequent layers, the first-layer kernel size {5, 7, 11, 15, 21, 31} and kernel sizes in subsequent layers {3, 4, …, 11}, pooling type (average or max) with pool size 2, the number and width of fully connected layers {1, 2, 3, 4} with each layer containing between 4 and 256 units, activation functions (ReLU, ELU, Leaky ReLU, or SELU), use of batch normalization, dropout rate [0, 0.5], weight decay [0, 0.02], learning rate [5 × 10^−5^, 1 × 10^−2^], and batch size {32, 64, 128, 256}.

The joint random search must cover a space within a fixed budget of 300 trials, so the range of each factor was concentrated around the region identified as the optimal ablation. Configurations were sampled with a fixed NumPy pseudorandom generator to ensure that the same 300 architectures were examined for each dataset. Each candidate model was trained under the protocol of Section 2.8 and ranked on their validation-set *R*^2^. The configuration with the best performance was selected as CNN-RS. CNN-Baseline and CNN-RS were then trained and evaluated on all six datasets with 25 seeds each. The overall workflow diagram for five models (PLSR, SVR, XGBoost, CNN-Baseline, CNN-RS) were shown in Fig. 2b.

#### 2.6.3. Input variants for the fully optimized comparison

To determine whether convolutional networks benefit from the preprocessing and wavelength selection applied to chemometrics and machine learning models, each dataset was subjected to additional analysis using three input variants that differed only in terms of the input variables received by the network.

##### Raw

This input had not underwent spectral preprocessing. It utilized the full spectrum and was scaled only along the column axis. This configuration was also used in the main comparison.

##### Preprocessed

This input underwent spectral preprocessing and utilized the full spectrum with column-wise scaling.

##### Fully optimized

The input based on the optimal preprocessing strategy and was limited to the subset of wavelengths derived from PLSR.

An optimal preprocessing strategy was selected for the CNN independently of the PLSR selection. Twelve strategies from Section 2.3.1 were screened across three seeds (42–44) by reducing the epoch budget using a fixed CNN-Baseline architecture and the strategy with the highest average validation-set *R*^2^ was retained. Both CNN-Baseline and CNN-RS were evaluated across all three variants, and the architecture search described in Section 2.6.2 was independently repeated for each variant using the same random seed, so that all three variants examined the same sequence of 300 candidate architectures, differing only in their inputs. The CNN-Baseline architecture was tuned on the full-length spectra, and a kernel of width 11 spanned a very different fraction of a 175-wavelength subset than a 700-wavelength spectrum. Therefore, variants evaluated with an unadapted architecture would be penalized for reasons unrelated to their input.

Since each architecture was evaluated under each variant, each pair of variants resulted in up to 300 comparisons of validation-set *R*^2^. These findings were summarized using proportions of architectures biased toward a particular variant and the Wilcoxon signed-rank test. The Wilcoxon signed-rank test was preferred to the paired *t*-test. Because the data diverged toward larger negative *R*^2^ values in some sampling configurations, which increased the variance of the differences and compromises the ability of parametric tests to detect differences.

### 2.7. Performance metrics

The predictive performance, accuracy, and robustness of the calibrated model were rigorously quantified on independent validation and test sets using five complementary statistical metrics.

Let *y_i_* denote the reference (measured) value of the constituent for sample *i*, *ŷ_i_* denote the corresponding model-predicted value, *y̅* represent the arithmetic mean of the reference values, and *N* represent the total number of samples evaluated in the dataset. The performance metrics are mathematically defined as follows:

Coefficient of Determination (*R*^2^): Measure the proportion of variance in the dependent variable that can be predicted by the independent variable.

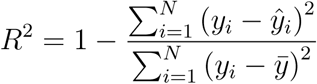

Root Mean Square Error of Prediction (RMSEP): The absolute root means square deviation between the quantitative forecast and the reference value.

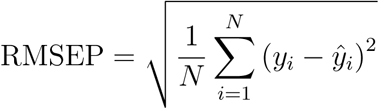

Mean Absolute Error (MAE): It represents the mean absolute value of prediction errors and provides a linear scoring metric in which all individual variations are weighted equally.

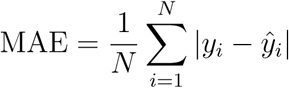

Ratio of Performance to Deviation (RPD): It evaluates the analytical utility of the calibration model by examining the inherent variability in the target population using standardized RMSEP.

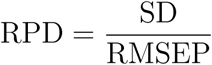

where SD is the standard deviation of the reference values, computed as:

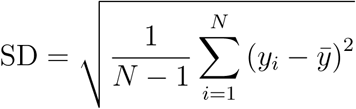

Range Error Ratio (RER): It provides an alternative normalized index of model’s performance by evaluating the prediction error relative to the total span of the constituent values.

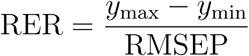

where *y*_max_ and *y*_min_ represent the maximum and minimum measured values within the reference dataset, respectively.

Higher *R*^2^, RPD and RER values and lower RMSEP and MAE values indicate better predictive performance (Fearn, 2002; Smyth et al., 2008; Williams et al., 2019).

### 2.8. Model training, evaluation and implementation

All data operations, model architectures, and statistical evaluations were implemented in Python 3.12.3. PLSR, SVR and cross-validated grid search used scikit-learn 1.7.0; XGBoost models used the XGBoost library 3.0.2; the 1D-CNN were implemented in PyTorch 2.7.0 with CUDA 12.8 acceleration (Paszke et al., 2019); statistical testing used SciPy 1.15.3. Computation was performed under Ubuntu (WSL 2) on a workstation equipped with an Intel Core i7-10700K CPU (3.80 GHz), 48 GB RAM, and an NVIDIA GeForce RTX 5090 GPU (32 GB), which accelerated the training of the CNNs and XGBoost models.

Both networks were trained with the Adam optimizer to minimize the mean squared error (MSE) loss for up to 3000 epochs. For CNN-Baseline, the initial learning rate and batch size were fixed at the ablation-selected values in Table 2; for CNN-RS, they were part of the searched configuration and varied by dataset (Table S7). The early-stopping rule, learning-rate scheduler and gradient clipping described below were common to both networks. To prevent overfitting and ensure optimal convergence, an early stopping mechanism based on validation loss was implemented with a tolerance threshold of 200 epochs. Additionally, a ReduceLROnPlateau scheduler decayed the learning rate during training, and gradient norm clipped at 1.0 to prevent gradient explosion. The network weights from the epoch with minimum validation loss were retained by deep copying each parameter tensor at that epoch and restored before evaluation.

Since the weights for network training were initialized randomly, each architecture was trained 25 times with 42-66 seeds, while the data partition remained fixed. This isolated sensitivity to initialization rather than sensitivity to partitioning, which remained unchanged so that all models could be compared using the same data. PLSR ran only once because of deterministic mathematical nature. The results were reported as the mean *±* standard deviation of 25 runs. For the comparative analysis requiring a single deterministic run (such as plotting a scatter plot of predictions versus measurements), a representative seed was selected for each deep learning model. The representative run was chosen by identifying the specific seed with the smallest cumulative absolute deviation between individual performance metrics and the macro-average metrics calculated across 25 runs. Differences between models and among input variables were assessed using paired t-tests on the test-set *R*^2^ values of 25 seeds, paired on seeds. Effects estimated across the architecture search were assessed with the Wilcoxon signed rank test, as described in Section 2.6.3.

The magnitude of CNN-RS’s advantage over PLSR varies with the number of training samples. The test *R*^2^ values ranged from 0.022 to 0.041 when the samples were larger than 985, while it narrowed to 0.009 on the maize dataset (n=585), and a reversal occurred only on the wheat protein dataset (XDS, n=500). The mean value of PLSR (0.9114) was approximately one seed-level standard deviation higher than that of CNN-RS (0.9020 ± 0.0089). However, the XDS dataset differed from the other five datasets not only in terms of sample size, which was the only dataset derived from a second spectrometer with a wavelength range of 400-2498 nm rather than 1100-2498 nm and it had the narrowest composition range. These factors interacted with each other, so the reversal in results could not be attributed solely to sample size. It was worth noting that the CNN-RS model still outperformed the others on the smallest NIRS5000 dataset, which measured on maize moisture. SVR offered a good balance of accuracy and reproducibility, performing close to the best models on several datasets while showing essentially no run-to-run variability, and XGBoost performed the worst on every dataset (Section 3.4).

## 3. Results and Discussion

### 3.1. Overall model comparison

A comprehensive comparison of all five model across six datasets was presented in Table 3 and visualized as heatmaps in Figure 3. The best model varied depending on the dataset. CNN-RS achieved the highest test-set *R*^2^ on the five of the six datasets, while PLSR model performed best on the smallest protein dataset, wheat protein (XDS). In every dataset, the best model reached a test *R*^2^ of 0.82-0.98 and an RPD greater than 3.3 on five datasets, while it was 2.4 on maize moisture. It indicated that the calibration was sufficient for screening but short of quantitative use.

**Figure 3:**
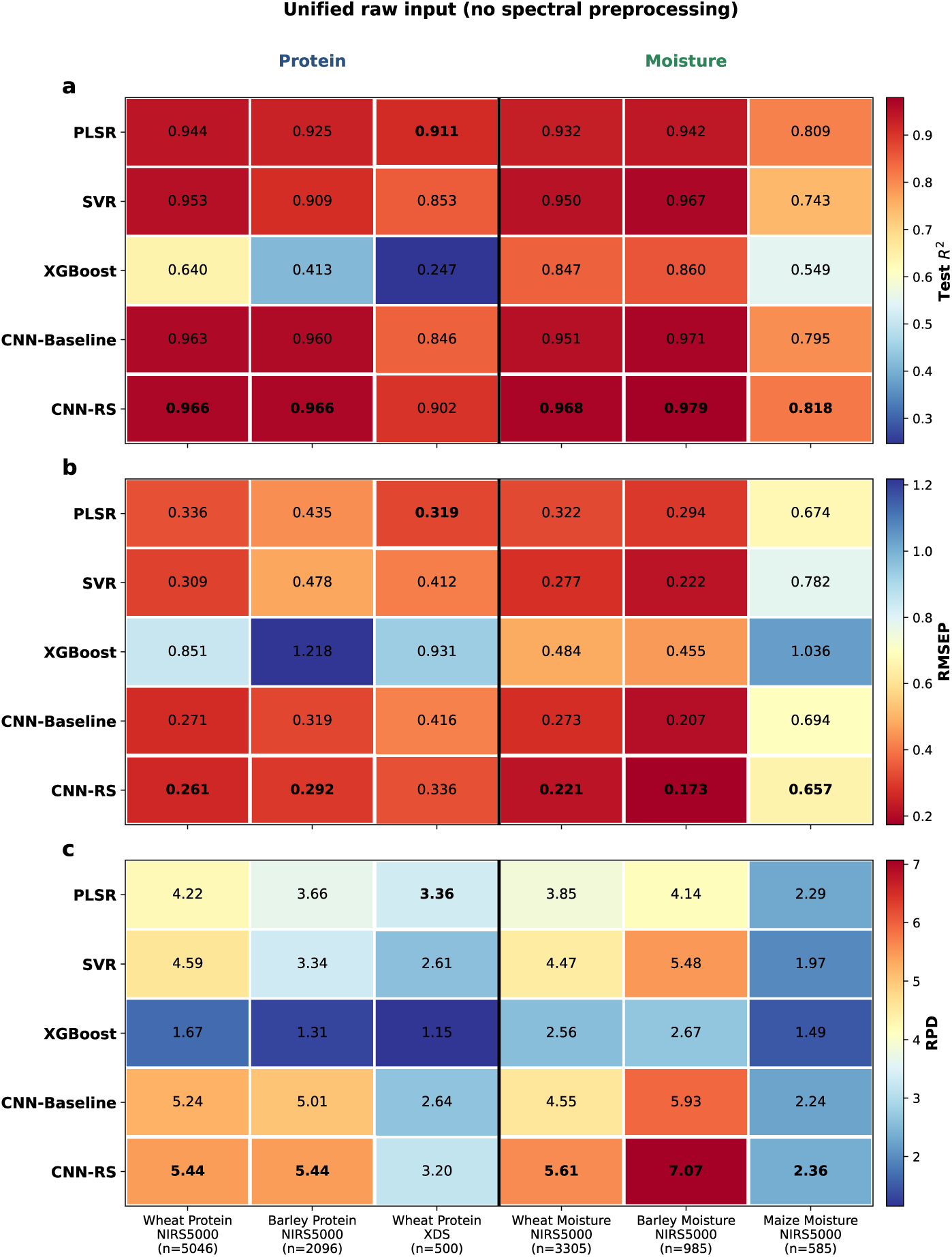
Heatmap summary of test-set performance for five models across six datasets using the unified raw input, in which each model receives the spectra with column-wise scaling only: (a) coefficient of determination (*R*^2^), (b) root mean square error of prediction (RMSEP), and (c) ratio of performance to deviation (RPD). Rows are models and columns are datasets, with three protein datasets to the left and the three moisture datasets to the right of the vertical divider. Each panel has its own color scale, oriented so that red always indicates better performance, and the best-performing model of the value is displayed in bold. Values are means over 25 seeds (42-66) for SVR, XGBoost, CNN-Baseline, CNN-RS and a single deterministic run of PLSR.

**Table 3:**
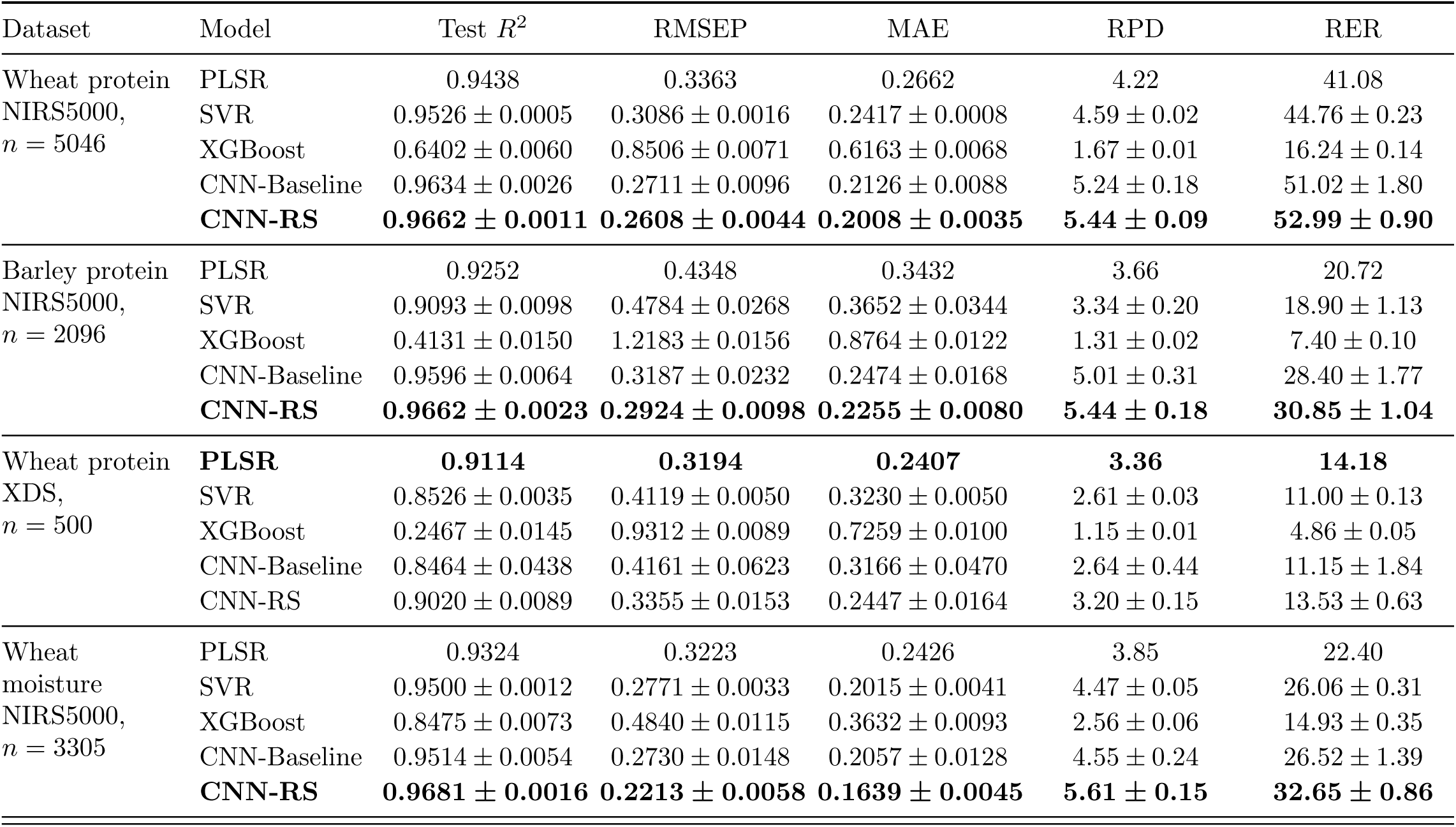
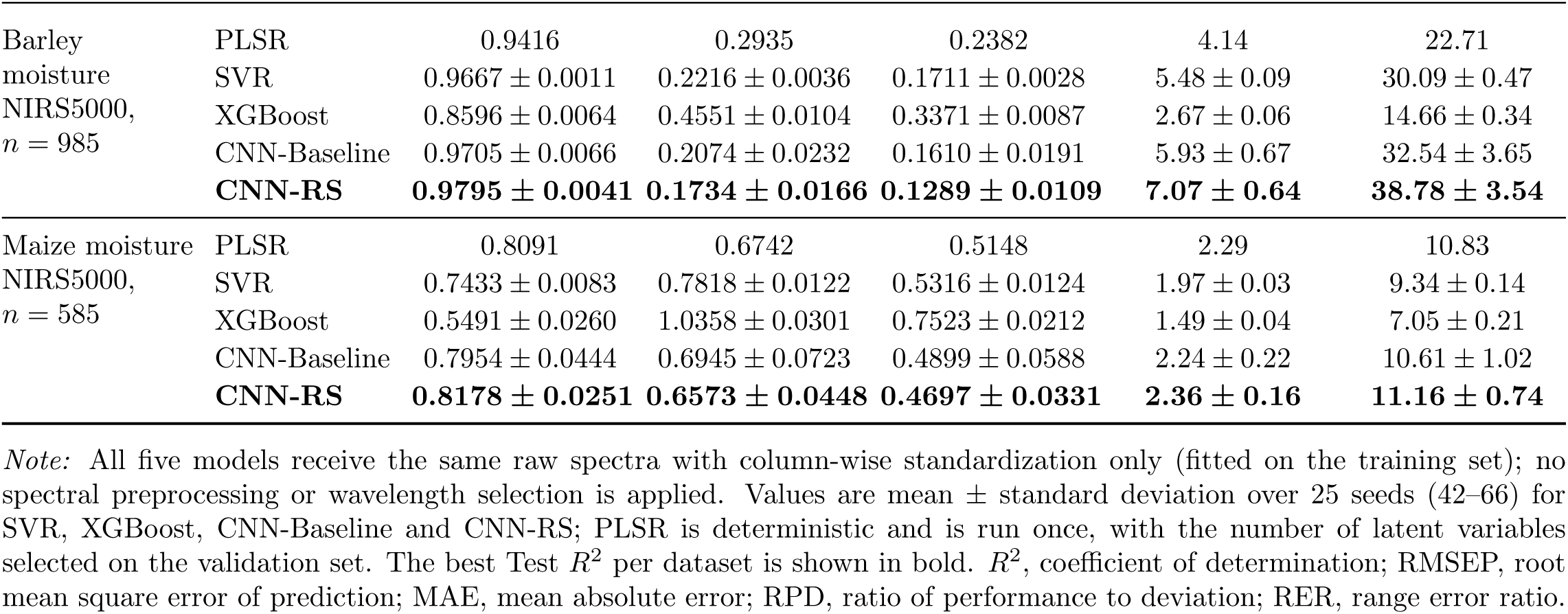
Test-set performance of all models across the six datasets using the unified raw input.

In addition to sample size, two other mechanisms may have contributed to this reversal. First, the additional spectral range contained almost no useful signal: the validation-based PLSR selection did not retain any channels below 680 nm, and of the 350 channels below 1100 nm, only 55 were retained, 45 of which were concentrated in the water band at 970 nm and the N-H second overtone near 1020 nm. PLS projections are designed to suppress variance unrelated to the response, whereas the CNN had to learn to reduce the weights of uninformative channels based on only 0.33 training samples per wavelength (compared to 0.58 for maize moisture). Second, the narrow reference range mathematically lowers the *R*^2^ values: when the RMSEP values of the raw inputs approached the levels observed for wheat protein on the NIRS5000 (0.3194 vs. 0.3363), the *R*^2^ values for all models are inevitably lower. Under all three input conditions, PLSR lead over both CNNs persisted under all three input conditions (0.008-0.013 in *R*^2^ relative to CNN-RS), indicating that a mere data-volume effect, in line with the conditional view of CNN benefits (Passos, 2026).

These rankings applied to the unified raw input, chosen so that differences would reflect the models rather than the preprocessing each happened to receive. Tables 4 and 5, as well as Figures 4 and 5, reported the same datasets after applying optimal preprocessing to every model and after additionally applying wavelength selection. The model’s performance varied significantly. With preprocessing alone, the best model was SVR on three datasets, CNN-RS on two and PLSR on one. Once wavelength selection was also incorporated, SVR was the most accurate on four of the six datasets. Therefore, the advantage of the convolutional networks in Table 3 was their strong robustness to unprocessed input than as a higher attainable accuracy. These findings are useful because they answer different questions: which model to choose given a fixed preparation process, and which to choose given the freedom and expertise to optimize it.

**Figure 4:**
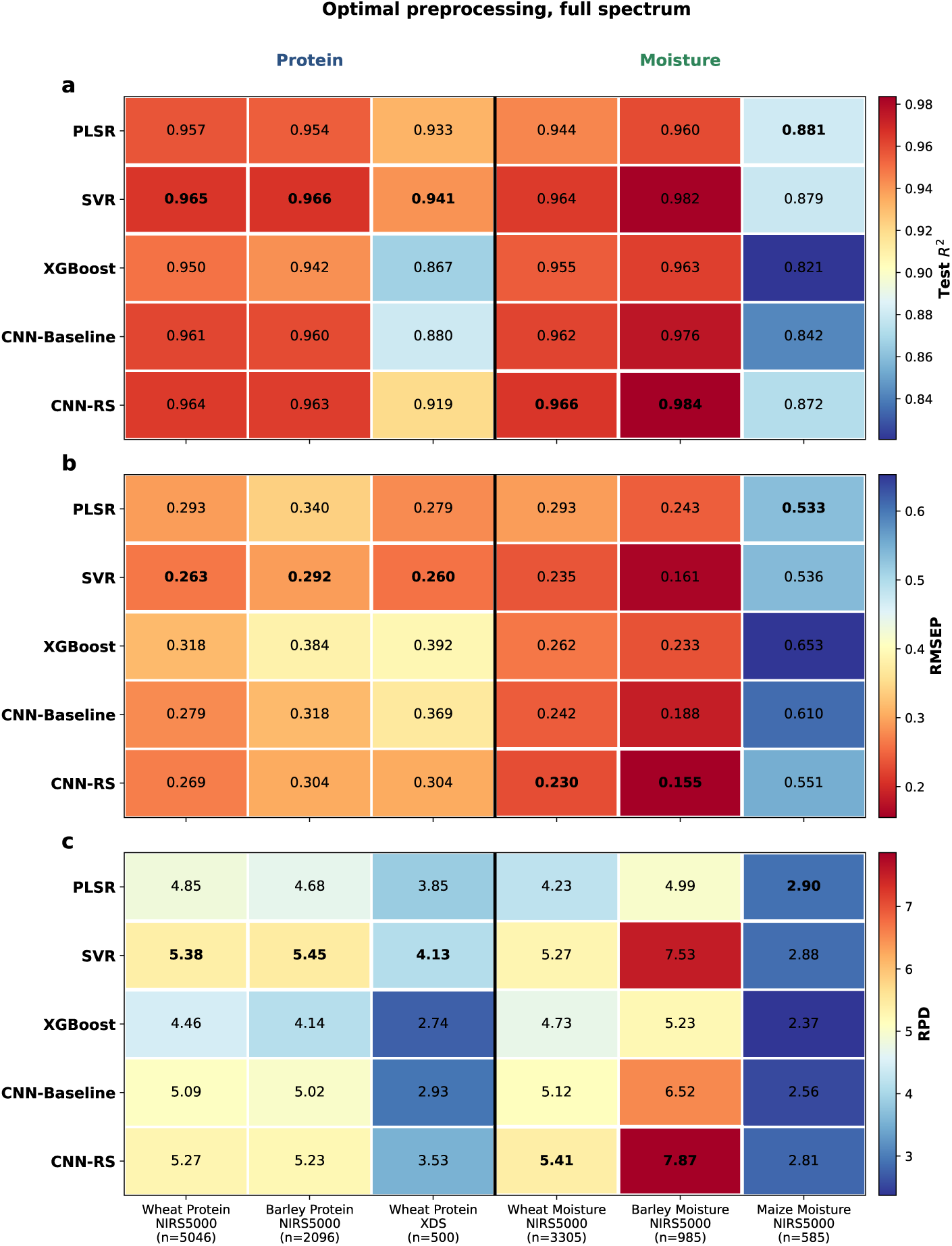
Heatmap summary of test-set performance for five models across six datasets with optimal spectral preprocessing applied to every model over the full spectrum: (a) coefficient of determination (*R*^2^), (b) root mean square error of prediction (RMSEP), and (c) ratio of performance to deviation (RPD). The preprocessing was selected independently for each model and dataset by ranking the same twelve strategies on the validation set; no wavelength selection is applied here. Layout, color scales, highlighting and the seed protocol are as in Figure 3.

**Figure 5:**
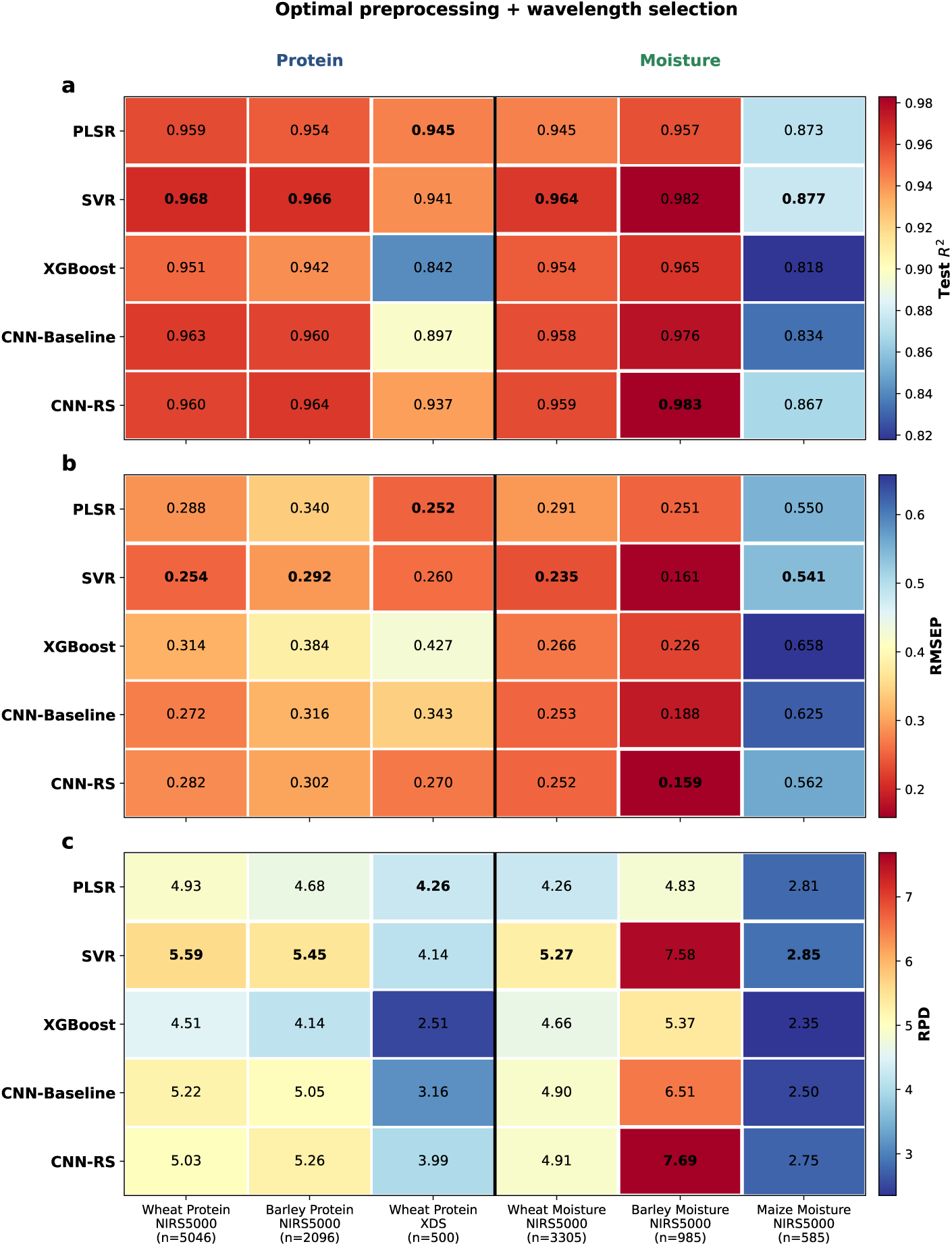
Heatmap summary of test-set performance for five models across six datasets with the preprocessing and wavelength selection applied to every model: (a) coefficient of determination (*R*^2^), (b) root mean square error of prediction (RMSEP), and (c) ratio of performance to deviation (RPD). Each model retained the preprocessing strategy selected on the validation set, while SVR, XGBoost, CNN-Baseline, and CNN-RS used the wave-length subset selected by PLSR on the validation set of the same dataset, so the effect of wavelength selection was measured on the same subset across these four models; PLSR used its retained configuration. Layout, color scales, highlighting and the seed protocol are as in Figure 3.

**Table 4:**
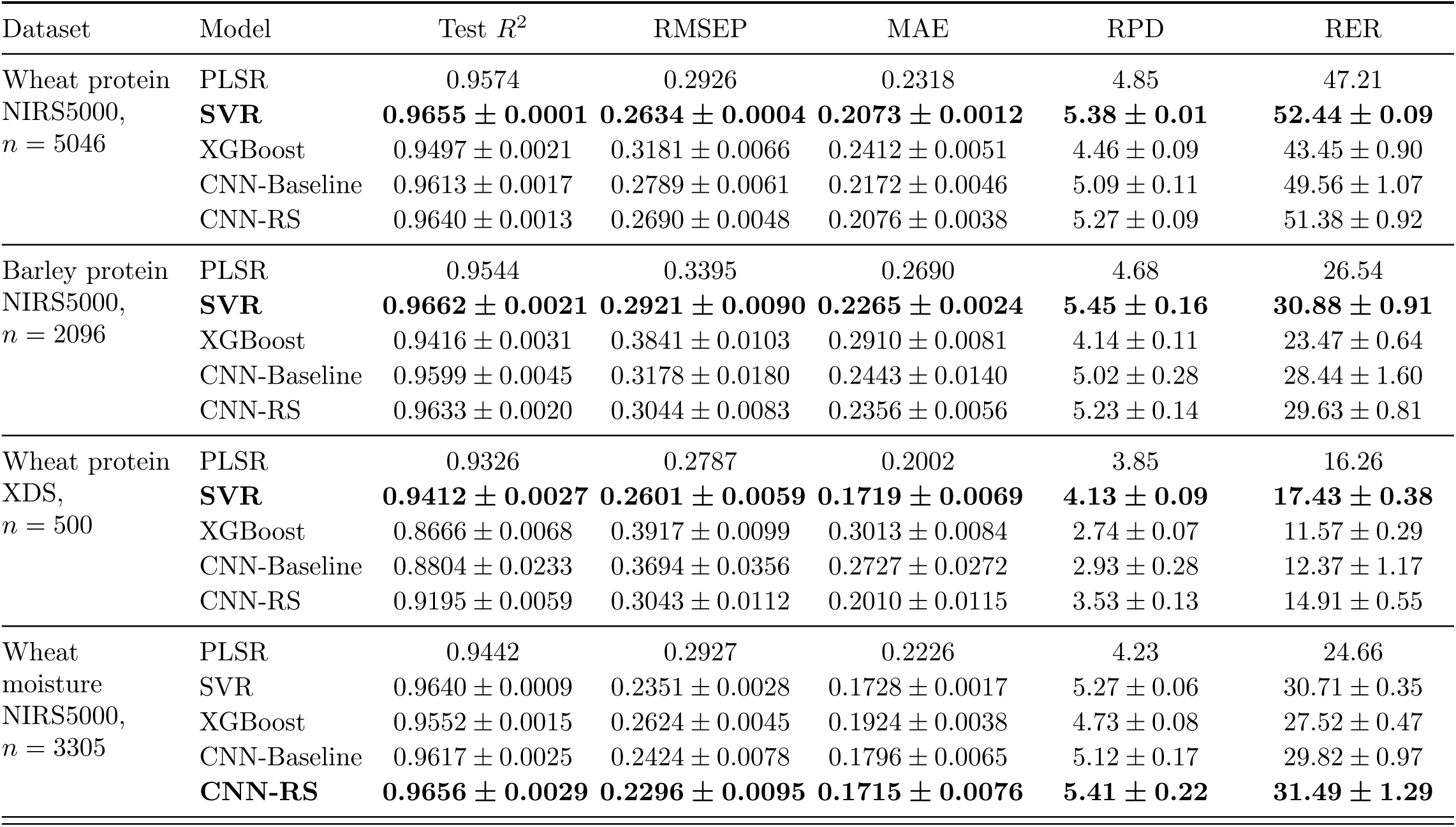
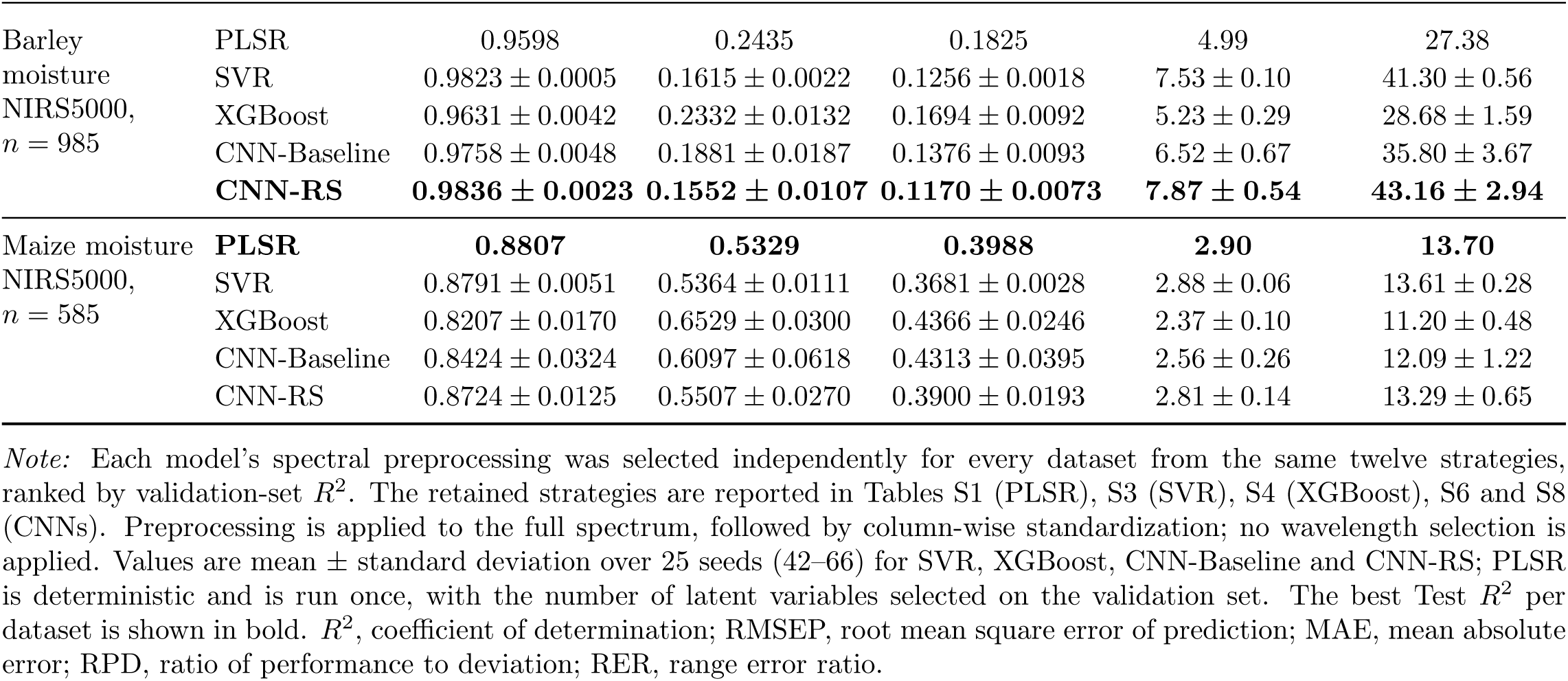
Test-set performance of all models across the six datasets with spectral preprocessing applied to every model.

| Dataset | Model | Test $R^2$ | RMSEP | MAE | RPD | RER |
| --- | --- | --- | --- | --- | --- | --- |
| Wheat protein<br>NIRS5000,<br>$n = 5046$ | PLSR | 0.9574 | 0.2926 | 0.2318 | 4.85 | 47.21 |
|  | **SVR** | **0.9655 ± 0.0001** | **0.2634 ± 0.0004** | **0.2073 ± 0.0012** | **5.38 ± 0.01** | **52.44 ± 0.09** |
| | XGBoost | 0.9497 ± 0.0021 | 0.3181 ± 0.0066 | 0.2412 ± 0.0051 | 4.46 ± 0.09 | 43.45 ± 0.90 |
| | CNN-Baseline | 0.9613 ± 0.0017 | 0.2789 ± 0.0061 | 0.2172 ± 0.0046 | 5.09 ± 0.11 | 49.56 ± 1.07 |
| | CNN-RS | 0.9640 ± 0.0013 | 0.2690 ± 0.0048 | 0.2076 ± 0.0038 | 5.27 ± 0.09 | 51.38 ± 0.92 |
| Barley protein<br>NIRS5000,<br>$n = 2096$ | PLSR | 0.9544 | 0.3395 | 0.2690 | 4.68 | 26.54 |
|  | **SVR** | **0.9662 ± 0.0021** | **0.2921 ± 0.0090** | **0.2265 ± 0.0024** | **5.45 ± 0.16** | **30.88 ± 0.91** |
| | XGBoost | 0.9416 ± 0.0031 | 0.3841 ± 0.0103 | 0.2910 ± 0.0081 | 4.14 ± 0.11 | 23.47 ± 0.64 |
| | CNN-Baseline | 0.9599 ± 0.0045 | 0.3178 ± 0.0180 | 0.2443 ± 0.0140 | 5.02 ± 0.28 | 28.44 ± 1.60 |
| | CNN-RS | 0.9633 ± 0.0020 | 0.3044 ± 0.0083 | 0.2356 ± 0.0056 | 5.23 ± 0.14 | 29.63 ± 0.81 |
| Wheat protein<br>XDS,<br>$n = 500$ | PLSR | 0.9326 | 0.2787 | 0.2002 | 3.85 | 16.26 |
|  | **SVR** | **0.9412 ± 0.0027** | **0.2601 ± 0.0059** | **0.1719 ± 0.0069** | **4.13 ± 0.09** | **17.43 ± 0.38** |
| | XGBoost | 0.8666 ± 0.0068 | 0.3917 ± 0.0099 | 0.3013 ± 0.0084 | 2.74 ± 0.07 | 11.57 ± 0.29 |
| | CNN-Baseline | 0.8804 ± 0.0233 | 0.3694 ± 0.0356 | 0.2727 ± 0.0272 | 2.93 ± 0.28 | 12.37 ± 1.17 |
| | CNN-RS | 0.9195 ± 0.0059 | 0.3043 ± 0.0112 | 0.2010 ± 0.0115 | 3.53 ± 0.13 | 14.91 ± 0.55 |
| Wheat<br>moisture<br>NIRS5000,<br>$n = 3305$ | PLSR | 0.9442 | 0.2927 | 0.2226 | 4.23 | 24.66 |
| | SVR | 0.9640 ± 0.0009 | 0.2351 ± 0.0028 | 0.1728 ± 0.0017 | 5.27 ± 0.06 | 30.71 ± 0.35 |
| | XGBoost | 0.9552 ± 0.0015 | 0.2624 ± 0.0045 | 0.1924 ± 0.0038 | 4.73 ± 0.08 | 27.52 ± 0.47 |
| | CNN-Baseline | 0.9617 ± 0.0025 | 0.2424 ± 0.0078 | 0.1796 ± 0.0065 | 5.12 ± 0.17 | 29.82 ± 0.97 |
|  | **CNN-RS** | **0.9656 ± 0.0029** | **0.2296 ± 0.0095** | **0.1715 ± 0.0076** | **5.41 ± 0.22** | **31.49 ± 1.29** |

Table 4 continued from the previous page
| Dataset | Model | Test $R^2$ | RMSEP | MAE | RPD | RER |
| --- | --- | --- | --- | --- | --- | --- |
| Barley<br>moisture<br>NIRS5000,<br>$n = 985$ | PLSR | 0.9598 | 0.2435 | 0.1825 | 4.99 | 27.38 |
| | SVR | $0.9823 \pm 0.0005$ | $0.1615 \pm 0.0022$ | $0.1256 \pm 0.0018$ | $7.53 \pm 0.10$ | $41.30 \pm 0.56$ |
| | XGBoost | $0.9631 \pm 0.0042$ | $0.2332 \pm 0.0132$ | $0.1694 \pm 0.0092$ | $5.23 \pm 0.29$ | $28.68 \pm 1.59$ |
| | CNN-Baseline | $0.9758 \pm 0.0048$ | $0.1881 \pm 0.0187$ | $0.1376 \pm 0.0093$ | $6.52 \pm 0.67$ | $35.80 \pm 3.67$ |
|  | **CNN-RS** | **$0.9836 \pm 0.0023$** | **$0.1552 \pm 0.0107$** | **$0.1170 \pm 0.0073$** | **$7.87 \pm 0.54$** | **$43.16 \pm 2.94$** |
| Maize moisture<br>NIRS5000,<br>$n = 585$ | **PLSR** | **0.8807** | **0.5329** | **0.3988** | **2.90** | **13.70** |
| | SVR | $0.8791 \pm 0.0051$ | $0.5364 \pm 0.0111$ | $0.3681 \pm 0.0028$ | $2.88 \pm 0.06$ | $13.61 \pm 0.28$ |
| | XGBoost | $0.8207 \pm 0.0170$ | $0.6529 \pm 0.0300$ | $0.4366 \pm 0.0246$ | $2.37 \pm 0.10$ | $11.20 \pm 0.48$ |
| | CNN-Baseline | $0.8424 \pm 0.0324$ | $0.6097 \pm 0.0618$ | $0.4313 \pm 0.0395$ | $2.56 \pm 0.26$ | $12.09 \pm 1.22$ |
| | CNN-RS | $0.8724 \pm 0.0125$ | $0.5507 \pm 0.0270$ | $0.3900 \pm 0.0193$ | $2.81 \pm 0.14$ | $13.29 \pm 0.65$ |

*Note:* Each model's spectral preprocessing was selected independently for every dataset from the same twelve strategies, ranked by validation-set $R^2$ . The retained strategies are reported in Tables S1 (PLSR), S3 (SVR), S4 (XGBoost), S6 and S8 (CNNs). Preprocessing is applied to the full spectrum, followed by column-wise standardization; no wavelength selection is applied. Values are mean ± standard deviation over 25 seeds (42–66) for SVR, XGBoost, CNN-Baseline and CNN-RS; PLSR is deterministic and is run once, with the number of latent variables selected on the validation set. The best Test $R^2$ per dataset is shown in bold. $R^2$ , coefficient of determination; RMSEP, root mean square error of prediction; MAE, mean absolute error; RPD, ratio of performance to deviation; RER, range error ratio.

**Table 5:** Test-set performance of all models across the six datasets with spectral preprocessing and wavelength selection applied to every model.

| Dataset | Model | Test $R^2$ | RMSEP | MAE | RPD | RER |
| --- | --- | --- | --- | --- | --- | --- |
| Wheat protein<br>NIRS5000,<br>$n = 5046$ | PLSR | 0.9589 | 0.2876 | 0.2288 | 4.93 | 48.03 |
|  | **SVR** | **0.9680 ± 0.0003** | **0.2538 ± 0.0010** | **0.1984 ± 0.0005** | **5.59 ± 0.02** | **54.44 ± 0.22** |
| | XGBoost | 0.9508 ± 0.0016 | 0.3144 ± 0.0052 | 0.2369 ± 0.0045 | 4.51 ± 0.07 | 43.95 ± 0.73 |
| | CNN-Baseline | 0.9632 ± 0.0017 | 0.2719 ± 0.0062 | 0.2093 ± 0.0050 | 5.22 ± 0.12 | 50.83 ± 1.14 |
| | CNN-RS | 0.9604 ± 0.0019 | 0.2822 ± 0.0068 | 0.2175 ± 0.0053 | 5.03 ± 0.12 | 48.98 ± 1.18 |
| Barley protein<br>NIRS5000,<br>$n = 2096$ | PLSR | 0.9544 | 0.3395 | 0.2690 | 4.68 | 26.54 |
|  | **SVR** | **0.9662 ± 0.0021** | **0.2921 ± 0.0090** | **0.2265 ± 0.0024** | **5.45 ± 0.16** | **30.88 ± 0.91** |
| | XGBoost | 0.9416 ± 0.0031 | 0.3841 ± 0.0103 | 0.2910 ± 0.0081 | 4.14 ± 0.11 | 23.47 ± 0.64 |
| | CNN-Baseline | 0.9604 ± 0.0041 | 0.3160 ± 0.0165 | 0.2428 ± 0.0119 | 5.05 ± 0.26 | 28.58 ± 1.49 |
| | CNN-RS | 0.9638 ± 0.0017 | 0.3024 ± 0.0072 | 0.2301 ± 0.0060 | 5.26 ± 0.13 | 29.81 ± 0.71 |
| Wheat protein<br>XDS,<br>$n = 500$ | **PLSR** | **0.9448** | **0.2521** | **0.1893** | **4.26** | **17.97** |
| | SVR | 0.9415 ± 0.0028 | 0.2595 ± 0.0063 | 0.1771 ± 0.0073 | 4.14 ± 0.11 | 17.46 ± 0.46 |
| | XGBoost | 0.8416 ± 0.0078 | 0.4269 ± 0.0105 | 0.3300 ± 0.0099 | 2.51 ± 0.06 | 10.62 ± 0.26 |
| | CNN-Baseline | 0.8972 ± 0.0183 | 0.3427 ± 0.0306 | 0.2501 ± 0.0247 | 3.16 ± 0.28 | 13.32 ± 1.20 |
| | CNN-RS | 0.9368 ± 0.0054 | 0.2695 ± 0.0115 | 0.1863 ± 0.0117 | 3.99 ± 0.17 | 16.84 ± 0.72 |
| Wheat<br>moisture<br>NIRS5000,<br>$n = 3305$ | PLSR | 0.9448 | 0.2913 | 0.2206 | 4.26 | 24.78 |
|  | **SVR** | **0.9640 ± 0.0007** | **0.2352 ± 0.0024** | **0.1728 ± 0.0032** | **5.27 ± 0.05** | **30.70 ± 0.31** |
| | XGBoost | 0.9539 ± 0.0014 | 0.2660 ± 0.0042 | 0.1945 ± 0.0034 | 4.66 ± 0.07 | 27.15 ± 0.43 |
| | CNN-Baseline | 0.9582 ± 0.0038 | 0.2533 ± 0.0115 | 0.1885 ± 0.0076 | 4.90 ± 0.22 | 28.56 ± 1.26 |
| | CNN-RS | 0.9585 ± 0.0017 | 0.2525 ± 0.0051 | 0.1876 ± 0.0050 | 4.91 ± 0.10 | 28.61 ± 0.59 |

Table 5 continued from the previous page
| Dataset | Model | Test $R^2$ | RMSEP | MAE | RPD | RER |
| --- | --- | --- | --- | --- | --- | --- |
| Barley<br>moisture<br>NIRS5000,<br>$n = 985$ | PLSR | 0.9572 | 0.2513 | 0.1913 | 4.83 | 26.53 |
| | SVR | 0.9825 ± 0.0017 | 0.1607 ± 0.0077 | 0.1245 ± 0.0034 | 7.58 ± 0.35 | 41.59 ± 1.93 |
| | XGBoost | 0.9653 ± 0.0024 | 0.2264 ± 0.0078 | 0.1667 ± 0.0069 | 5.37 ± 0.18 | 29.49 ± 0.99 |
| | CNN-Baseline | 0.9760 ± 0.0035 | 0.1878 ± 0.0137 | 0.1426 ± 0.0107 | 6.51 ± 0.49 | 35.70 ± 2.67 |
|  | **CNN-RS** | **0.9829 ± 0.0020** | **0.1586 ± 0.0094** | **0.1228 ± 0.0084** | **7.69 ± 0.45** | **42.18 ± 2.47** |
| Maize moisture<br>NIRS5000,<br>$n = 585$ | PLSR | 0.8731 | 0.5498 | 0.4222 | 2.81 | 13.28 |
|  | **SVR** | **0.8771 ± 0.0032** | **0.5410 ± 0.0070** | **0.3706 ± 0.0014** | **2.85 ± 0.04** | **13.50 ± 0.17** |
| | XGBoost | 0.8179 ± 0.0162 | 0.6580 ± 0.0286 | 0.4368 ± 0.0210 | 2.35 ± 0.10 | 11.11 ± 0.46 |
| | CNN-Baseline | 0.8338 ± 0.0389 | 0.6252 ± 0.0720 | 0.4438 ± 0.0486 | 2.50 ± 0.28 | 11.82 ± 1.34 |
| | CNN-RS | 0.8670 ± 0.0128 | 0.5621 ± 0.0273 | 0.3982 ± 0.0246 | 2.75 ± 0.14 | 13.02 ± 0.64 |

*Note:* Each model retains its corresponding preprocessing from Table 4, but is now limited to the wavelength subset selected via PLSR validation for this dataset (a single subset shared by all five models, selection strategies and subset size in Table S2). Spectral preprocessing is performed on the full spectrum prior to wavelength subsetting, followed by column-wise standardization after subsetting; both the hyperparameters and CNN architecture were re-optimized based on this input. Values are mean ± standard deviation over 25 seeds (42–66) for SVR, XGBoost, CNN-Baseline and CNN-RS; PLSR is deterministic and is run once, with the number of latent variables selected on the validation set. The best Test $R^2$ per dataset is shown in bold. $R^2$ , coefficient of determination; RMSEP, root mean square error of prediction; MAE, mean absolute error; RPD, ratio of performance to deviation; RER, range error ratio.

The preprocessing required for each model varies. The strategies selected independently for CNNs differed from those adopted by PLSR on any of the six datasets, and the direction was consistent: PLSR selected the second-order derivative strategy on five of six, while the screening process selected the first-order derivative strategy for CNNs on five of six. Yu et al. (2025) reported that the PLSR model selected first derivatives combined with multiplicative scatter correction, and ResNet-18 and Transformer model selected first derivatives and first derivatives combined with SNV, respectively. Deep learning models generally tended to adopt milder transformations. Two studies examining different particles, instruments, and architectures yielded identical ranking results, suggesting that this effect reflects characteristics of the model class rather than those specific datasets. Second-order derivative could resolve the overlapping absorption bands into clearer features. Their relationship with concentration was closer to linear, which a linear model required. Convolutional neural networks can extract the same information from smaller first-order derivatives, thereby preserving more signal and amplifying less noise. Therefore, preprocessing strategies optimized for one class of models should not be transferred to another class of models. Apart from preprocessing, wavelength selection had a very limited impact on the model performance. Under the validation-based selection, it retained the full spectra for one of the six datasets, barley protein. When a subset was retained as input, the further changes in test *R*^2^ relative to preprocessing alone ranged from *−*0.002 to +0.003 for SVR and from *−*0.009 to +0.017 for the CNNs. The only substantial gain occurred on wheat protein (XDS), so this impact was smaller than was often assumed.

From a practical standpoint, calibration-set size and composition jointly influence model choice. With several hundred samples, particularly when spectra contain regions carrying little constituent signal, optimized PLSR or SVR is preferable. With several thousand samples, 1D-CNNs can achieve competitive accuracy without preprocessing exploration while remaining more robust to unprocessed spectra.

### 3.2. Predicted versus measured relationships

Figure 6 and 7 showed a comparison of the predicted values from all five models with the measured values for the large dataset (wheat protein, NIRS5000) and the small dataset (maize moisture) with same raw input, and the remaining four datasets were shown in Figures S1-S4. Figures 8 and 9, with S5-S8, repeated the comparison after preprocessing, and Figures 10 and 11, with S9-S12, after preprocessing and wavelength selection. On the raw spectra, the fitted slope separated the models more sharply than *R*^2^ did. The average fitted slopes for PLSR, CNN-Baseline, CNN-RS and SVR across six datasets were 0.943, 0.937, 0.930, and 0.894, respectively. XGboost had a lower value of 0.579. This value dropped further to 0.295 on the wheat protein (XDS) dataset and 0.412 on the barley protein dataset. A slope of 0.295 indicated that a sample with one percentage point above the mean was predicted only 0.3 points above it. The model essentially stopped tracking the component. It is consistent with the test *R*^2^ of 0.247, although *R*^2^ alone did not make the failure mode visible.

**Figure 6:**
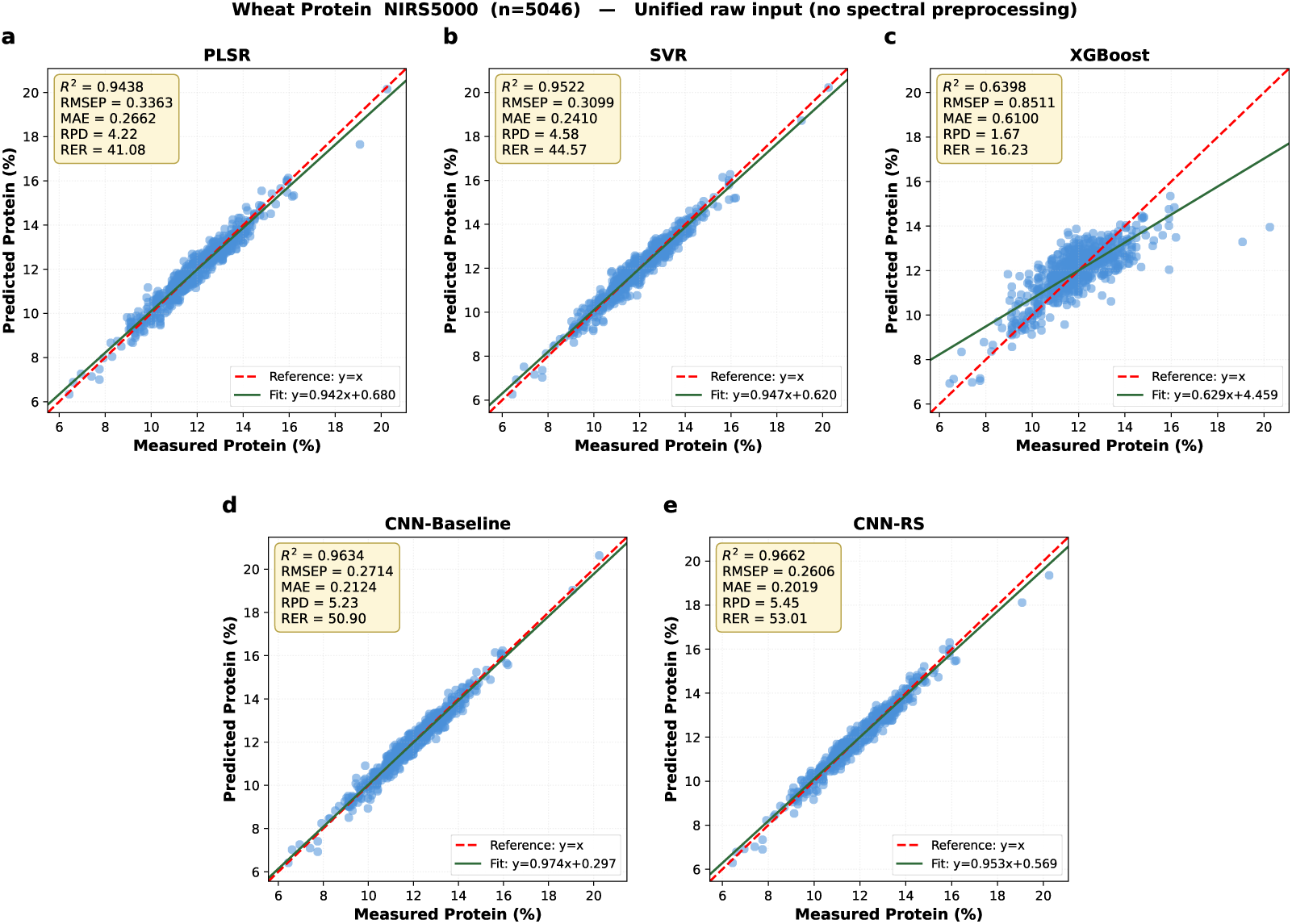
Measured versus predicted values of the five models on the largest dataset, wheat protein dataset (NIRS5000, *n* = 5046), under the unified raw input: (a) PLSR, (b) SVR, (c) XGBoost, (d) CNN-Baseline, and (e) CNN-RS. Test-set samples are shown as blue circles. The red dashed line is the reference (*y* = *x*) and the green solid line is the linear least squares fit (equation inset). The inset box reports the test-set *R*^2^, RMSEP, MAE, RPD, and RER. The CNNs, SVR and XGBoost panels show the representative seed, defined as the run, a representative seed is selected whose metrics is closest to the mean of the 25 seeds, and PLSR is deterministic.

**Figure 7:**
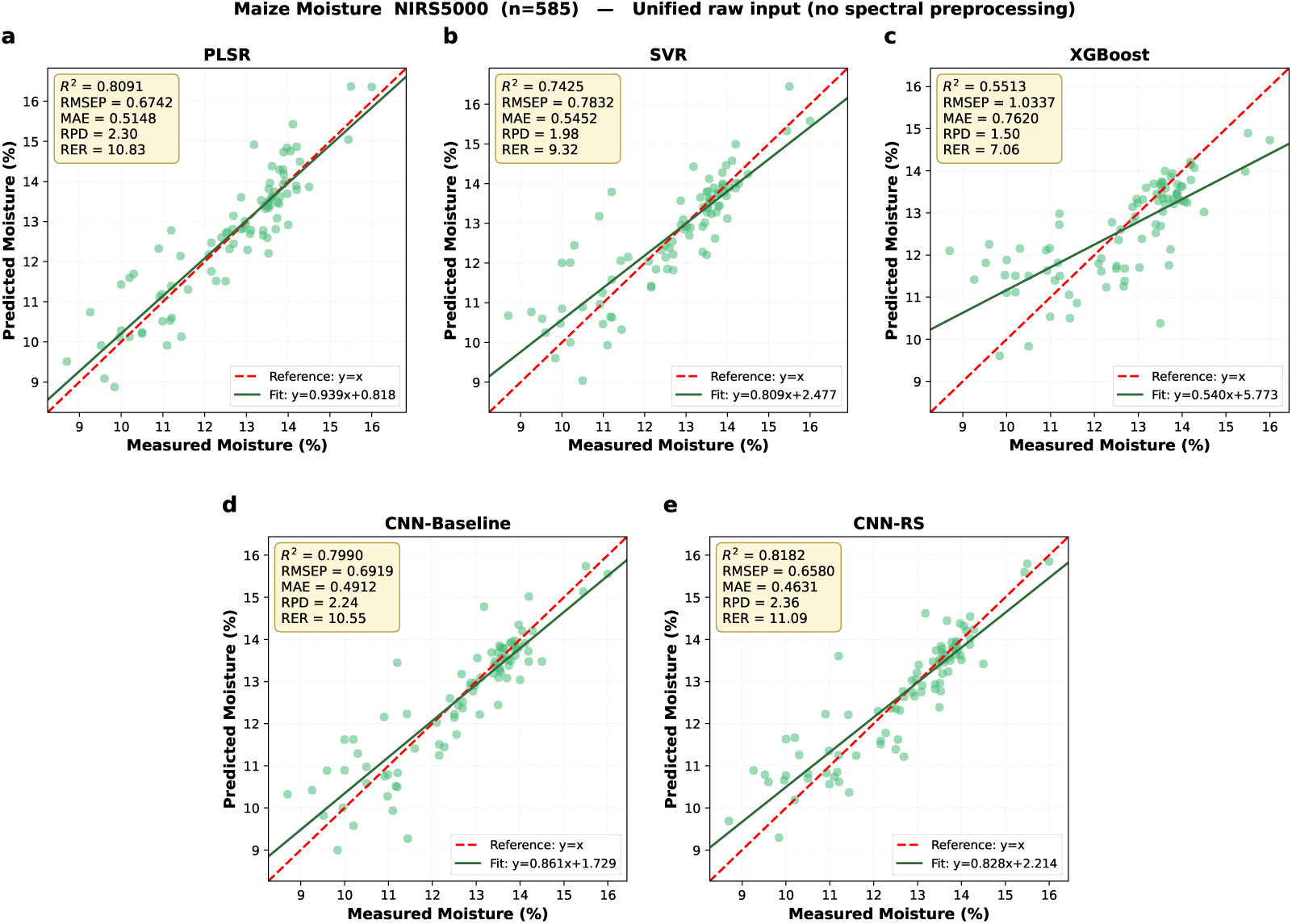
Measured versus predicted values of five models on the smallest dataset, maize moisture (NIRS5000, *n* = 585), under the unified raw input. Test-set samples are shown as green circles, and all other conventions, inset statistics and the choice of run are as in Figure 6.

**Figure 8:**
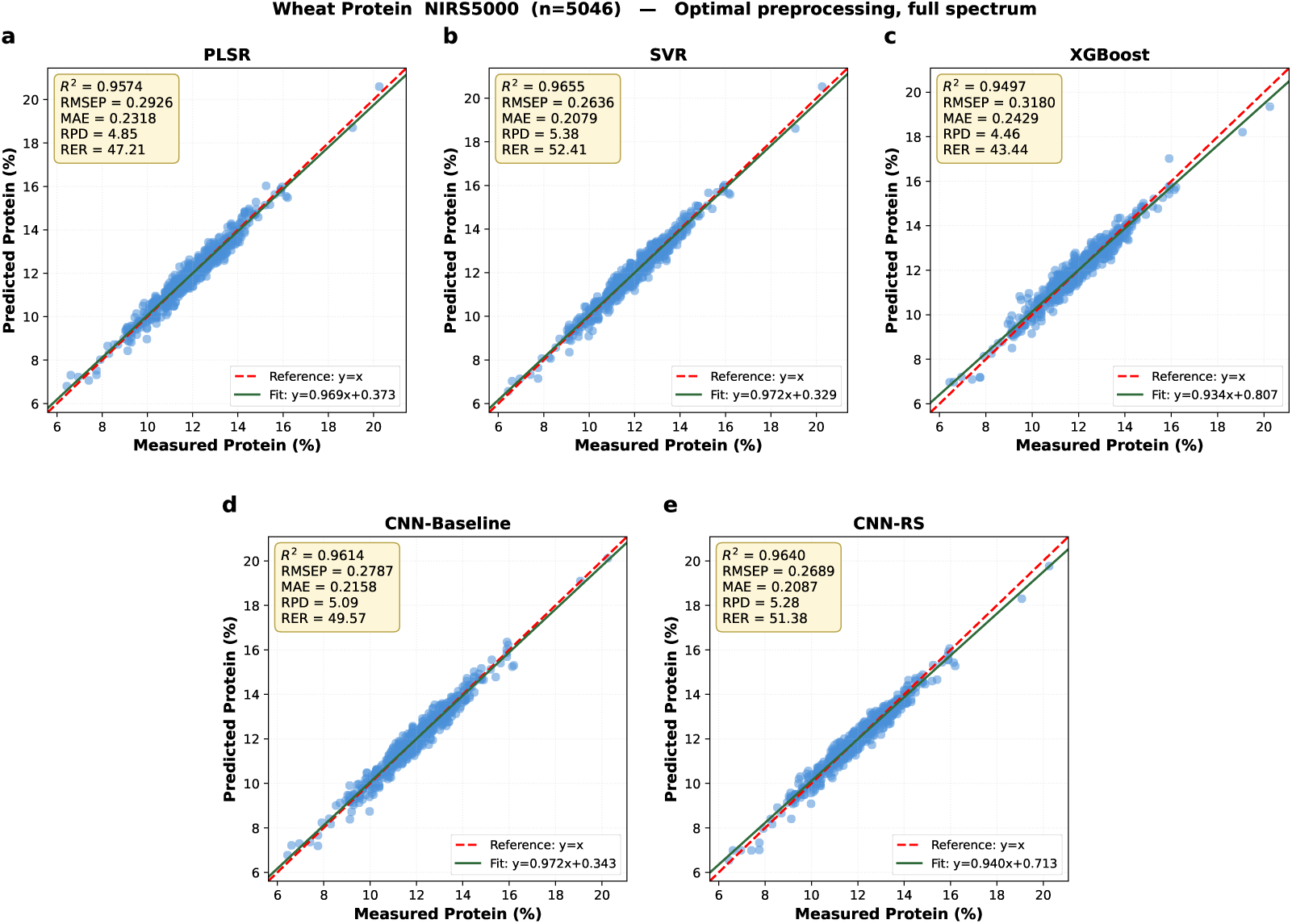
Measured versus predicted values on wheat protein (NIRS5000, *n* = 5046) after applying optimal preprocessing methods for each model on the validation set to the full spectra. SVR, XGBoost, CNN-Baseline and CNN-RS are shown at their representative seeds and PLSR is deterministic; all other conventions are as in Figure 6.

**Figure 9:**
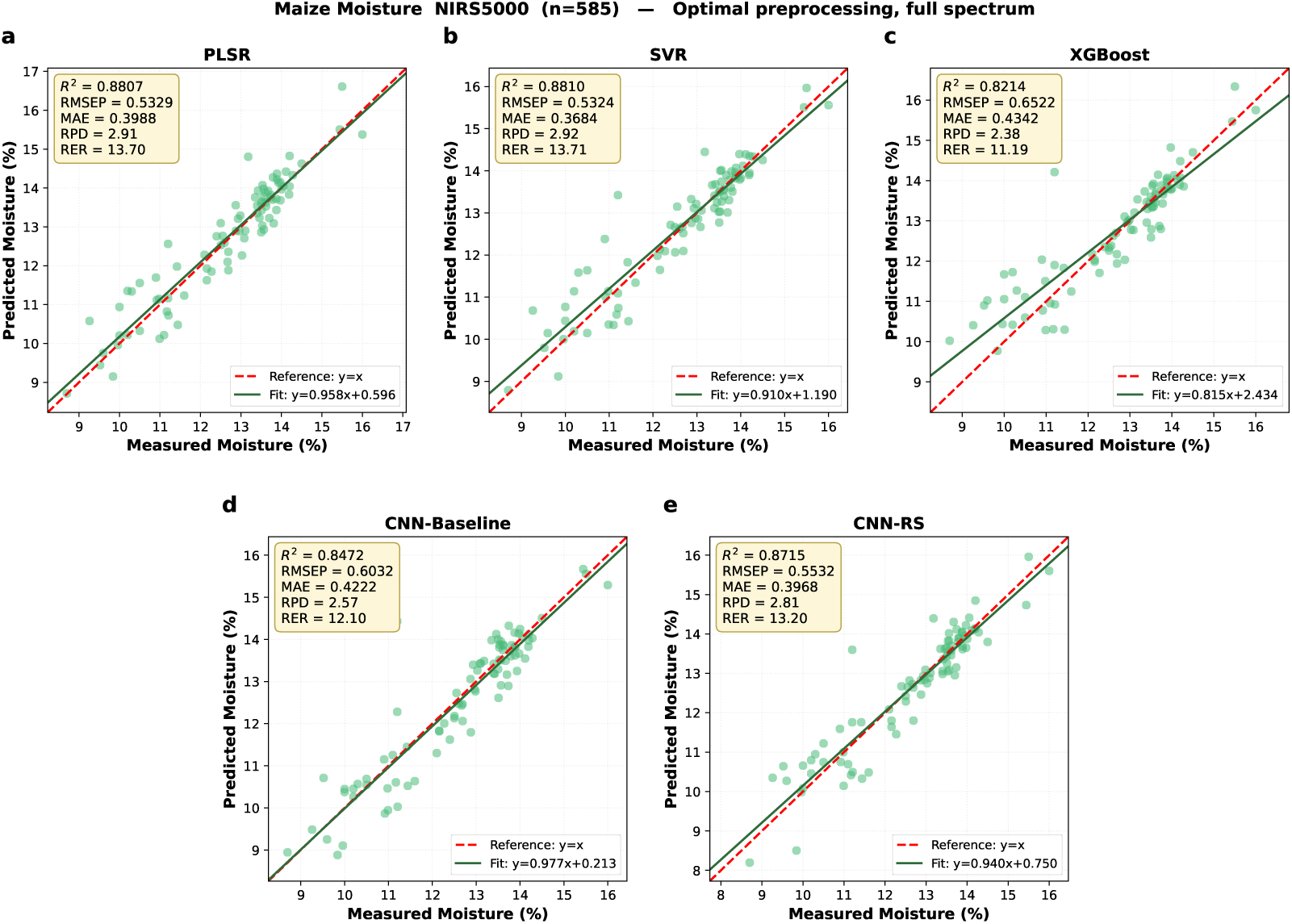
Measured versus predicted values on maize moisture (NIRS5000, *n* = 585) after applying optimal preprocessing methods for each model on the validation set to the full spectra. Test-set samples are shown in green; SVR, XGBoost and both CNNs are shown at their representative seeds, and all other conventions are as in Figure 6.

**Figure 10:**
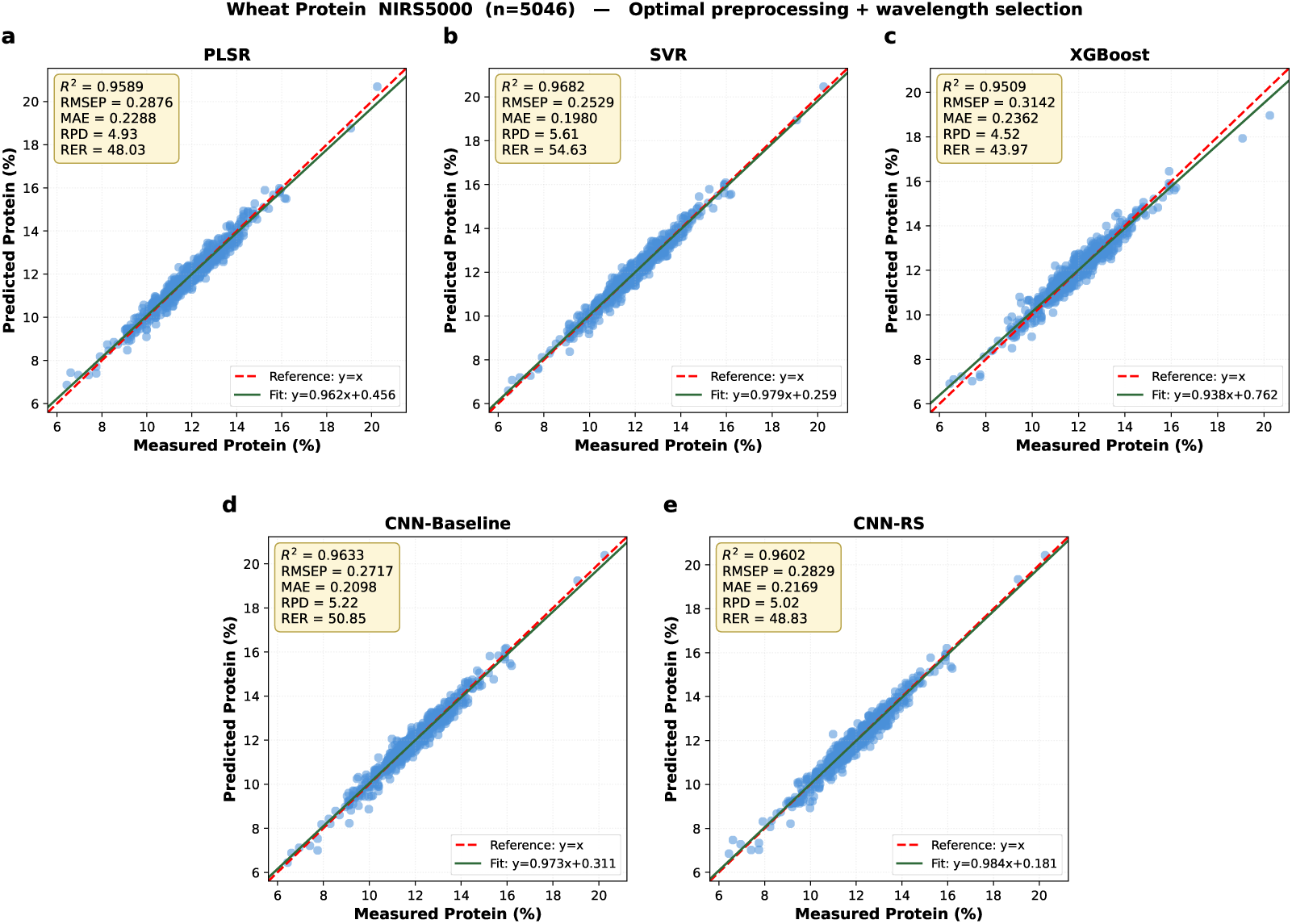
Measured versus predicted values on wheat protein (NIRS5000, *n* = 5046) with both preprocessing and wavelength selection: each model retains its preprocessing method selected for its validation set, while SVR, XGBoost and both CNNs share the wavelength subsets selected by PLSR and they are shown at their representative seeds; all other conventions are as in Figure 6.

**Figure 11:**
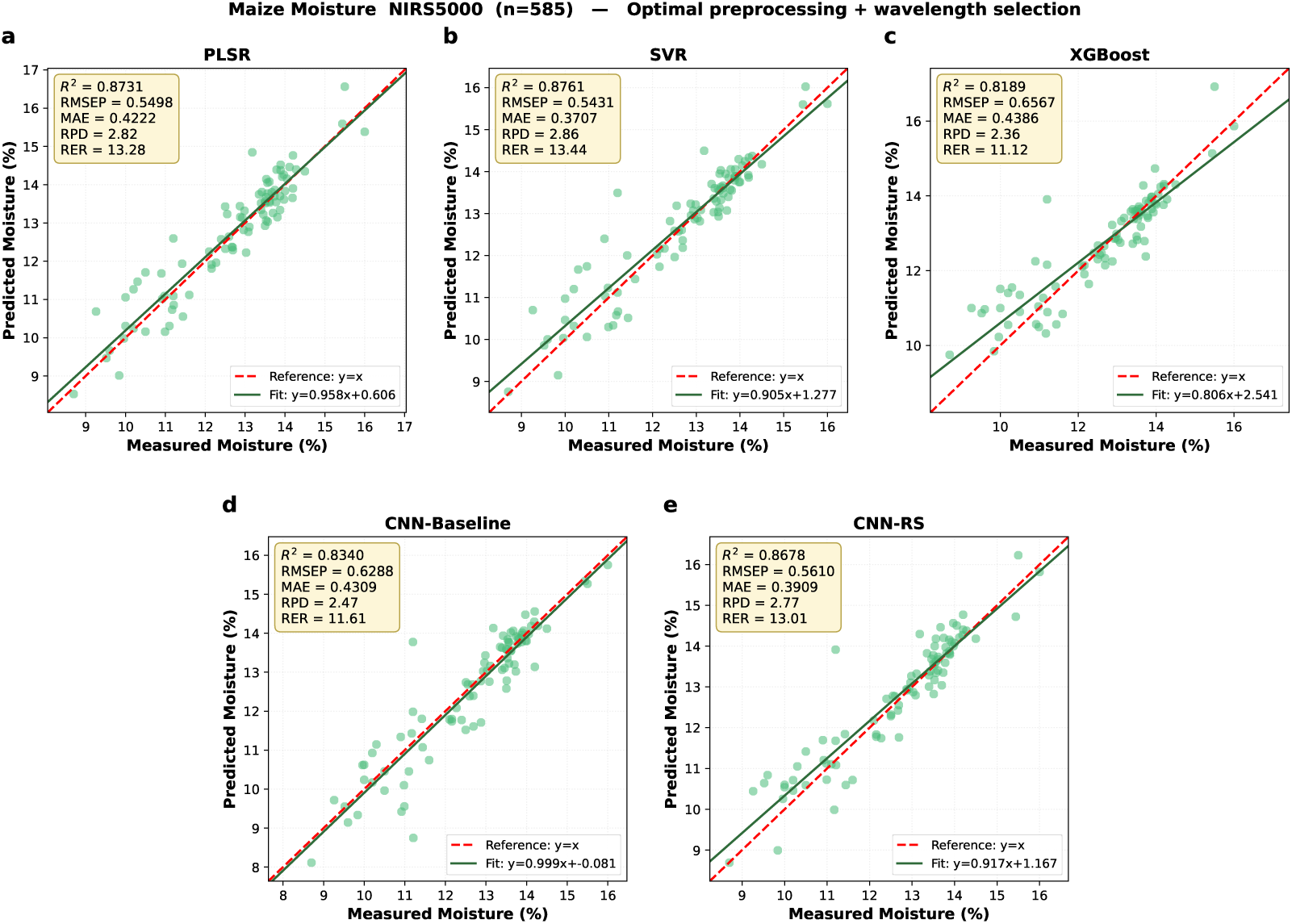
Measured versus predicted values on maize moisture (NIRS5000, *n* = 585) with both preprocessing and wavelength selection: SVR, XGBoost and both CNNs share the PLSR selected subset and PLSR uses its retained configuration. Test-set samples are shown in green; four seed-based models are shown at their representative seeds, and all other conventions are as in Figure 6.

Preprocessing corrected this directly. Compared to the unit slope, XGBoost presented a 74% reduction in mean deviation (from 0.421 to 0.111), while SVR saw a 61% reduction (from 0.106 to 0.041). Meanwhile, the worst case for XGBoost was a rise in the slope from 0.295 to 0.820 on wheat protein (XDS). Therefore, this improvement was reflected not only in the increase in explained variance but also in the restoration of the proportional relationship between predicted and measured values. PLSR’s response to preprocessing was similar to that of other models. For example, its fitting slope increased from 0.939 in the raw spectra to 0.958 after preprocessing in maize moisture. Wavelength selection resulted only small further changes.

Convolutional neural network fell in the middle and it was consistent with the sample-size dependence reported in Section 3.1. On the four larger datasets, their slopes on the raw spectra ranged from 0.94 to 0.99, and preprocessing had little effect on them. On the two smallest datasets, these values significantly lower: CNN-Baseline achieved values of 0.861 and 0.865 on the maize moisture and wheat protein (XDS) datasets, respectively, while CNN-RS achieved a value of 0.828 on the maize moisture dataset. Preprocessing raised the maize values to 0.977 (CNN-Baseline) and 0.940 (CNN-RS), and they became 0.999 and 0.917 with the wavelength subset added. When training data was sufficient, neural networks could recover proportional relationships without explicit correction. However, when training data was scarce, they cannot recover them, and this deficiency presented in scatter plot as a systematic offset rather than a diffuse cloud.

### 3.3. Performance and optimization of the baseline chemometric model (PLSR)

The PLSR model was optimized in two stages for each dataset. A comparison of 12 spectral preprocessing strategies (Table S1) showed that the best-performing strategy depended on the dataset. Preprocessing methods based on scatter and derivatives (e.g., SNV and second-order derivatives) were generally the most effective, indicating that the selection of appropriate preprocessing was crucial for linear models. Based on optimal preprocessing, a comparison of wavelength selection (Table S2) showed that restricting the model to a subset of retained wavelength information resulted in a slight improvement in prediction accuracy compared to full-spectrum predictions in some cases, which also yielded a more concise and interpretable calibration. Regression coefficient analysis (RCA, Figure S13) showed that the wavelength with the highest weight coincided with protein and water-related absorption regions in the NIR range, providing a chemical basis for the selected variables. Through these optimization steps, PLSR calibrated all six datasets to a competitive standard (Table 5). Test-set *R*^2^ exceeded 0.94 on five of the six datasets, with the strongest result on wheat protein (NIRS5000; *R*^2^=0.9589, RPD=4.93) and the weakest on maize moisture (*R*^2^=0.8731, RPD=2.81), the second-smallest dataset, and the one on which every model recorded its largest errors despite the widest constituent range (8.10-19.20%). Importantly, PLSR led wheat protein (XDS, n=500) in both the raw comparison (*R*^2^=0.9114) and the fully optimized one (*R*^2^=0.9448, RMSEP=0.2521), and led maize moisture (n=585) under preprocessing alone (*R*^2^=0.8807, RMSEP=0.5329). This reflected the small number of parameters and resistance to overfitting of linear models when training data were limited relative to the input dimensionality. In short, it provided a strong and consistent chemometric benchmark.

The optimized PLSR from our study is a stronger baseline than those commonly found in the literature. Because PLSR is a benchmark for evaluating machine learning and deep learning models, the rigor of its optimization process matters as much as that of the models compared against it. In many published studies, PLSR models have been improved through spectral preprocessing alone, and the spectral range is left unoptimized, which may result in a weaker baseline than what the method can achieve. In this study, both the preprocessing and wavelength selection for each dataset were optimized on the validation set. Wavelength subsets were retained on five of the six datasets and only barley protein was the full spectrum preferred. On test set, the subsets improved the *R*^2^ on three datasets, while slightly reduced it on barley moisture (−0.003) and maize moisture (−0.008), which was exactly what would be expected when selection was based strictly on the validation set.

On the small wheat protein (XDS, n=500) dataset, our optimized PLSR gave a lower test RMSEP than that reported by Shi and Yu (2017), although our test *R*^2^ (0.9448) was below their value (0.97). This difference was partly attributed to their much smaller and differently distributed calibration set (48 training and 23 test samples), since *R*^2^ was sensitive to variance of the reference values. On the small maize moisture dataset (n=585, 409 training, 88 validation and 88 test samples), our optimized PLSR outperformed the model of Shi and Yu (2017) on all metrics except RMSEP, and its test *R*^2^ of 0.8731 exceeded the values reported by Ferreira et al. (2014) and Zhang et al. (2023). These comparisons indicated that a PLSR baseline optimized over both preprocessing and wavelength selection provided a more competitive and appropriate benchmark for evaluating deep learning models than the only preprocessing that PLSR baseline commonly reported.

### 3.4. Machine learning reference models: SVR and XGBoost

#### Support vector regression

SVR was accurate and exceptionally stable, with virtually no run-to-run variability (Figure 6b; Table S3; Figures S1b-S4b). The *R*^2^ values for the test set ranged from 0.7433 for maize moisture to 0.967 for barley moisture. On larger datasets, it performed close to the strongest models (wheat protein: *R*^2^=0.9526, RPD=4.59; wheat moisture: *R*^2^=0.9500, RPD=4.47). The standard deviation across 25 seeds ranged from 0.0005 and 0.0098 in *R*^2^, and it was below that of both CNNs on five of the six datasets by roughly an order of magnitude relative to CNN-Baseline on the two smallest, with barley protein the exception (0.0098, above both CNNs), reflecting seeds for which the varying cross-validation folds selected different hyperparameters. This variance approaching zero was a structural property rather than a favorable result because SVR hyperparameters were selected through an exhaustive and deterministic grid search of 384 combinations. For fixed hyperparameters, the *ε*-insensitive loss with an RBF kernel yielded a convex optimization problem with a unique solution. The residual seed-to-seed variability therefore arises only where the varying cross-validation folds select different hyperparameters, as on barley protein. This run-to-run reproducibility is a practical advantage that accuracy alone does not capture. SVR proved more sensitive to the input representation than its accuracy on the larger datasets. After applying optimal preprocessing, the test *R*^2^ increased by 0.013 for wheat protein, by 0.057 for barley protein, 0.089 for wheat protein (XDS), and 0.136 for maize moisture (Table S3; Figures 8b, 10b, and S5b-S12b). The gain was systematically greatest where raw input performance was weakest, indicating that the gaps in Table 3 stemmed from scatter and baseline fluctuations rather than the kernel function’s inability to represent this relationship. The fitted slope also confirmed this trend: it increased from an average of 0.894 in the raw spectra to 0.959 after preprocessing, and even the value of worst case improved from 0.799 on wheat protein (XDS) to 0.918 (Section 3.2). After applying two input stages, SVR achieved the highest accuracy among the five models in four of the six datasets. Consequently, its relatively modest performance reflected the deliberate raw input conditions used in this comparison, rather than any inherent limitations of the method itself.

#### Extreme gradient boosting

XGBoost performed worst of the five models on each dataset under the unified raw input (Figure 6c, Table S4; Figures S1c-S4c). Accuracy was severely limited on the three protein datasets, with test-set *R*^2^ values of 0.6402 (wheat proteins), 0.4131 (barley proteins), and 0.2467 (wheat proteins, XDS), and moderate on the moisture datasets (0.8475, 0.8596, and 0.5491). In all cases, the model fitted the training data almost perfectly while generalizing poorly, which was a clear sign of over-fitting. The scatter plots also showed pronounced compression toward the mean. The fitted slope quantified this, 0.295 for wheat protein (XDS) and 0.412 for barley protein. This indicated that a sample one unit above the mean would have a predicted value only 0.3 to 0.4 units above the mean. A model exhibiting these characteristics was essentially unable to track this component. Two features in the data can explain this. Gradient-boosted trees performed segmentation on a single wavelength, so baseline shifts and scattering caused each sample to shift along each axis, and the thresholds learned during training could no longer distinguish the same samples in the test set (Chen and Guestrin, 2016). For proteins, this bias was more severe than water, because protein detection relied on weak and overlapping N-H features, whereas water had a strong and distinct absorption spectrum. For axis-aligned methods, the components that were most difficult to distinguish from the scattering baseline were precisely the ones they performed worst on.

After applying optimal preprocessing, XGBoost’s performance improved from 0.6402 to 0.9497 on the wheat protein dataset, from 0.4131 to 0.9416 on the barley protein dataset, and it improved from 0.2467 to 0.8666 on the wheat protein (XDS) dataset (Table S4; Figures 8c, 10c and S5c-S12c). The model performed the weakest in Table 3, which improved to a level that was only 0.02 behind the best-performing model. The mean fitted slope increased from 0.579 to 0.889, and the worst case rose from 0.295 to 0.820. As with SVR, the greatest improvement was observed when the raw input results were at their worst. Therefore, XGBoost was the model whose effectiveness depended entirely on explicit spectral correction. In addition, wavelength selection had little impact on other models, while it caused XGBoost’s accuracy on the wheat protein (XDS) dataset to drop from 0.8677 to 0.8416 (Table S4). This represented the most significant negative impact of wavelength selection observed in this study and it occurred on the smallest and the only dataset acquired with the XDS spectrometer.

### 3.5. Quantitative evaluation of 1D-CNN architecture

#### 3.5.1. Architecture, regularization and optimization

A systematic one-factor-at-a-time (OFAT) ablation study was conducted using a representative wheat protein (NIRS5000) dataset to determine the empirical effects of 13 structural and training hyperparameters (Table S5). The *R*^2^ on the validation set peaked with three convolutional layers, but the *R*^2^ for the single-layer architecture differed from this peak by only 0.005, and the single-layer architecture was ultimately retained in accordance with the principle of parsimony. Reducing the first layer to one filter both minimized the number of parameters and yielded the best validation metrics; a kernel width of 11 was optimal among odds widths from 3 to 19 and three fully connected layers maximized the accuracy. In the width test, the largest configuration achieved the highest validation *R*^2^ value, but the compact [32, 18, 12] block attained the same *R*^2^ value on the validation set while reducing the number of parameters by more than two-thirds, and was therefore retained. Among the regularization and optimization factors, dropout led to a decline in performance at every tested rate, while batch normalization proved to be critical, which the network could not be trained at the standard learning rate without it, and the validation R² value dropped by nearly 0.22. Weight decay performed best at zero, with a high penalty leading to divergence. Adam converged reliably where SGD underfitted the spectral profile, a batch size of 256 gave the most stable convergence, and ELU outperformed the five other activation functions tested. Average pooling with a kernel of 2 matched the accuracy of no pooling at a fraction of the parameter count. The resulting CNN-Baseline was specified as a single filter, single convolution layer with kernel size of 11, batch normalization, ELU activation and average pooling, without dropout or weight decay (Table 2, Fig. 2a). Such a minimal architecture was sufficient, and this finding was consistent with Cui and Fearn (2018), who showed that compact CNNs were very suited for NIR calibration and with practical guidance that emphasis target network design over raw model capacity (Yang et al., 2019). This suggested that deep and high-capacity architectures borrowed from image classification were unnecessary for one-dimensional NIR spectroscopy.

#### 3.5.2. Spectral preprocessing interactions

The impact of traditional row-wise preprocessing on the 1D-CNN was investigated on the representative wheat protein dataset (NIRS5000, n=5046; Table S6). Raw spectra with column-wise scaling alone provided the highest validation-set accuracy, and none of the twelve preprocessing strategies improved on it. Across all six datasets, the benefit of preprocessing was inversely proportional to the sample size: slightly negative on the largest, negligible to modest on the intermediate ones, and substantial on the two smallest, where it raised test R² by 0.034 (wheat protein, XDS) and 0.047 (maize moisture) for CNN-Baseline and by 0.017 and 0.055 for CNN-RS. This trend was not strictly monotonic. This test *R*^2^ value for CNN-Baseline increased by 0.010 on the wheat moisture (n=3305) while CNN-RS showed no improvement. Therefore, the independence of the convolutional network from explicit preprocessing is real but conditional on sample size.

This finding was consistent with previous studies that CNNs depended far less on preprocessing than PLSR (Acquarelli et al., 2017; Bjerrum et al., 2017; Cui and Fearn, 2018). Cui and Fearn (2018) showed that simple CNNs applied directly to raw NIR spectra match or exceed conventional calibrations. Acquarelli et al. (2017) also reported that trained convolutional filters reproduce the smoothing and differentiation of classical preprocessing and can replace it, and that applying preprocessing anyway sometimes reduce the performance. Bjerrum et al. (2017) similarly advocated learning these operations within the network instead of applying them by hand. Passos (2026) argued that when using convolutional neural networks (CNNs) to analyze near-infrared (NIR) spectra, learning robust alternative preprocessing models from the data required sufficient signal information. However, the small-scale datasets commonly encountered in near infrared chemometrics might not provide enough signal for neural network to learn the optimal transformation while performing prediction tasks. Furthermore, he also argued that scattering corrections such as SNV and MSC primarily addressed physical effects that were essentially orthogonal to chemical labels, so neural networks trained using chemical-supervised loss had little incentive to reliably learn these effects. Both predictions were consistent with our observations: the substitution strategy held on the four datasets with more than 900 samples and broke down on the two with fewer than 600. An observational pattern across datasets also differ in spectral range and constituent spread (Section 3.1), not a controlled sample-size threshold.

Preprocessing also allowed the network to reach its optimal state in far fewer epochs while maintaining the same accuracy. Across 12 model-dataset combinations (two networks × six datasets), the number of epochs required to achieve the minimum validation loss decreased in 11 combinations, with a median reduction of 49% (Wilcoxon signed-rank test on the paired epochs, p = 0.0010). Taking wheat protein as an example, the number of training epochs for the CNN-Baseline decreased from 1256 to 157. The noise in the validation curves of the raw input was also significantly greater, with recurring fluctuations of one to two orders of magnitude, while these fluctuations virtually disappeared after preprocessing (Figures 12 and 13). It suggested that scatter and baseline corrections simplified the learning problem, as the network had to learn to ignore these effects by itself. When the training set was large, the convolutional layers still perform this correction internally. For smaller datasets, when training efficiency was a priority, spectral preprocessing remained beneficial.

**Figure 12:**
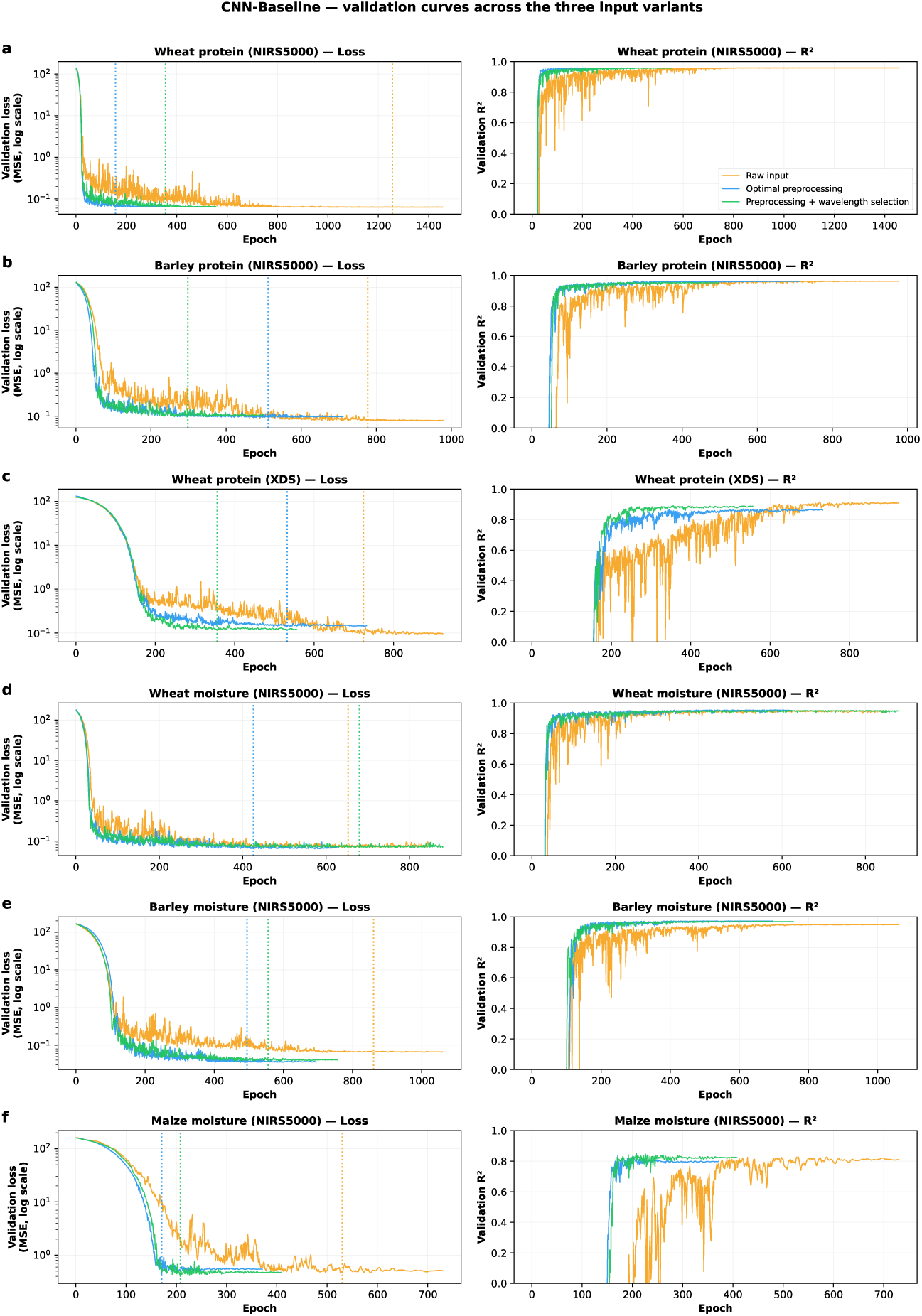
Validation curves of the CNN-Baseline model under the three input conditions. Orange, raw input; blue, optimal preprocessing across the full spectrum; green, preprocessing combined with wavelength selection. Left panels show the validation loss (mean square error, on a logarithmic scale), while the right panels show the validation *R*^2^, both against epoch. The vertical dashed lines mark each condition’s minimum validation loss, and early stopping ends the three curves at different epochs. Curves are for the representative seed of each dataset and condition. For barley protein, the preprocessed and fully optimized inputs are consistent. The full spectrum having been retained, so the green curve overlaps the blue.

**Figure 13:**
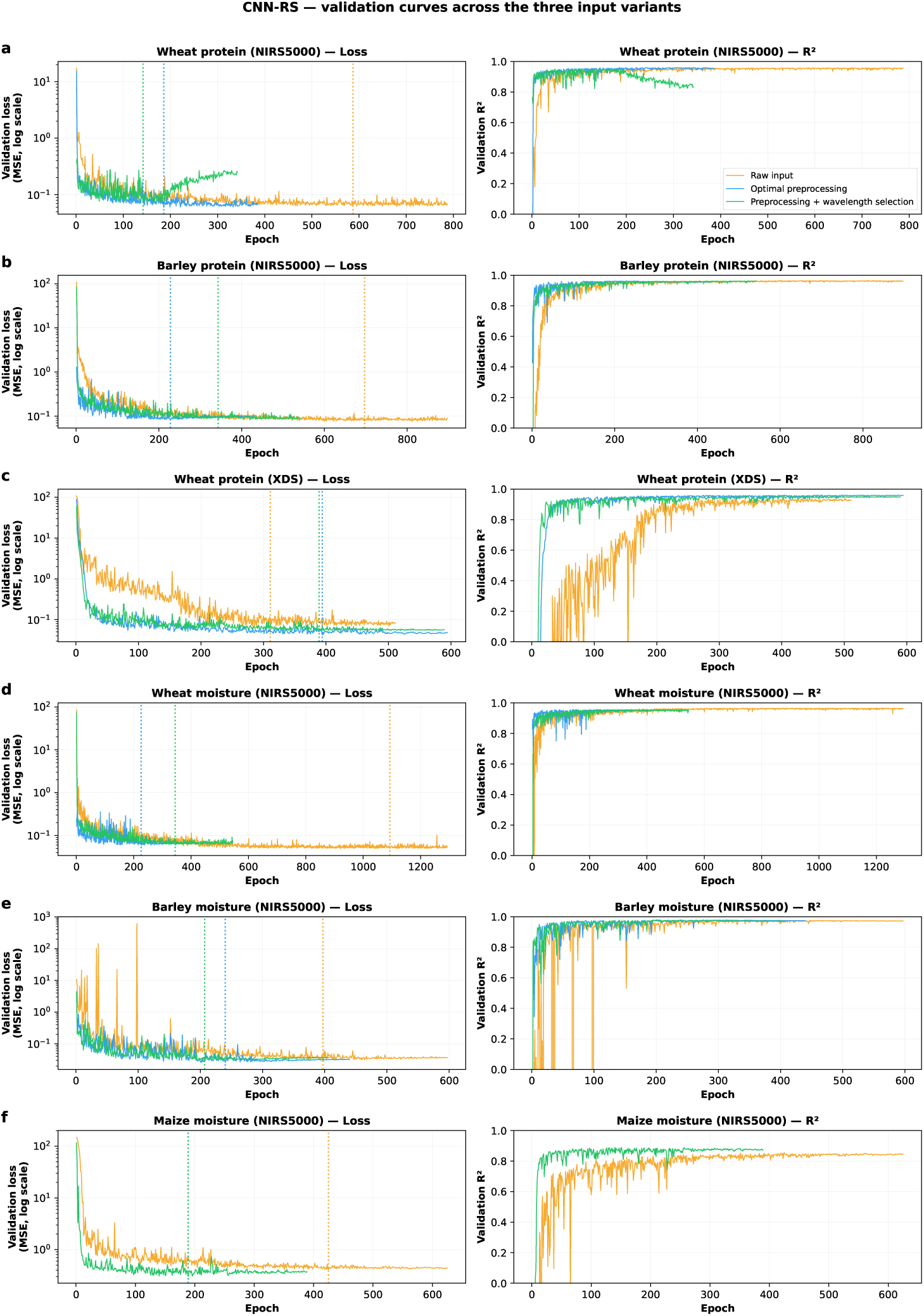
Validation curves of CNN-RS under three input conditions; panels, colors and conventions as in Figure 12. The architecture is searched independently for each input condition, so each curve reflects the joint effect of the input and the architecture selected for it.

### 3.6. CNN performance: CNN-Baseline versus CNN-RS

The final lightweight architecture (CNN-Baseline; Table 2, Figure 2a) was evaluated alongside the automatically searched model (CNN-RS) on all six datasets, each with 25 independent initialization seeds. Both networks trained stably, with loss decreasing steadily and the validation *R*^2^ converging before the early-stopping threshold was reached (Figures 14 and 15), resulting in a compact scatter plot (Figure 6d, e; Figures S1-S4). Under the unified raw input, CNN-RS achieved higher accuracy than CNN-Baseline on all six datasets, but by margins that depended strongly on sample size. On the four larger datasets, it exceeded CNN-Baseline by between 0.003 to 0.017 in test *R*^2^. These margins were comparable to the seed-to-seed standard deviation of CNN-Baseline itself. The advantage of CNN-RS was more pronounced for the two smallest datasets. The test *R*^2^ values were 0.9020 for CNN-RS and 0.8464 for CNN-Baseline on the wheat protein measured by XDS (n=500), while the corresponding values were 0.8178 and 0.7954. The selected architectures were listed in Table S7, and the complete CNN-RS results were provided in Table S8.

**Figure 14:**
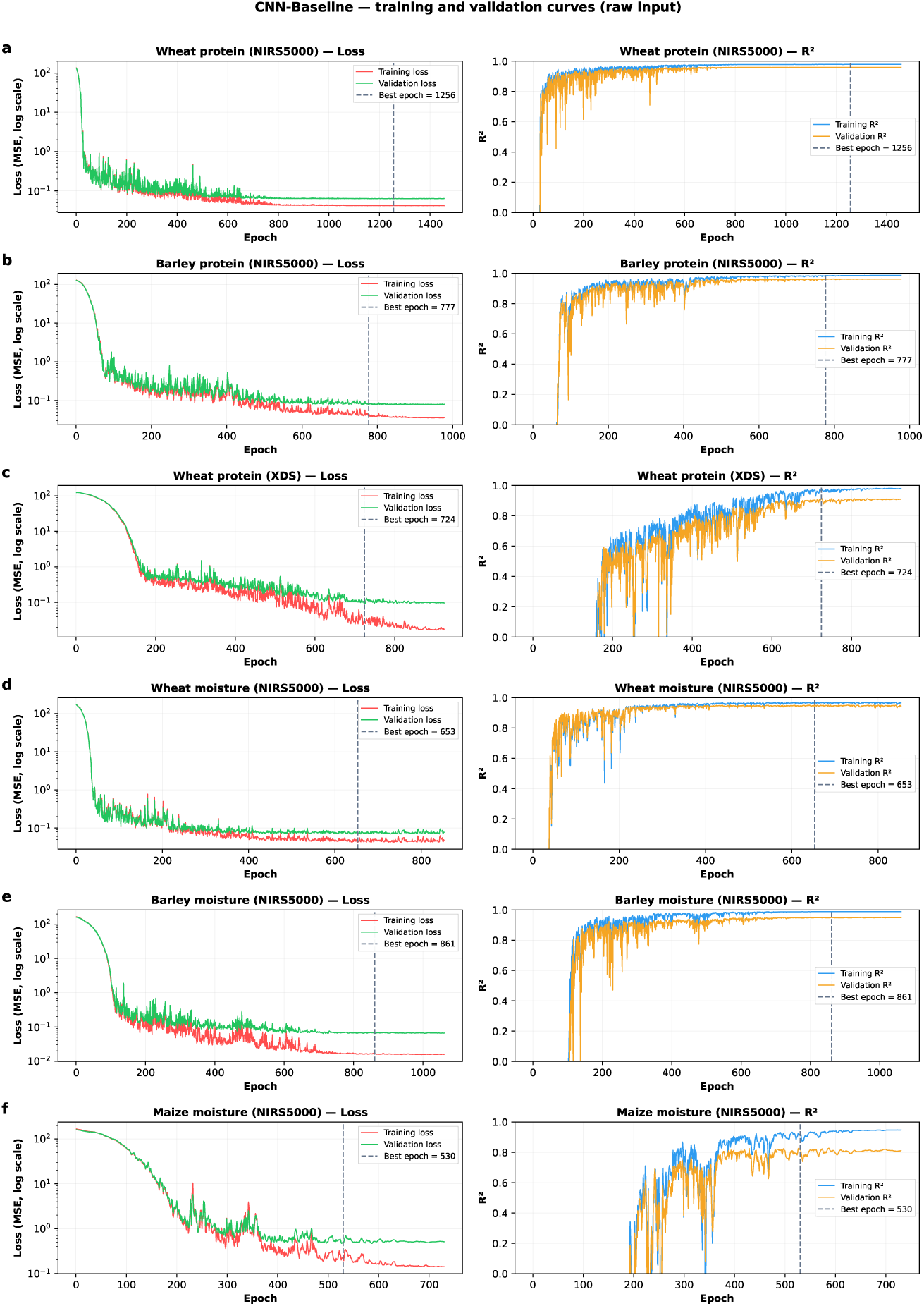
Training and validation curves of the CNN-Baseline model (the architecture derived from the one-factor-at-a-time ablation study) on the unified raw input. For each dataset, the left panel shows training (red) and validation (green) loss (mean square error, logarithmic scale) and the right panel shows training (blue) and validation (orange) *R*^2^, both against training epoch. The vertical dashed line marks the epoch of minimum validation loss, whose weights were retained for evaluation. Curves are shown for the representative seed of each dataset. Training runs for up to 3000 epochs with early stopping after 200 epochs without validation improvement, so the curves end shortly after the marked epoch.

**Figure 15:**
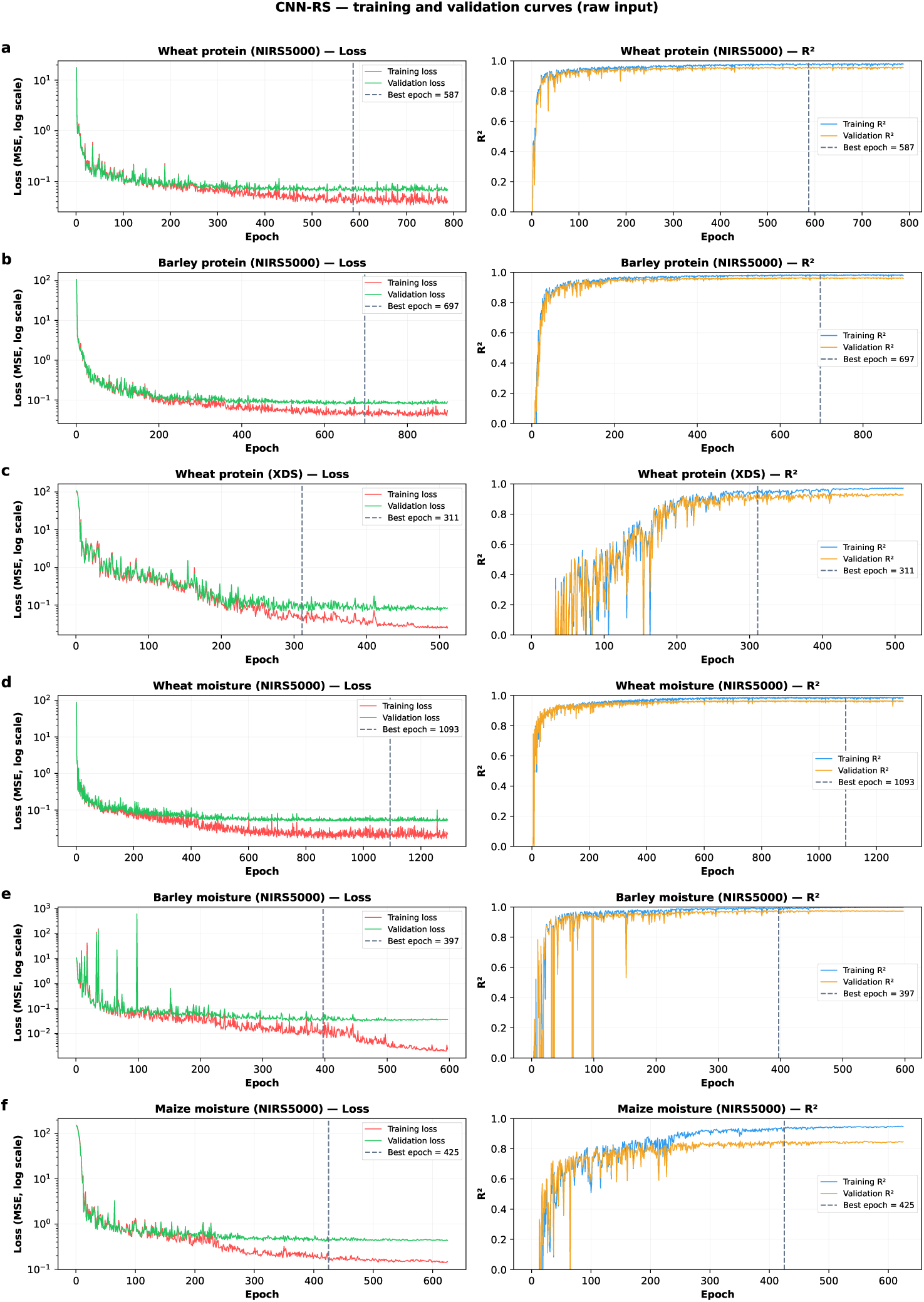
Training and validation curves of CNN-RS model on the unified raw input; panels, colors and conventions as in Figure 14. The architecture search was repeated independently for each input condition, so the architectures shown here were selected on uncorrected spectra and differed between datasets.

Reproducibility showed the same overall trend. For CNN-Baseline, the standard deviation of test *R*^2^ increased from 0.0026 on the largest dataset to about 0.0440 on both of the smallest datasets, while the standard deviation of *R*^2^ for CNN-RS remained below 0.026. Thus, CNN-Baseline not only exhibited lower accuracy but also significantly poorer reproducibility under sample limited conditions. This aligned with the documented data requirements for high-capacity models (Yang et al., 2019; Zhang et al., 2021) and was the reason for proposing spectral augmentation strategies (Bjerrum et al., 2017).

The conclusion is not that architecture search is unnecessary, but that its values is conditional on data volume. Where several thousand calibration samples are available, the single-filter architecture obtained through ablation studies achieved *R*^2^ values within only a few thousandths from the result of searching through 300 candidate architectures, and the computational cost is far lower than the latter. This aligns with practical guidelines that emphasize the superiority of domain-knowledge-based design over brute-force search (Yang et al., 2019). It also aligns with the conditional framework proposed by Passos (2026), which identified sample size, data augmentation, and regularization as moderating variables that determine whether increasing depth or capacity was beneficial. Architecture search becomes more worthwhile when only a few hundred samples are available.

A further observation concerns the behavior of architecture search. Across the eighteen searches reported in Table S7, only six distinct architectures are selected, with two of them accounting for 13 of the 18 selections. This indicates that the study consistently identifies a relatively small region within the architecture space, rather than different solutions for each set of conditions. Furthermore, the optimal configuration does not depend on the input: the number of convolutional layers was reduced in two datasets compared to original spectrum, while it remains unchanged in other four. These findings suggest that only a few configurations consistently perform well across different grain types, compositions, instruments, and input representations, rather than each condition having its own unique optimum. The ranking within that leading group was less dependable: the configuration ranked first on the validation set was not the best over 25 seeds on wheat protein (XDS), the third-ranked configuration achieving a higher average test *R*^2^ value despite a lower single-seed validation performance. Overall, these results support the use of pre-specified and compact architectures in practice, particularly when multiple candidate configurations exhibit similar performance. Architecture search remains valuable, especially for smaller datasets, but its better results should be viewed as strong candidates rather than the single optimal solution.

## 4. Conclusions

This study examined how architecture and input preparation govern model choice in the near-infrared calibration of cereal grains. A simple one-dimensional convolutional neural network (1D-CNN) was first derived by one-factor-at-a-time ablation, consisting of a single convolutional layer with one filter, a kernel size of 11 and batch normalization, and was then benchmarked against PLSR, SVR, XGBoost, and a random-search CNN (CNN-RS) across six datasets. Every model except the deterministic PLSR was evaluated over 25 random seeds, under three input conditions: raw spectra, optimal preprocessing, and the preprocessing combined with wavelength subsets.

Under a raw input, CNN-RS was the most accurate model on five of the six datasets, and PLSR on the smallest protein dataset (XDS), which is also the only cross-instrument dataset and the one with narrowest constituent range, so this single reversal cannot be attributed to sample size alone, and CNN-RS model exceeded the CNN-Baseline on every dataset under this input. When training data was abundant, the *R*^2^ value of a single-filter architecture obtained without any search came within a few thousandths of *R*^2^ of a 300-candidate search. However, when training was scarce, the cost of the search was well worth it, and the repeatability of the CNN-Baseline model was significantly lower, with the standard deviation rising from 0.003 to 0.044.

The rankings also depended largely on the data each model receives. When each model was provided with its own preprocessing and wavelength subset, SVR performed most accurately on four of the six datasets, while XGBoost rose from being the worst-performing model in the benchmark to within 0.02 of the best, with its test *R*^2^ value on the barley protein dataset rising from 0.413 to 0.941. Therefore, the advantage of convolutional neural networks over raw spectra lay in their robustness to unprocessed input data rather than in attainable accuracy. On the four larger datasets, preprocessing had no effect on the accuracy of CNNs, but it improved accuracy by 0.034 and 0.047 on the two smallest datasets, respectively. Furthermore, it brought the networks to their optimum in fewer epochs in 11 out of 12 model-dataset combinations, with a median reduction of 49%. The claim that convolutional layer could replace explicit preprocessing holds, but conditionally: it required sufficient training data, and even where accuracy was unaffected, preprocessing reached the optimum in about half the epochs and made the early-stopping point more predictable. The preprocessing methods required for each model also varied. The strategies independently selected for convolutional neural networks (CNNs) differed from most of those chosen by PLSR, and this difference was systematic: PLSR selected the second-order derivative strategy on five of the six datasets, while CNNs selected the first-order derivative strategy for five of the six datasets. Preprocessing workflows optimized for linear models should not be directly applied to neural networks, which could perform their own filtering.

Overall, these results supported the approach that balanced sample size and input features when selecting a calibration model for cereal grain NIR spectra. With a few hundred samples, an optimized PLSR or SVR models were more suitable and stable. With several thousand, the compact 1D-CNN performed at least as well, without the need to search for preprocessing methods, and additional network capacity brought little further benefit. More broadly, the results support the following principle: simpler models should generally be preferred for one-dimensional spectral data, provided the input each model received was stated explicitly. Otherwise, the comparison reflected the data preprocessing workflow as much as the models themselves.

## Acknowledgements

The Natural Sciences and Engineering Research Council of Canada (NSERC-Individual Discovery Grant and NSERC-CRD Grant), the Ministry of Agriculture Strategic Research Chair Program Fund and the Agricultural Development Fund (ADF) are acknowledged.

## Declaration of generative AI and AI-assisted technologies in the writing process

During the preparation of this work the author used ChatGPT-5.6 (OpenAI) and Claude-5 (Anthropic) to refine the language, improve coherence and clarity of the manuscript. After using this tool/service, the author reviewed and edited the content as needed and takes full responsibility for the content of the publication.

