## Supplemental Material for "One-dimensional CNNs for Near-Infrared Prediction of Protein and Moisture in Cereal Grains: The Effects of Architecture and Input Preparation"

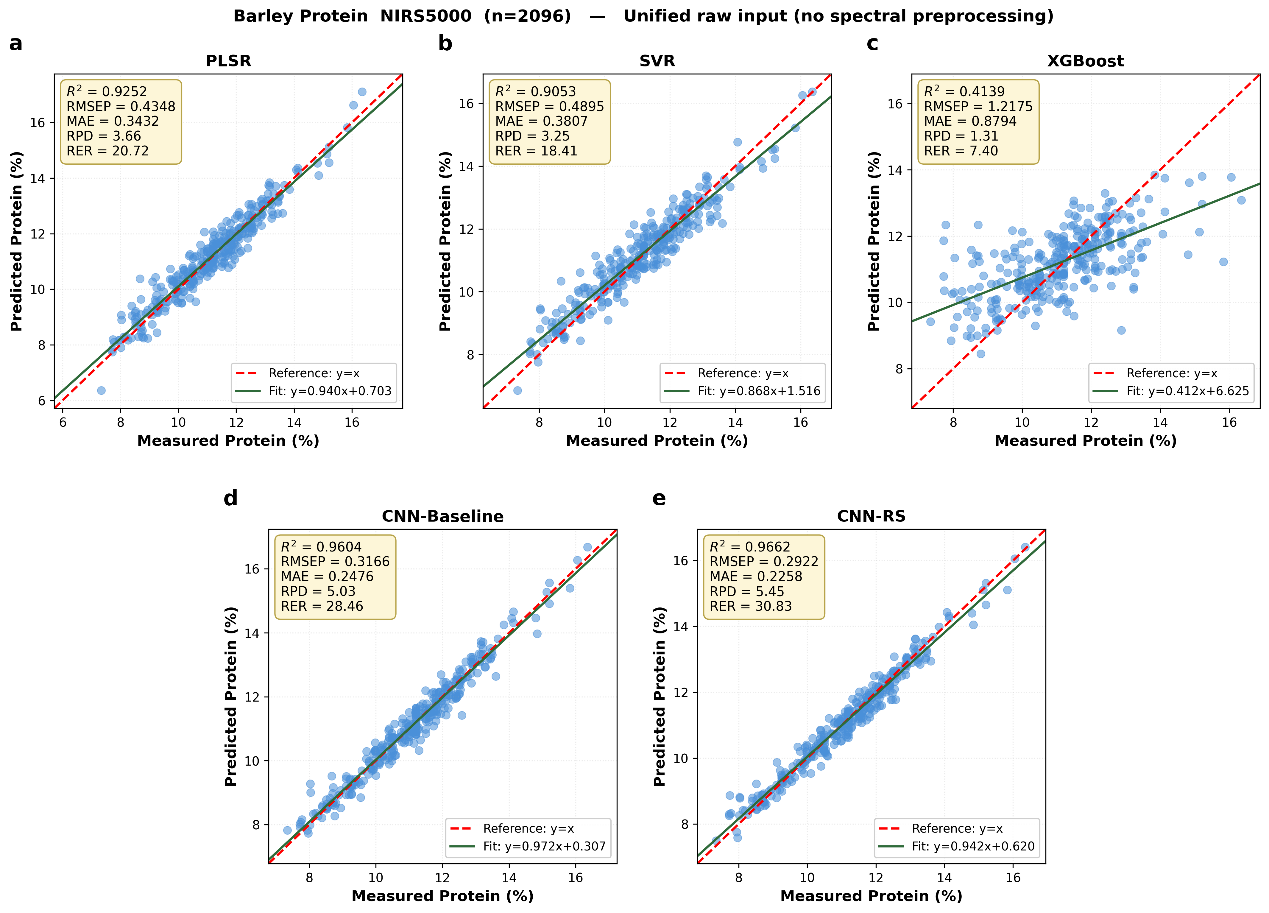

**Figure S1.** Measured versus predicted values of the five models on barley protein (NIRS5000, n=2096) under the unified raw input. Test-set samples are presented as blue circles; SVR, XGBoost and both CNNs are shown at their representative seeds. Plotting conventions, inset statics and the choice of run follow Figure 6.

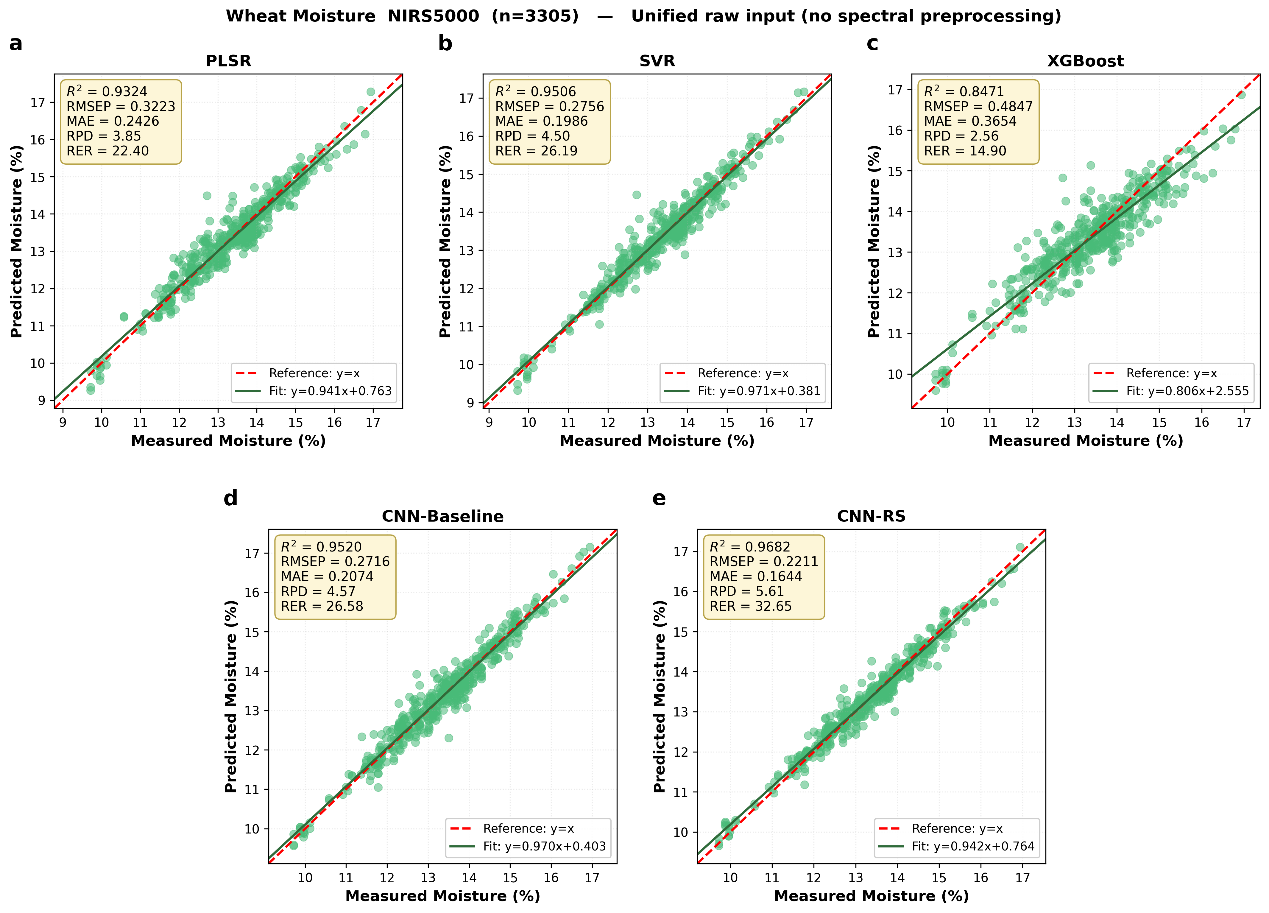

**Figure S2.** Measured versus predicted values of the five models on wheat moisture (NIRS5000, n=3305) under the unified raw input. Test-set samples are shown as green circles; SVR, XGBoost and both CNNs are shown at their representative seeds. Conventions and the choice of run are as in Figure 6.

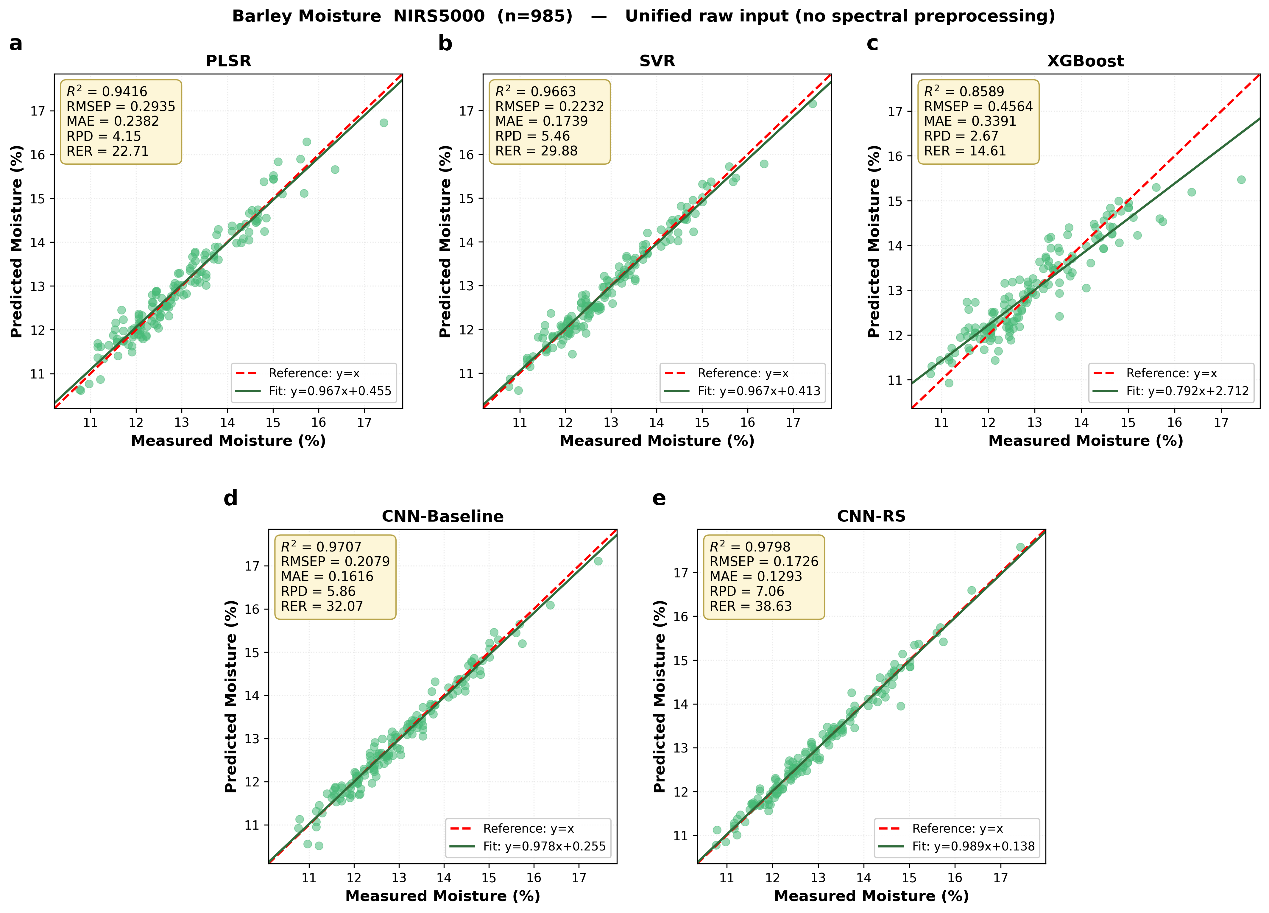

**Figure S3.** Measured versus predicted values of the five models on the barley moisture (NIRS5000, n=985) under the unified raw input. Test-set samples are shown as green circles; SVR, XGBoost and both CNNs are shown at their representative seeds. Conventions and the choice of run are as in Figure 6.

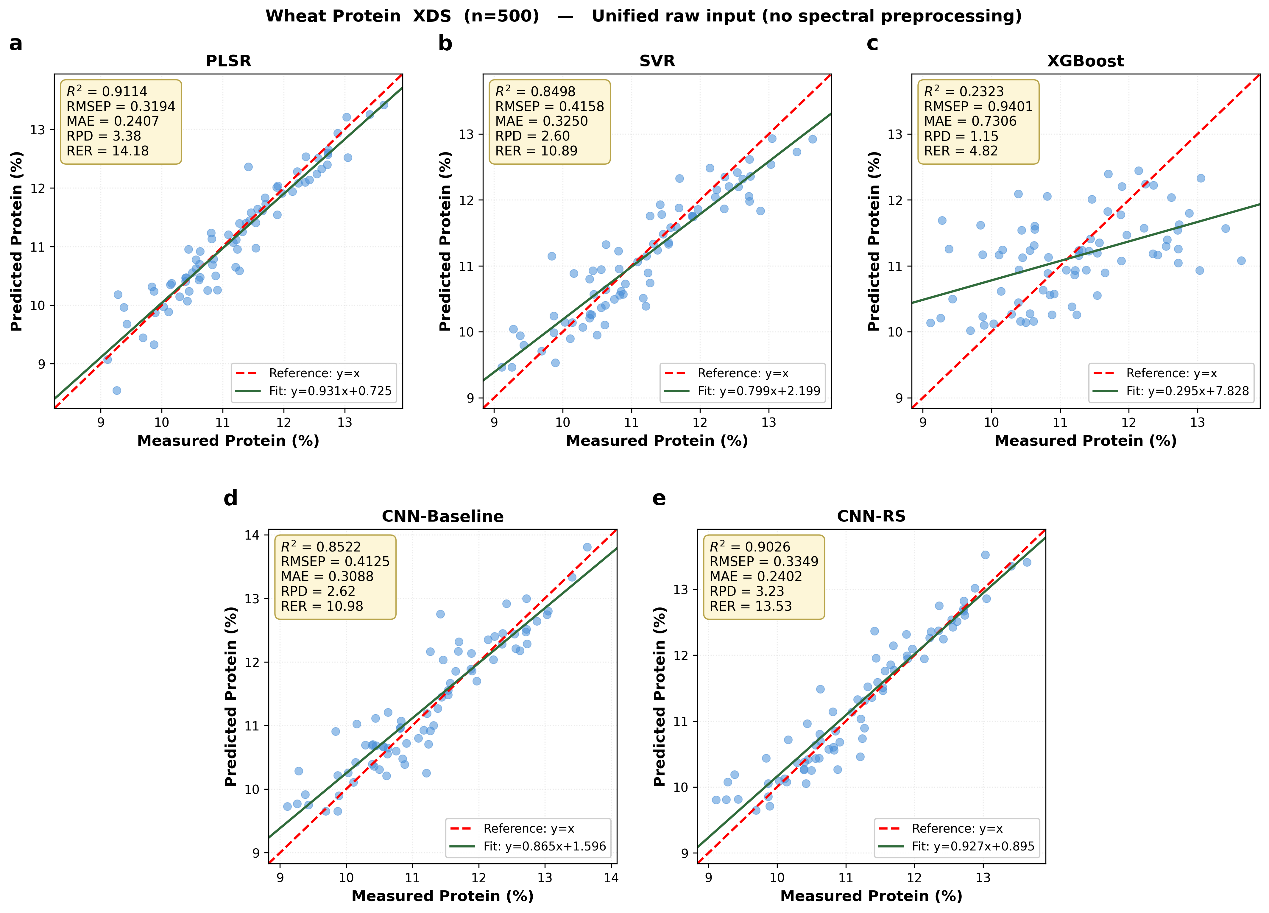

**Figure S4.** Measured versus predicted values of the five models on wheat protein (XDS, n=500), which contains 1050 wavelengths ranging from 400-2498 nm, under the unified raw input. Test-set samples are shown as blue circles; SVR, XGBoost and both CNNs are shown at their representative seeds. Conventions and the choice of run are as in Figure 6.

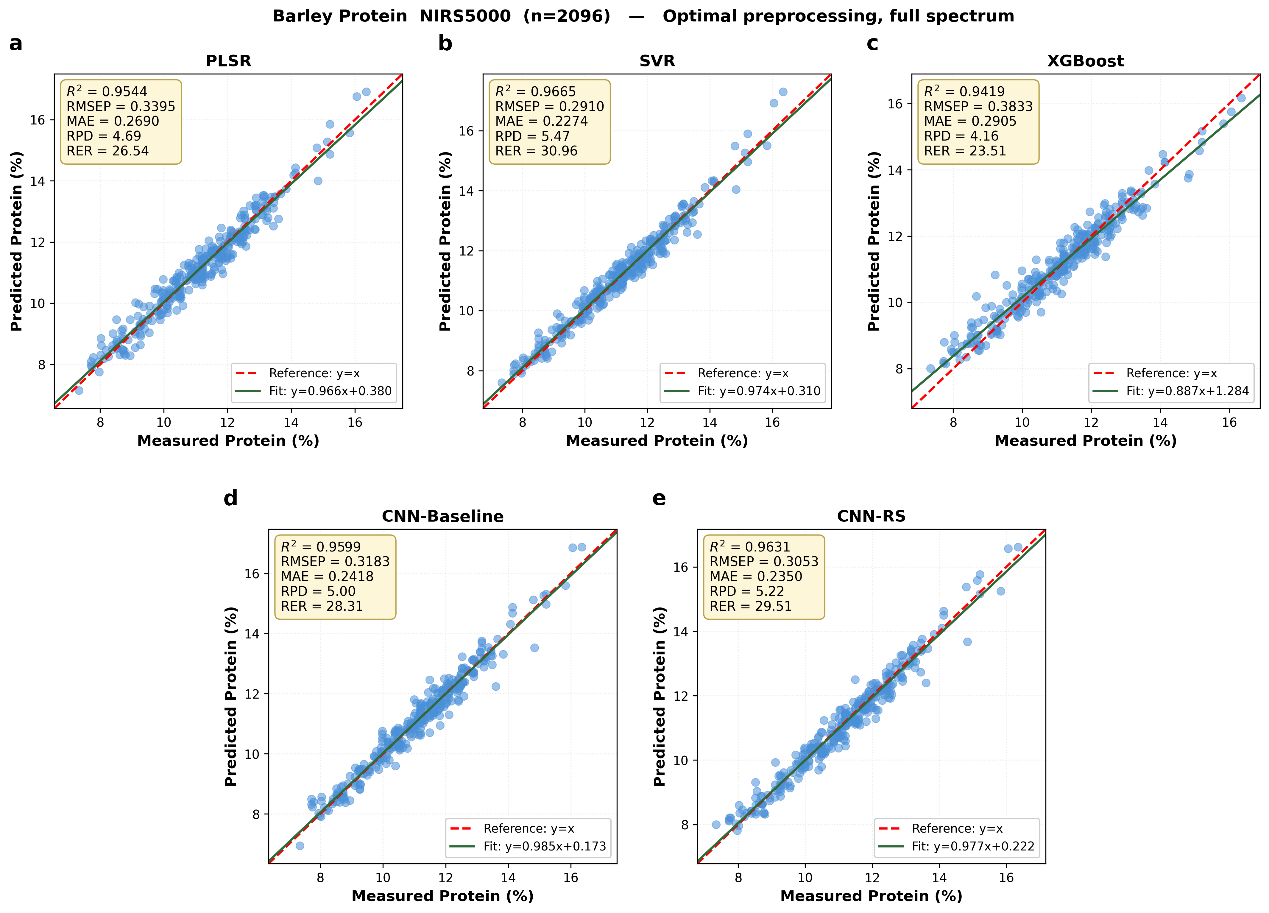

**Figure S5.** Measured versus predicted values of the five models on barley protein (NIRS5000, n=2096) with optimal preprocessing methods for each model on the validation set to the full spectra. Test-set samples are shown in blue; SVR, XGBoost and both CNNs are shown at their representative seeds, and other conventions are as in Figure 6.

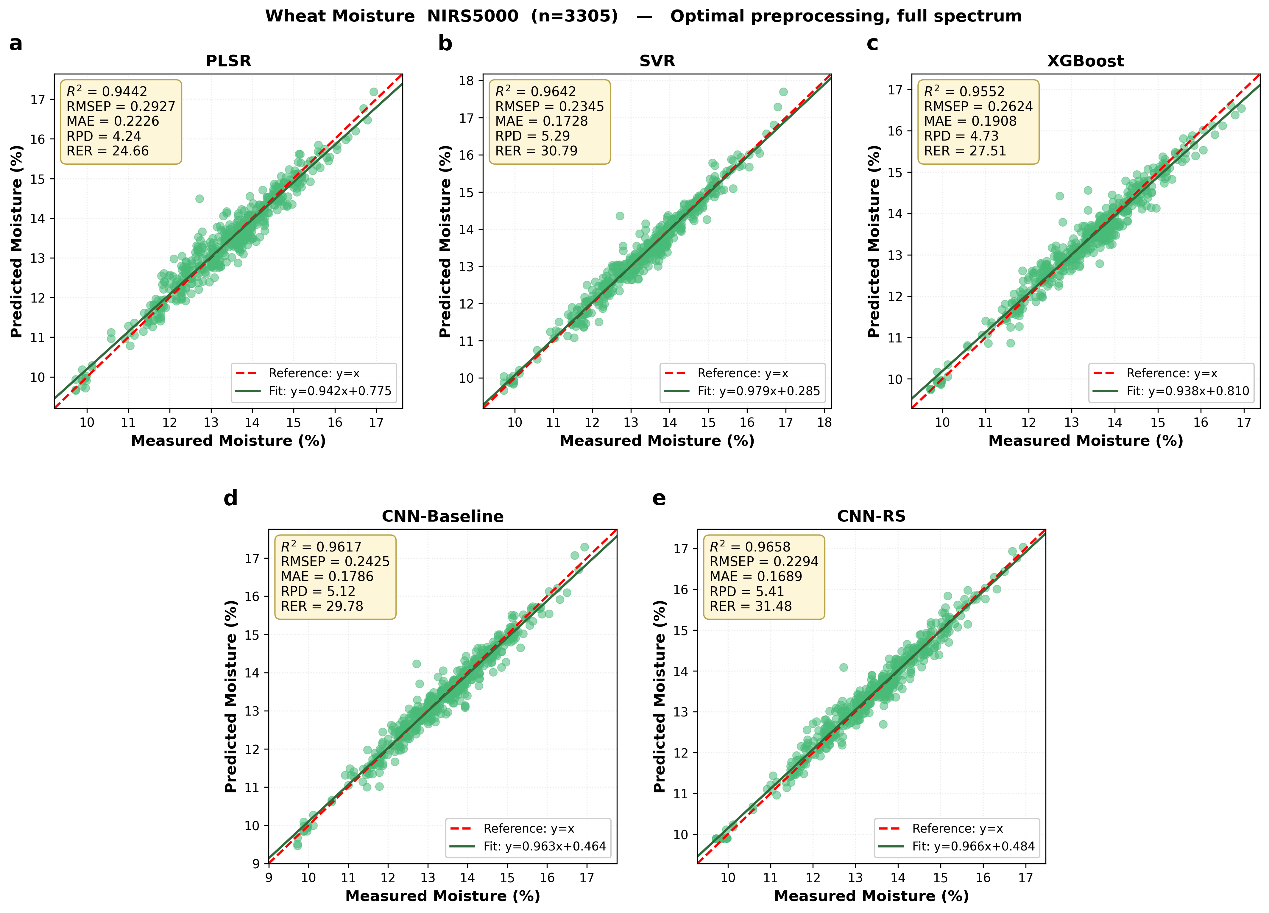

**Figure S6.** Measured versus predicted values of the five models on wheat moisture (NIRS5000, n=3305) with optimal preprocessing methods for each model on the validation set to the full spectra. Test-set samples are shown in green; SVR, XGBoost and both CNNs are shown at their representative seeds, and other conventions are as in Figure 6.

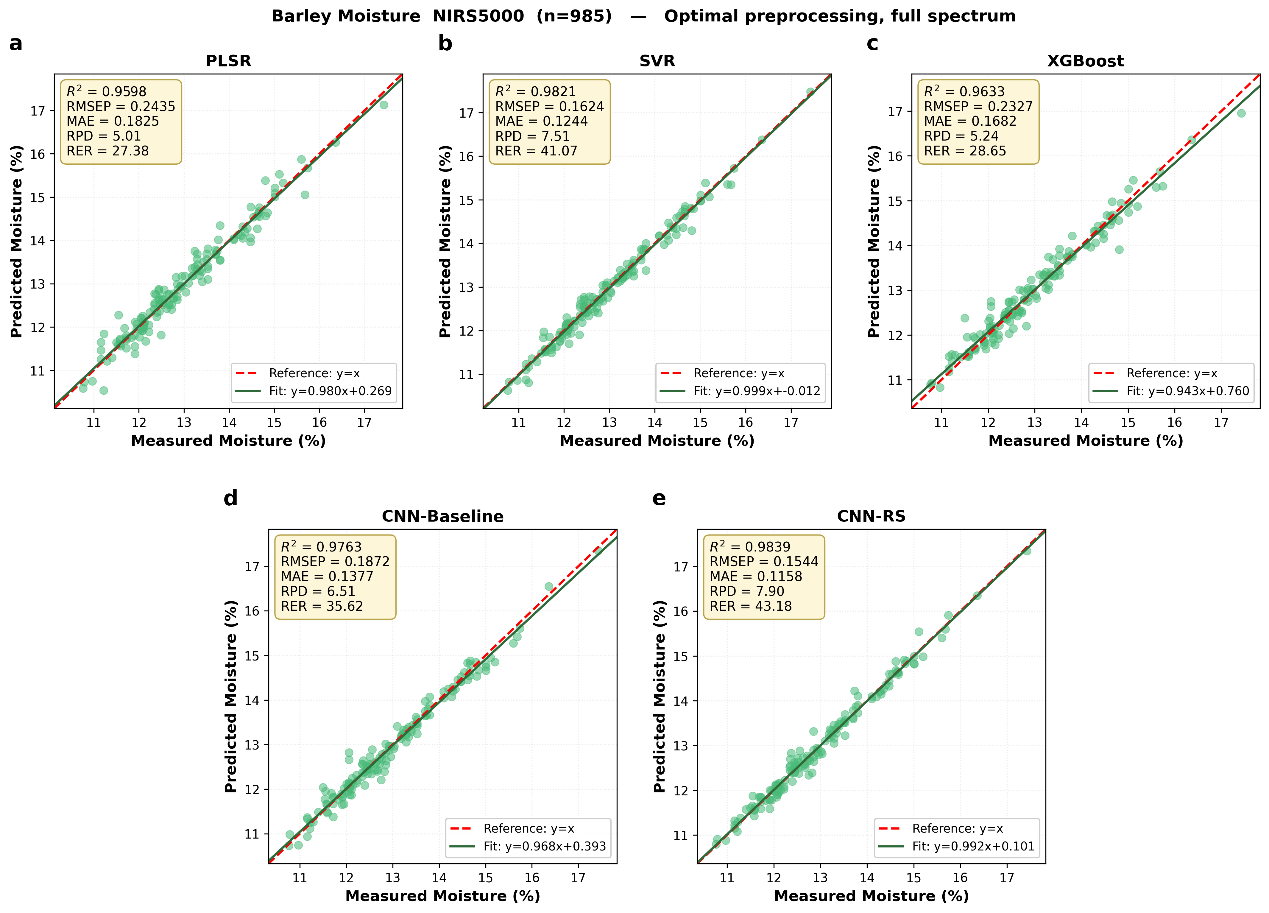

**Figure S7.** Measured versus predicted values of the five models on barley moisture (NIRS5000, n=985) with optimal preprocessing methods for each model on the validation set to the full spectra. Test-set samples are shown in green; SVR, XGBoost and both CNNs are shown at their representative seeds, and other conventions are as in Figure 6.

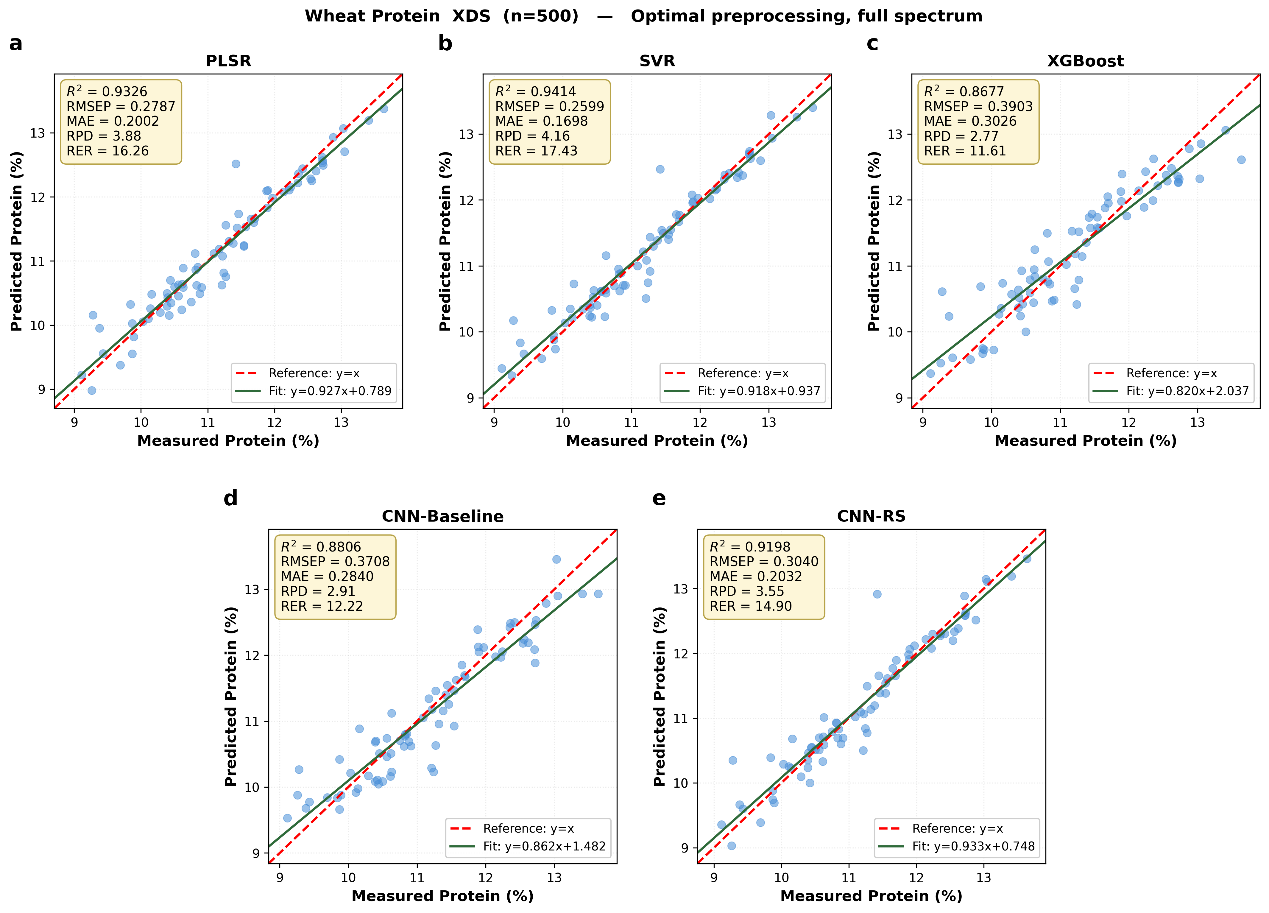

**Figure S8.** Measured versus predicted values of the five models on wheat protein (XDS, n=500) with optimal preprocessing methods for each model on the validation set to the full spectra. Test-set samples are shown in blue; SVR, XGBoost and both CNNs are shown at their representative seeds, and other conventions are as in Figure 6.

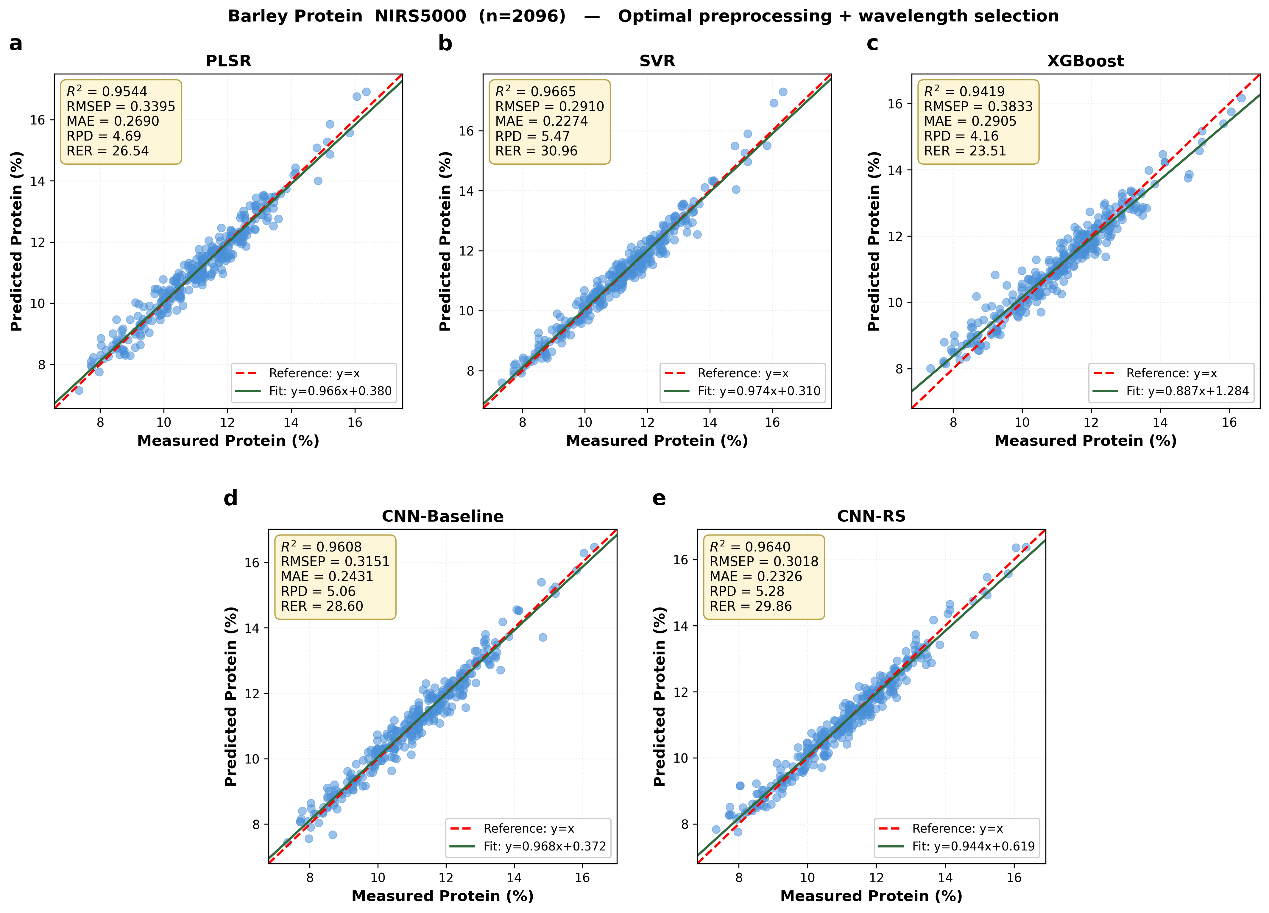

**Figure S9.** Measured versus predicted values of the five models on barley protein (NIRS5000, n=2096) with both optimal preprocessing and wavelength selection. The validation-based selection retained the full spectrum for this dataset, so the fully optimized input is same with the preprocessed one: the PLSR, SVR and XGBoost panels are same with Figure S5, while the CNN panels are retrained and CNN-RS is also researched under this condition, so the results can differ slightly. Test-set samples are shown in blue; all other conventions are as in Figure 6.

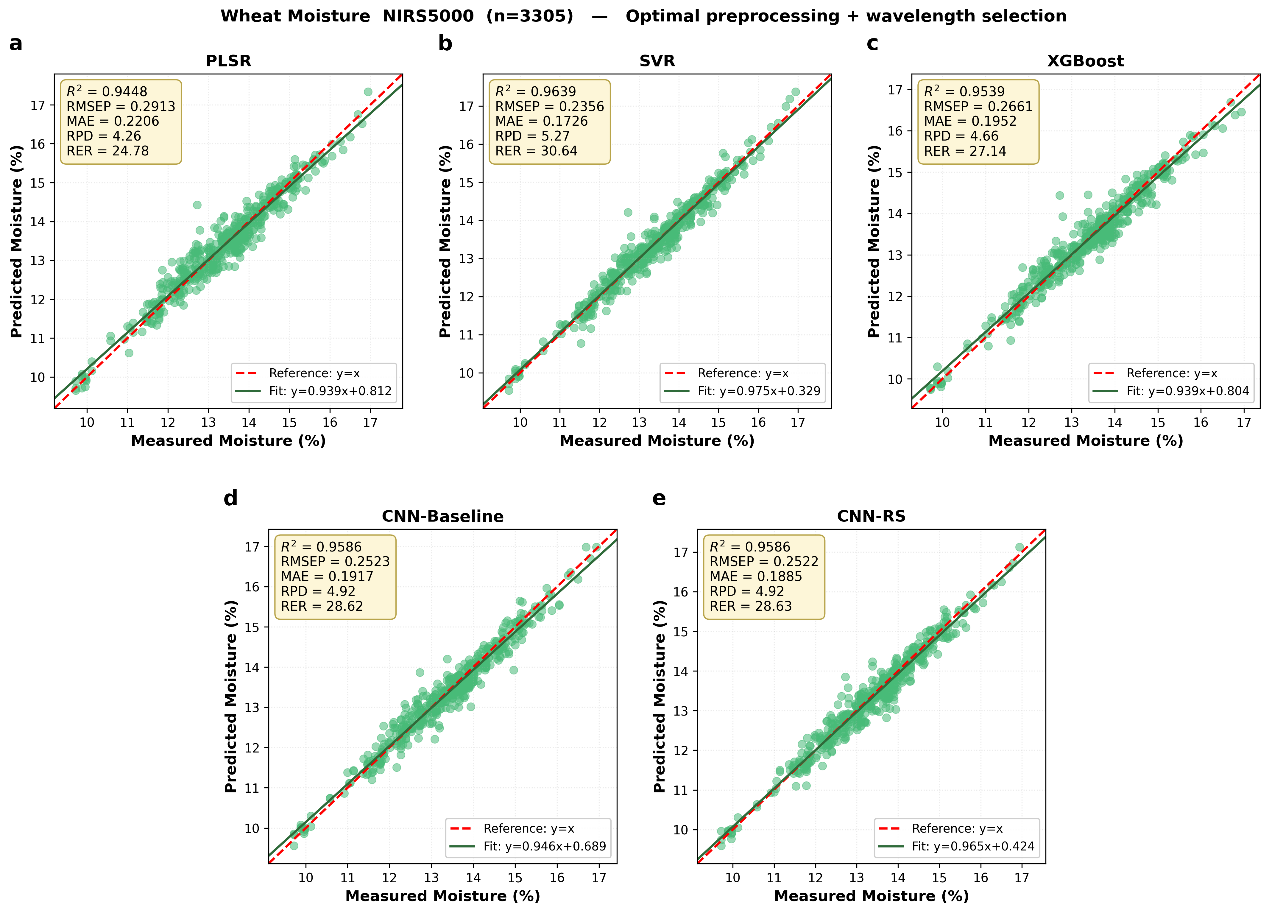

**Figure S10.** Measured versus predicted values of the five models on wheat moisture (NIRS5000, n=3305) with both optimal preprocessing and wavelength selection: SVR, XGBoost and both CNNs share the wavelength subset derived from the PLSR regression coefficients for this dataset, shown at their representative seeds. Test-set samples are shown in green; all other conventions are as in Figure 6.

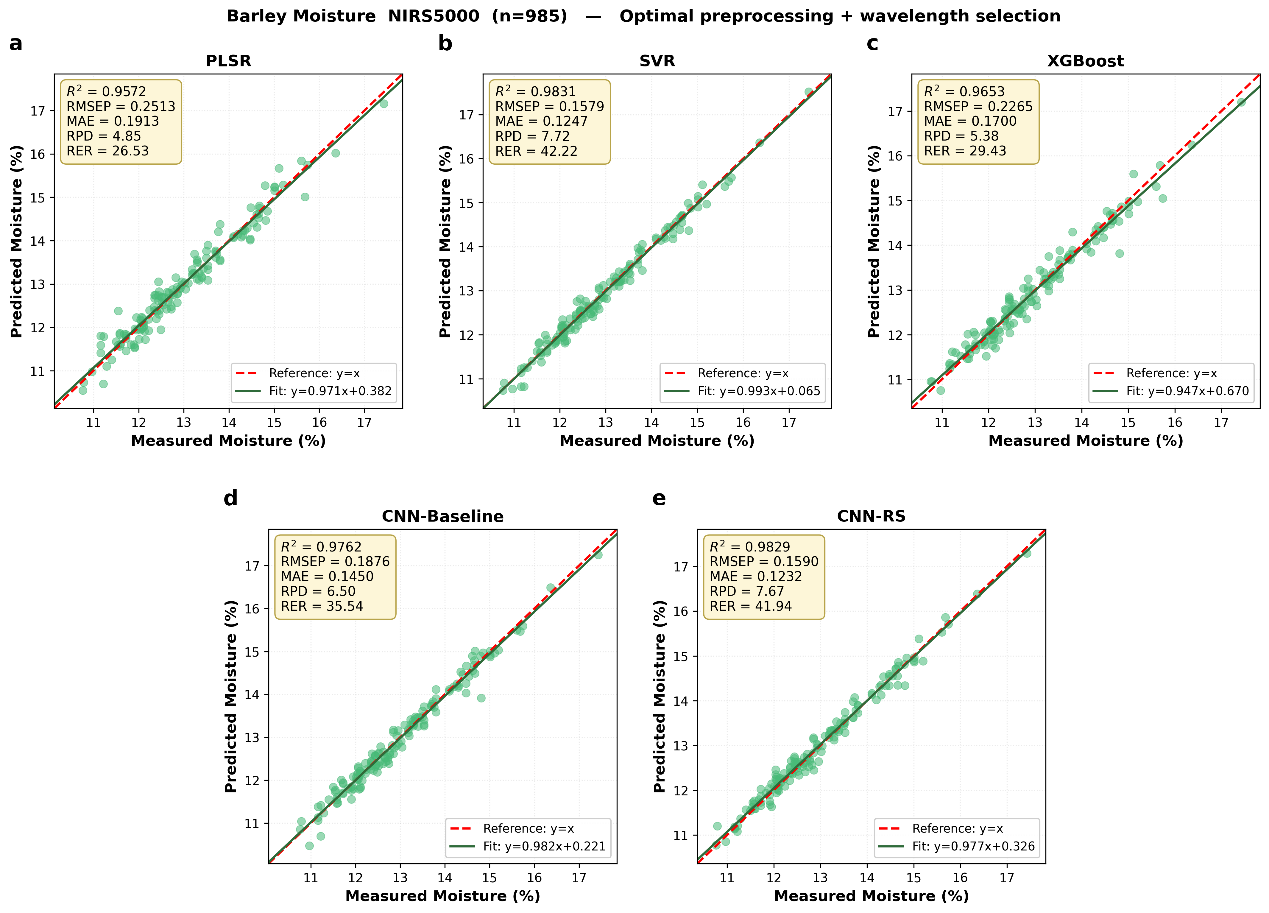

**Figure S11.** Measured versus predicted values of the five models on barley moisture (NIRS5000, n=985) with both optimal preprocessing and wavelength selection: SVR, XGBoost and both CNNs share the wavelength subset derived from the PLSR regression coefficients for this dataset, shown at their representative seeds. Test-set samples are shown in green; all other conventions are as in Figure 6.

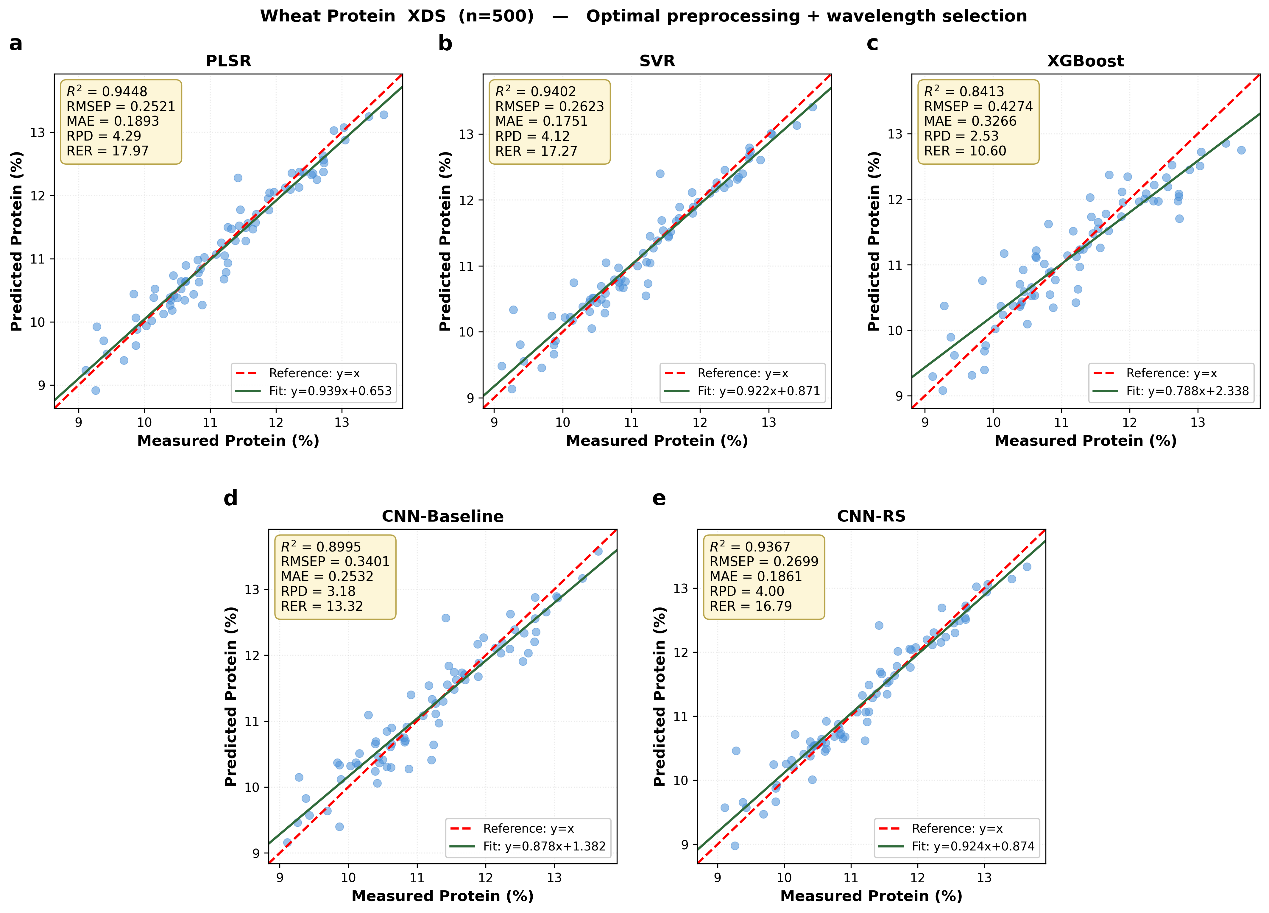

**Figure S12.** Measured versus predicted values of the five models on wheat protein (XDS, n=500) with both optimal preprocessing and wavelength selection: SVR, XGBoost and both CNNs share the wavelength subset derived from the PLSR regression coefficients for this dataset, shown at their representative seeds. Test-set samples are shown in blue; all other conventions are as in Figure 6.

**Table S1.** Comparison of spectral preprocessing methods for PLSR models across the six datasets.

**(a) Wheat protein (NIRS5000, n = 5046)**

| **Preprocessing** | **LVs** | **Train R²** | **Val R²** | **Test R²** | **RMSEP** | **MAE** | **RPD** | **RER** |
| --- | --- | --- | --- | --- | --- | --- | --- | --- |
| **snv-sd** | **20** | **0.9607** | **0.9528** | **0.9574** | **0.2926** | **0.2318** | **4.85** | **47.21** |
| sd-snv | 20 | 0.9585 | 0.9497 | 0.9560 | 0.2975 | 0.2367 | 4.77 | 46.44 |
| sd | 20 | 0.9564 | 0.9484 | 0.9530 | 0.3074 | 0.2463 | 4.61 | 44.94 |
| snv-fd | 20 | 0.9538 | 0.9448 | 0.9552 | 0.3002 | 0.2365 | 4.72 | 46.02 |
| fd | 20 | 0.9493 | 0.9404 | 0.9526 | 0.3088 | 0.2444 | 4.59 | 44.74 |
| fd-snv | 20 | 0.9511 | 0.9396 | 0.9536 | 0.3055 | 0.2419 | 4.64 | 45.21 |
| snv-detrend | 20 | 0.9460 | 0.9354 | 0.9486 | 0.3216 | 0.2572 | 4.41 | 42.95 |
| msc | 20 | 0.9445 | 0.9319 | 0.9473 | 0.3255 | 0.2569 | 4.36 | 42.44 |
| snv | 20 | 0.9446 | 0.9314 | 0.9470 | 0.3265 | 0.2564 | 4.34 | 42.31 |
| detrend | 20 | 0.9386 | 0.9268 | 0.9428 | 0.3391 | 0.2703 | 4.18 | 40.74 |
| none | 20 | 0.9362 | 0.9261 | 0.9438 | 0.3363 | 0.2662 | 4.22 | 41.08 |
| baseline | 20 | 0.9294 | 0.9187 | 0.9354 | 0.3603 | 0.2898 | 3.94 | 38.34 |

**(b) Barley protein (NIRS5000, n = 2096)**

| **Preprocessing** | **LVs** | **Train R²** | **Val R²** | **Test R²** | **RMSEP** | **MAE** | **RPD** | **RER** |
| --- | --- | --- | --- | --- | --- | --- | --- | --- |
| **sd** | **20** | **0.9599** | **0.9529** | **0.9544** | **0.3395** | **0.2690** | **4.68** | **26.54** |
| snv-sd | 19 | 0.9588 | 0.9500 | 0.9552 | 0.3368 | 0.2654 | 4.72 | 26.75 |
| fd | 20 | 0.9499 | 0.9478 | 0.9484 | 0.3613 | 0.2805 | 4.40 | 24.94 |
| sd-snv | 19 | 0.9559 | 0.9477 | 0.9513 | 0.3510 | 0.2789 | 4.53 | 25.67 |
| snv-fd | 20 | 0.9503 | 0.9436 | 0.9457 | 0.3704 | 0.2881 | 4.29 | 24.32 |
| fd-snv | 20 | 0.9484 | 0.9421 | 0.9444 | 0.3749 | 0.2927 | 4.24 | 24.03 |
| detrend | 20 | 0.9409 | 0.9301 | 0.9425 | 0.3813 | 0.3029 | 4.17 | 23.63 |
| snv-detrend | 20 | 0.9414 | 0.9274 | 0.9431 | 0.3795 | 0.3056 | 4.19 | 23.74 |
| baseline | 20 | 0.9321 | 0.9155 | 0.9376 | 0.3974 | 0.3155 | 4.00 | 22.67 |
| none | 20 | 0.9240 | 0.9066 | 0.9252 | 0.4348 | 0.3432 | 3.66 | 20.72 |
| snv | 20 | 0.9225 | 0.8984 | 0.9273 | 0.4288 | 0.3384 | 3.71 | 21.01 |
| msc | 20 | 0.9216 | 0.8970 | 0.9263 | 0.4317 | 0.3434 | 3.68 | 20.87 |

**(c) Wheat protein (XDS, n = 500)**

| **Preprocessing** | **LVs** | **Train R²** | **Val R²** | **Test R²** | **RMSEP** | **MAE** | **RPD** | **RER** |
| --- | --- | --- | --- | --- | --- | --- | --- | --- |
| **snv-fd** | **15** | **0.9555** | **0.9434** | **0.9326** | **0.2787** | **0.2002** | **3.85** | **16.26** |
| fd-snv | 20 | 0.9600 | 0.9425 | 0.9378 | 0.2675 | 0.1995 | 4.01 | 16.93 |
| msc | 20 | 0.9483 | 0.9385 | 0.9316 | 0.2807 | 0.1947 | 3.82 | 16.14 |
| snv-detrend | 20 | 0.9471 | 0.9345 | 0.9253 | 0.2933 | 0.2120 | 3.66 | 15.44 |
| snv | 20 | 0.9467 | 0.9322 | 0.9285 | 0.2869 | 0.2018 | 3.74 | 15.79 |
| fd | 15 | 0.9487 | 0.9267 | 0.9240 | 0.2957 | 0.2180 | 3.63 | 15.32 |
| detrend | 20 | 0.9403 | 0.9265 | 0.9132 | 0.3161 | 0.2343 | 3.39 | 14.33 |
| snv-sd | 14 | 0.9692 | 0.9212 | 0.9372 | 0.2690 | 0.2130 | 3.99 | 16.84 |
| none | 20 | 0.9313 | 0.9202 | 0.9114 | 0.3194 | 0.2407 | 3.36 | 14.18 |
| baseline | 19 | 0.9332 | 0.9154 | 0.9053 | 0.3302 | 0.2435 | 3.25 | 13.72 |
| sd-snv | 15 | 0.9673 | 0.9150 | 0.9288 | 0.2863 | 0.2262 | 3.75 | 15.82 |
| sd | 13 | 0.9628 | 0.9117 | 0.9220 | 0.2996 | 0.2366 | 3.58 | 15.12 |

**(d) Wheat moisture (NIRS5000, n = 3305)**

| **Preprocessing** | **LVs** | **Train R²** | **Val R²** | **Test R²** | **RMSEP** | **MAE** | **RPD** | **RER** |
| --- | --- | --- | --- | --- | --- | --- | --- | --- |
| **snv-sd** | **20** | **0.9529** | **0.9426** | **0.9442** | **0.2927** | **0.2226** | **4.23** | **24.66** |
| snv-fd | 20 | 0.9407 | 0.9385 | 0.9392 | 0.3056 | 0.2339 | 4.06 | 23.63 |
| sd-snv | 20 | 0.9510 | 0.9383 | 0.9403 | 0.3028 | 0.2307 | 4.09 | 23.85 |
| fd-snv | 20 | 0.9384 | 0.9372 | 0.9378 | 0.3092 | 0.2371 | 4.01 | 23.35 |
| snv-detrend | 20 | 0.9353 | 0.9368 | 0.9370 | 0.3110 | 0.2368 | 3.99 | 23.21 |
| sd | 17 | 0.9448 | 0.9365 | 0.9372 | 0.3106 | 0.2320 | 3.99 | 23.24 |
| msc | 20 | 0.9337 | 0.9356 | 0.9371 | 0.3108 | 0.2358 | 3.99 | 23.23 |
| snv | 20 | 0.9333 | 0.9354 | 0.9366 | 0.3122 | 0.2355 | 3.97 | 23.13 |
| fd | 20 | 0.9356 | 0.9319 | 0.9336 | 0.3194 | 0.2439 | 3.88 | 22.60 |
| none | 20 | 0.9302 | 0.9282 | 0.9324 | 0.3223 | 0.2426 | 3.85 | 22.40 |
| detrend | 20 | 0.9302 | 0.9279 | 0.9331 | 0.3206 | 0.2429 | 3.87 | 22.52 |
| baseline | 19 | 0.9239 | 0.9233 | 0.9289 | 0.3306 | 0.2486 | 3.75 | 21.84 |

**(e) Barley moisture (NIRS5000, n = 985)**

| **Preprocessing** | **LVs** | **Train R²** | **Val R²** | **Test R²** | **RMSEP** | **MAE** | **RPD** | **RER** |
| --- | --- | --- | --- | --- | --- | --- | --- | --- |
| **snv-sd** | **19** | **0.9679** | **0.9387** | **0.9598** | **0.2435** | **0.1825** | **4.99** | **27.38** |
| sd-snv | 19 | 0.9681 | 0.9366 | 0.9583 | 0.2480 | 0.1865 | 4.90 | 26.88 |
| sd | 20 | 0.9693 | 0.9357 | 0.9591 | 0.2458 | 0.1897 | 4.94 | 27.13 |
| fd-snv | 20 | 0.9575 | 0.9276 | 0.9554 | 0.2565 | 0.2035 | 4.74 | 25.99 |
| snv-fd | 16 | 0.9488 | 0.9260 | 0.9513 | 0.2681 | 0.2162 | 4.53 | 24.87 |
| baseline | 20 | 0.9408 | 0.9252 | 0.9402 | 0.2971 | 0.2299 | 4.09 | 22.44 |
| snv | 16 | 0.9365 | 0.9252 | 0.9321 | 0.3166 | 0.2501 | 3.84 | 21.06 |
| snv-detrend | 20 | 0.9489 | 0.9240 | 0.9544 | 0.2596 | 0.2084 | 4.68 | 25.68 |
| msc | 16 | 0.9363 | 0.9224 | 0.9312 | 0.3188 | 0.2546 | 3.81 | 20.91 |
| fd | 20 | 0.9569 | 0.9212 | 0.9557 | 0.2558 | 0.2009 | 4.75 | 26.07 |
| none | 19 | 0.9424 | 0.9204 | 0.9416 | 0.2935 | 0.2382 | 4.14 | 22.71 |
| detrend | 15 | 0.9381 | 0.9189 | 0.9309 | 0.3194 | 0.2483 | 3.80 | 20.88 |

**(f) Maize moisture (NIRS5000, n = 585)**

| **Preprocessing** | **LVs** | **Train R²** | **Val R²** | **Test R²** | **RMSEP** | **MAE** | **RPD** | **RER** |
| --- | --- | --- | --- | --- | --- | --- | --- | --- |
| **sd-snv** | **20** | **0.9541** | **0.8933** | **0.8807** | **0.5329** | **0.3988** | **2.90** | **13.70** |
| snv-sd | 20 | 0.9531 | 0.8914 | 0.8769 | 0.5414 | 0.4054 | 2.85 | 13.48 |
| sd | 20 | 0.9397 | 0.8789 | 0.8611 | 0.5752 | 0.4480 | 2.68 | 12.69 |
| fd-snv | 20 | 0.9269 | 0.8608 | 0.8728 | 0.5504 | 0.4099 | 2.80 | 13.26 |
| snv-fd | 20 | 0.9278 | 0.8607 | 0.8695 | 0.5576 | 0.4126 | 2.77 | 13.09 |
| baseline | 20 | 0.8886 | 0.8587 | 0.8186 | 0.6573 | 0.5163 | 2.35 | 11.11 |
| snv-detrend | 20 | 0.9159 | 0.8587 | 0.8404 | 0.6165 | 0.4383 | 2.50 | 11.84 |
| snv | 20 | 0.9104 | 0.8547 | 0.8321 | 0.6323 | 0.4551 | 2.44 | 11.55 |
| msc | 18 | 0.9063 | 0.8542 | 0.8240 | 0.6475 | 0.4613 | 2.38 | 11.27 |
| fd | 19 | 0.9016 | 0.8457 | 0.8480 | 0.6016 | 0.4508 | 2.57 | 12.13 |
| detrend | 20 | 0.8997 | 0.8433 | 0.8260 | 0.6437 | 0.4772 | 2.40 | 11.34 |
| none | 20 | 0.8862 | 0.8412 | 0.8091 | 0.6742 | 0.5148 | 2.29 | 10.83 |

Note: Twelve preprocessing strategies were compared: none (raw spectra), baseline correction, detrend, standard normal variate (SNV), multiplicative scatter correction (MSC), first derivative (FD), second derivative (SD), and their combinations (SNV + SD, SNV + FD, SD + SNV, FD + SNV, SNV + detrend). For each method, the number of PLS latent variables (LVs) and the training, validation and test-set metrics are reported. R^2^: coefficient of determination; RMSEP: root means square error prediction absolute error; RPD: ratio of performance to deviation; RER: range error ratio. Within each dataset, the methods are ranked by the R^2^ value of the validation set; the preprocessing with the best performance (in bold) is carried over to the wavelength selection stage.

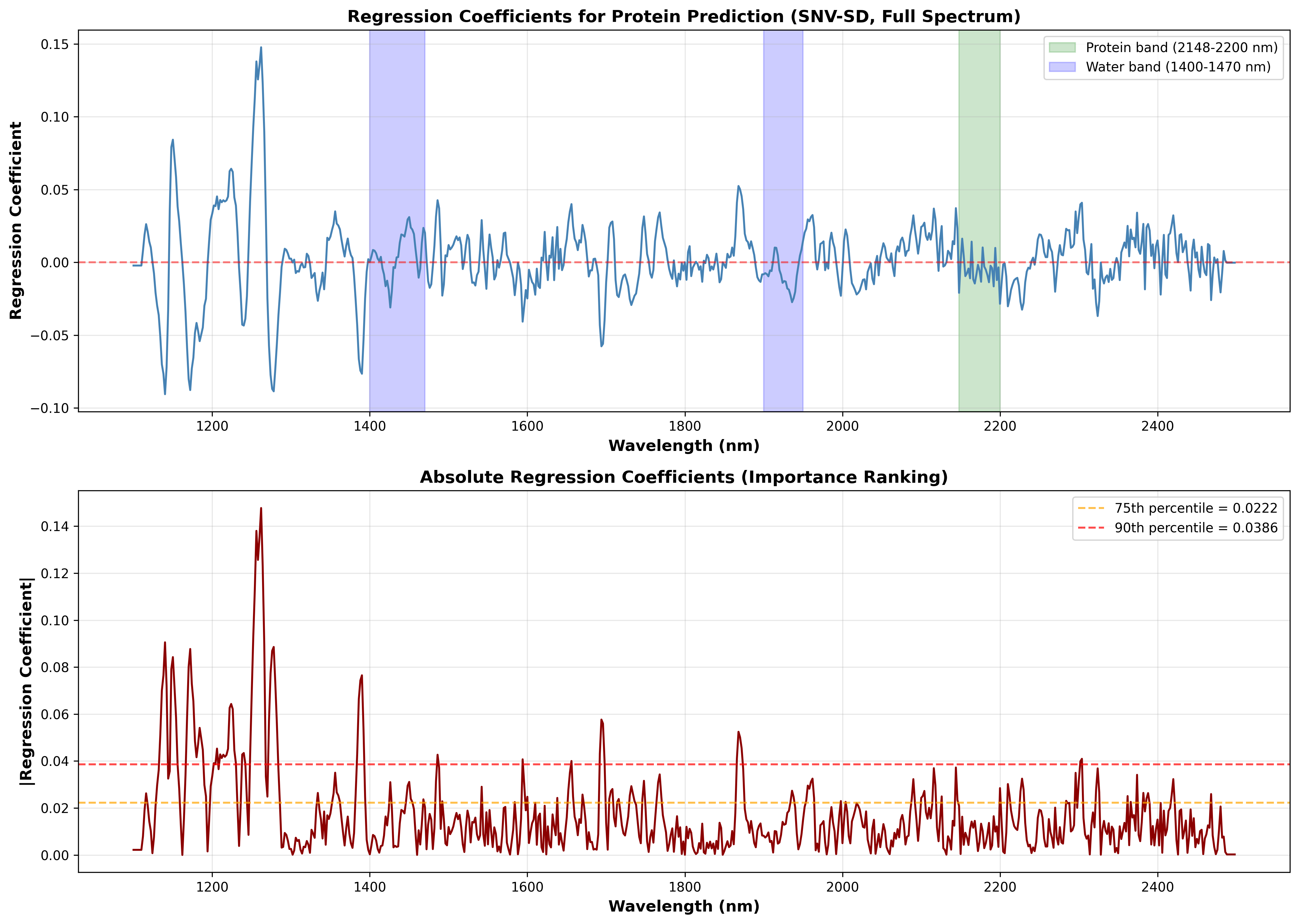

**Figure S13.** Regression Coefficient Analysis (RCA) of the PLSR model for the wheat protein dataset (NIRS5000, n=5046). (Top) Regression coefficients for full spectrum (1100-2498 nm) after SNV and SD preprocessing. Positive and negative coefficients indicate a positive or negative correlation between wavelength and protein content, respectively. The red dashed line represents zero. The shaded areas indicate characteristic absorption bands (water, 1400-1470 nm, 1900-1950 nm; protein, 2148-2200 nm). (Bottom) Absolute regression coefficients were used for importance ranking. The dashed lines mark the 75^th^ (0.0222) and 90^th^ (0.0386) percentile thresholds, which define the importance-based wavelength selection strategies evaluated in Table S2 (e.g., Top 25%, Top 10%). The results were shown only for this representative dataset and same analysis was performed on all six datasets.

**Table S2.** Comparison of wavelength-selection strategies for PLSR models across six datasets.

**(a) Wheat protein (NIRS5000, n = 5046)**

| **Strategy** | **n_wl** | **Range (nm)** | **LVs** | **Val R²** | **Test R²** | **RMSEP** | **MAE** | **RPD** | **RER** |
| --- | --- | --- | --- | --- | --- | --- | --- | --- | --- |
| **Top_25%_Important** | **175** | **1116-2468** | **20** | **0.9534** | **0.9589** | **0.2876** | **0.2288** | **4.93** | **48.03** |
| Full_Spectrum | 700 | 1100-2498 | 20 | 0.9528 | 0.9574 | 0.2926 | 0.2318 | 4.85 | 47.21 |
| Top_20%_Important | 140 | 1116-2468 | 20 | 0.9528 | 0.9577 | 0.2918 | 0.2310 | 4.86 | 47.35 |
| Peaks_Plus_Protein | 601 | 1110-2498 | 18 | 0.9528 | 0.9569 | 0.2943 | 0.2335 | 4.82 | 46.95 |
| Peak_Regions_75pct | 590 | 1110-2498 | 19 | 0.9526 | 0.9569 | 0.2945 | 0.2337 | 4.81 | 46.91 |
| Top20%_Plus_Protein | 205 | 1116-2468 | 20 | 0.9524 | 0.9569 | 0.2945 | 0.2351 | 4.81 | 46.91 |
| Top_15%_Important | 105 | 1132-2420 | 20 | 0.9496 | 0.9549 | 0.3011 | 0.2384 | 4.71 | 45.88 |
| Top_10%_Important | 70 | 1134-2304 | 18 | 0.9406 | 0.9483 | 0.3225 | 0.2519 | 4.40 | 42.84 |
| Top_5%_Important | 35 | 1136-1694 | 20 | 0.9322 | 0.9371 | 0.3558 | 0.2750 | 3.99 | 38.83 |
| Extended_1400-2500 | 550 | 1400-2498 | 19 | 0.9165 | 0.9433 | 0.3376 | 0.2694 | 4.20 | 40.92 |
| Protein_Region_1900-2300 | 201 | 1900-2300 | 16 | 0.7483 | 0.7891 | 0.6511 | 0.4920 | 2.18 | 21.22 |
| Protein_Region_2100-2250 | 76 | 2100-2250 | 17 | 0.5669 | 0.5884 | 0.9097 | 0.6780 | 1.56 | 15.19 |
| Protein_Band_2148-2200 | 27 | 2148-2200 | 17 | 0.4424 | 0.4685 | 1.0338 | 0.7839 | 1.37 | 13.36 |

**(b) Barley protein (NIRS5000, n = 2096)**

| **Strategy** | **n_wl** | **Range (nm)** | **LVs** | **Val R²** | **Test R²** | **RMSEP** | **MAE** | **RPD** | **RER** |
| --- | --- | --- | --- | --- | --- | --- | --- | --- | --- |
| **Full_Spectrum** | **700** | **1100-2498** | **20** | **0.9529** | **0.9544** | **0.3395** | **0.2690** | **4.68** | **26.54** |
| Peaks_Plus_Protein | 637 | 1110-2494 | 20 | 0.9524 | 0.9550 | 0.3373 | 0.2689 | 4.72 | 26.71 |
| Peak_Regions_75pct | 592 | 1110-2494 | 20 | 0.9509 | 0.9545 | 0.3393 | 0.2696 | 4.69 | 26.56 |
| Top_20%_Important | 140 | 1124-2480 | 20 | 0.9505 | 0.9489 | 0.3595 | 0.2883 | 4.42 | 25.06 |
| Top_25%_Important | 175 | 1124-2480 | 20 | 0.9499 | 0.9539 | 0.3416 | 0.2704 | 4.66 | 26.37 |
| Top20%_Plus_Protein | 209 | 1124-2480 | 20 | 0.9498 | 0.9523 | 0.3473 | 0.2781 | 4.58 | 25.94 |
| Top_15%_Important | 105 | 1126-2464 | 20 | 0.9424 | 0.9435 | 0.3781 | 0.2993 | 4.21 | 23.83 |
| Top_10%_Important | 70 | 1132-2420 | 20 | 0.9328 | 0.9461 | 0.3692 | 0.2877 | 4.31 | 24.40 |
| Top_5%_Important | 35 | 1134-1820 | 20 | 0.9123 | 0.9243 | 0.4376 | 0.3430 | 3.63 | 20.59 |
| Extended_1400-2500 | 550 | 1400-2498 | 20 | 0.8846 | 0.8945 | 0.5166 | 0.4093 | 3.08 | 17.44 |
| Protein_Region_1900-2300 | 201 | 1900-2300 | 13 | 0.6325 | 0.7423 | 0.8073 | 0.6457 | 1.97 | 11.16 |
| Protein_Region_2100-2250 | 76 | 2100-2250 | 12 | 0.3571 | 0.3656 | 1.2667 | 1.0070 | 1.26 | 7.11 |
| Protein_Band_2148-2200 | 27 | 2148-2200 | 14 | 0.2616 | 0.2888 | 1.3412 | 1.0425 | 1.19 | 6.72 |

**(c) Wheat protein (XDS, n = 500)**

| **Strategy** | **n_wl** | **Range (nm)** | **LVs** | **Val R²** | **Test R²** | **RMSEP** | **MAE** | **RPD** | **RER** |
| --- | --- | --- | --- | --- | --- | --- | --- | --- | --- |
| **Top20%_Plus_Protein** | **272** | **682-2474** | **20** | **0.9459** | **0.9448** | **0.2521** | **0.1893** | **4.26** | **17.97** |
| Top_10%_Important | 105 | 962-2324 | 18 | 0.9448 | 0.9418 | 0.2588 | 0.1988 | 4.15 | 17.50 |
| Top_20%_Important | 210 | 682-2474 | 19 | 0.9445 | 0.9466 | 0.2480 | 0.1886 | 4.33 | 18.27 |
| Full_Spectrum | 1050 | 400-2498 | 15 | 0.9434 | 0.9326 | 0.2787 | 0.2002 | 3.85 | 16.26 |
| Peak_Regions_75pct | 706 | 656-2498 | 13 | 0.9426 | 0.9392 | 0.2646 | 0.1993 | 4.05 | 17.12 |
| Peaks_Plus_Protein | 706 | 656-2498 | 13 | 0.9426 | 0.9392 | 0.2646 | 0.1993 | 4.05 | 17.12 |
| Top_25%_Important | 263 | 662-2474 | 12 | 0.9424 | 0.9415 | 0.2596 | 0.1934 | 4.13 | 17.45 |
| Top_15%_Important | 158 | 686-2356 | 20 | 0.9422 | 0.9457 | 0.2500 | 0.1883 | 4.29 | 18.12 |
| Top_5%_Important | 53 | 998-2322 | 16 | 0.9416 | 0.9286 | 0.2867 | 0.2184 | 3.74 | 15.80 |
| Extended_1400-2500 | 550 | 1400-2498 | 15 | 0.9087 | 0.9136 | 0.3154 | 0.2237 | 3.40 | 14.36 |
| Protein_Region_1900-2300 | 201 | 1900-2300 | 10 | 0.8315 | 0.7990 | 0.4810 | 0.3746 | 2.23 | 9.42 |
| Protein_Region_2100-2250 | 76 | 2100-2250 | 8 | 0.7540 | 0.7227 | 0.5650 | 0.4450 | 1.90 | 8.02 |
| Protein_Band_2148-2200 | 27 | 2148-2200 | 18 | 0.6547 | 0.6510 | 0.6339 | 0.4911 | 1.69 | 7.15 |

**(d) Wheat moisture (NIRS5000, n = 3305)**

| **Strategy** | **n_wl** | **Range (nm)** | **LVs** | **Val R²** | **Test R²** | **RMSEP** | **MAE** | **RPD** | **RER** |
| --- | --- | --- | --- | --- | --- | --- | --- | --- | --- |
| **Top20%_Plus_Moisture** | **192** | **1124-2470** | **20** | **0.9436** | **0.9448** | **0.2913** | **0.2206** | **4.26** | **24.78** |
| Top_25%_Important | 175 | 1122-2478 | 20 | 0.9435 | 0.9451 | 0.2905 | 0.2218 | 4.27 | 24.85 |
| Top_20%_Important | 140 | 1124-2470 | 20 | 0.9431 | 0.9448 | 0.2913 | 0.2213 | 4.26 | 24.79 |
| Peaks_Plus_Moisture | 670 | 1106-2498 | 20 | 0.9426 | 0.9436 | 0.2944 | 0.2239 | 4.21 | 24.53 |
| Full_Spectrum | 700 | 1100-2498 | 20 | 0.9426 | 0.9442 | 0.2927 | 0.2226 | 4.23 | 24.66 |
| Peak_Regions_75pct | 658 | 1106-2498 | 20 | 0.9425 | 0.9434 | 0.2949 | 0.2249 | 4.20 | 24.48 |
| Extended_1350-2400 | 526 | 1350-2400 | 20 | 0.9416 | 0.9422 | 0.2981 | 0.2267 | 4.16 | 24.22 |
| Top_15%_Important | 105 | 1124-2470 | 20 | 0.9410 | 0.9443 | 0.2926 | 0.2223 | 4.24 | 24.68 |
| Top_10%_Important | 70 | 1124-2470 | 18 | 0.9394 | 0.9389 | 0.3064 | 0.2305 | 4.05 | 23.56 |
| Top_5%_Important | 35 | 1136-2470 | 19 | 0.9258 | 0.9289 | 0.3305 | 0.2530 | 3.75 | 21.85 |
| All_Moisture_Regions | 163 | 1400-2300 | 20 | 0.9111 | 0.9101 | 0.3716 | 0.2792 | 3.34 | 19.43 |
| Primary_Moisture_Bands | 62 | 1400-1950 | 16 | 0.8976 | 0.9082 | 0.3755 | 0.2815 | 3.30 | 19.23 |
| Water_Band_1400-1470 | 36 | 1400-1470 | 19 | 0.8785 | 0.8813 | 0.4271 | 0.3225 | 2.90 | 16.91 |
| Water_Band_1900-1950 | 26 | 1900-1950 | 19 | 0.7780 | 0.7972 | 0.5582 | 0.4461 | 2.22 | 12.93 |
| Water_Band_2100-2300 | 101 | 2100-2300 | 20 | 0.6952 | 0.6922 | 0.6877 | 0.5375 | 1.80 | 10.50 |

**(e) Barley moisture (NIRS5000, n = 985)**

| **Strategy** | **n_wl** | **Range (nm)** | **LVs** | **Val R²** | **Test R²** | **RMSEP** | **MAE** | **RPD** | **RER** |
| --- | --- | --- | --- | --- | --- | --- | --- | --- | --- |
| **Top20%_Plus_Moisture** | **196** | **1124-2480** | **20** | **0.9427** | **0.9572** | **0.2513** | **0.1913** | **4.83** | **26.53** |
| Peak_Regions_75pct | 596 | 1110-2498 | 19 | 0.9422 | 0.9604 | 0.2418 | 0.1863 | 5.03 | 27.57 |
| Top_20%_Important | 140 | 1124-2480 | 20 | 0.9416 | 0.9572 | 0.2513 | 0.1907 | 4.84 | 26.53 |
| Peaks_Plus_Moisture | 618 | 1110-2498 | 20 | 0.9409 | 0.9603 | 0.2420 | 0.1863 | 5.02 | 27.55 |
| Top_15%_Important | 105 | 1136-2480 | 19 | 0.9406 | 0.9529 | 0.2638 | 0.2037 | 4.61 | 25.27 |
| Top_25%_Important | 175 | 1124-2480 | 20 | 0.9404 | 0.9588 | 0.2467 | 0.1890 | 4.93 | 27.03 |
| Full_Spectrum | 700 | 1100-2498 | 19 | 0.9387 | 0.9598 | 0.2435 | 0.1825 | 4.99 | 27.38 |
| Top_10%_Important | 70 | 1136-2418 | 20 | 0.9346 | 0.9452 | 0.2846 | 0.2166 | 4.27 | 23.43 |
| Extended_1350-2400 | 526 | 1350-2400 | 19 | 0.9309 | 0.9574 | 0.2507 | 0.1926 | 4.85 | 26.60 |
| All_Moisture_Regions | 163 | 1400-2300 | 16 | 0.9150 | 0.9173 | 0.3494 | 0.2766 | 3.48 | 19.09 |
| Top_5%_Important | 35 | 1140-2400 | 17 | 0.9109 | 0.9107 | 0.3631 | 0.2691 | 3.35 | 18.36 |
| Primary_Moisture_Bands | 62 | 1400-1950 | 19 | 0.8903 | 0.8975 | 0.3890 | 0.3084 | 3.12 | 17.14 |
| Water_Band_1400-1470 | 36 | 1400-1470 | 15 | 0.8630 | 0.8753 | 0.4291 | 0.3467 | 2.83 | 15.54 |
| Water_Band_2100-2300 | 101 | 2100-2300 | 17 | 0.8425 | 0.8700 | 0.4381 | 0.3538 | 2.77 | 15.22 |
| Water_Band_1900-1950 | 26 | 1900-1950 | 14 | 0.8032 | 0.8010 | 0.5420 | 0.4243 | 2.24 | 12.30 |

**(f) Maize moisture (NIRS5000, n = 585)**

| **Strategy** | **n_wl** | **Range (nm)** | **LVs** | **Val R²** | **Test R²** | **RMSEP** | **MAE** | **RPD** | **RER** |
| --- | --- | --- | --- | --- | --- | --- | --- | --- | --- |
| **Peaks_Plus_Moisture** | **621** | **1140-2498** | **20** | **0.8936** | **0.8731** | **0.5498** | **0.4222** | **2.81** | **13.28** |
| Full_Spectrum | 700 | 1100-2498 | 20 | 0.8933 | 0.8807 | 0.5329 | 0.3988 | 2.90 | 13.70 |
| Peak_Regions_75pct | 605 | 1140-2498 | 19 | 0.8930 | 0.8744 | 0.5468 | 0.4224 | 2.82 | 13.35 |
| Top_25%_Important | 175 | 1164-2480 | 19 | 0.8915 | 0.8823 | 0.5293 | 0.4098 | 2.92 | 13.79 |
| Top20%_Plus_Moisture | 178 | 1164-2480 | 20 | 0.8888 | 0.8781 | 0.5388 | 0.4106 | 2.86 | 13.55 |
| Top_20%_Important | 140 | 1164-2480 | 20 | 0.8886 | 0.8759 | 0.5436 | 0.4184 | 2.84 | 13.43 |
| Extended_1350-2400 | 526 | 1350-2400 | 20 | 0.8854 | 0.8754 | 0.5447 | 0.4224 | 2.83 | 13.40 |
| Top_10%_Important | 70 | 1166-2480 | 17 | 0.8781 | 0.8741 | 0.5476 | 0.3923 | 2.82 | 13.33 |
| Top_15%_Important | 105 | 1164-2480 | 20 | 0.8767 | 0.8756 | 0.5444 | 0.4034 | 2.83 | 13.41 |
| Primary_Moisture_Bands | 62 | 1400-1950 | 20 | 0.8748 | 0.9013 | 0.4848 | 0.3662 | 3.18 | 15.06 |
| Top_5%_Important | 35 | 1166-2480 | 20 | 0.8622 | 0.8597 | 0.5780 | 0.4041 | 2.67 | 12.63 |
| All_Moisture_Regions | 163 | 1400-2300 | 18 | 0.8587 | 0.8777 | 0.5396 | 0.4140 | 2.86 | 13.53 |
| Water_Band_1400-1470 | 36 | 1400-1470 | 20 | 0.8513 | 0.8699 | 0.5566 | 0.3964 | 2.77 | 13.12 |
| Water_Band_1900-1950 | 26 | 1900-1950 | 12 | 0.8167 | 0.8016 | 0.6874 | 0.5074 | 2.25 | 10.62 |
| Water_Band_2100-2300 | 101 | 2100-2300 | 18 | 0.7403 | 0.8034 | 0.6843 | 0.5508 | 2.26 | 10.67 |

Note: For each dataset, the wavelength selection strategies were evaluated using the optimal preprocessing methods specified in Table S1. The strategies included Full Spectrum; importance-based selection, retaining the top 5% to 25% of wavelengths ranked by absolute PLS regression coefficients (Top_X%_Important; see figure S2); and bandwidth selection based on chemical information, targeting regions of absorption associated with proteins, water or water-related with compounds. n_wl denote the number of retained wavelengths and LVs denote the number of PLS latent variables. R^2^, coefficient of determination; RMSEP, root mean square error; MAE, mean absolute error; RPD, the ratio of performance to bias; RER, the range error ratio. In each dataset, the strategies were ranked by their R^2^ values on the validation set, and the final PLSR model employs the strategy with the best performance (bolded).

**Table S3.** The performance of support vector regression (SVR) on test set across six datasets under three input conditions.

|  | **Input** | **Preprocessing** | **Wave-**  **lengths** | **Test R²** | **Test**  **RMSEP** | **Test**  **MAE** | **Test**  **RPD** | **Test**  **RER** |
| --- | --- | --- | --- | --- | --- | --- | --- | --- |
| (a) Wheat protein (NIRS5000, n = 5046) | Raw | none | 700 | 0.9526 ± 0.0005 | 0.3086 ± 0.0016 | 0.2417 ± 0.0008 | 4.59 ± 0.02 | 44.76 ± 0.23 |
|  | Preprocessed | SD | 700 | 0.9655 ± 0.0001 | 0.2634 ± 0.0004 | 0.2073 ± 0.0012 | 5.38 ± 0.01 | 52.44 ± 0.09 |
|  | **Preprocessed + wavelength subset** | **SD** | **175** | **0.9680 ± 0.0003** | **0.2538 ± 0.0010** | **0.1984 ± 0.0005** | **5.59 ± 0.02** | **54.44 ± 0.22** |
| (b) Barley protein (NIRS5000, n = 2096) | Raw | none | 700 | 0.9093 ± 0.0098 | 0.4784 ± 0.0268 | 0.3652 ± 0.0344 | 3.34 ± 0.20 | 18.90 ± 1.13 |
|  | **Preprocessed** | **SNV-SD** | **700** | **0.9662 ± 0.0021** | **0.2921 ± 0.0090** | **0.2265 ± 0.0024** | **5.45 ± 0.16** | **30.88 ± 0.91** |
|  | Preprocessed + wavelength subset | SNV-SD | 700 | 0.9662 ± 0.0021 | 0.2921 ± 0.0090 | 0.2265 ± 0.0024 | 5.45 ± 0.16 | 30.88 ± 0.91 |
| (c) Wheat protein (XDS, n = 500) | Raw | none | 1050 | 0.8526 ± 0.0035 | 0.4119 ± 0.0050 | 0.3230 ± 0.0050 | 2.61 ± 0.03 | 11.00 ± 0.13 |
|  | Preprocessed | SNV-FD | 1050 | 0.9412 ± 0.0027 | 0.2601 ± 0.0059 | 0.1719 ± 0.0069 | 4.13 ± 0.09 | 17.43 ± 0.38 |
|  | **Preprocessed + wavelength subset** | **SNV-FD** | **272** | **0.9415 ± 0.0028** | **0.2595 ± 0.0063** | **0.1771 ± 0.0073** | **4.14 ± 0.11** | **17.46 ± 0.46** |
| (d) Wheat moisture (NIRS5000, n = 3305) | Raw | none | 700 | 0.9500 ± 0.0012 | 0.2771 ± 0.0033 | 0.2015 ± 0.0041 | 4.47 ± 0.05 | 26.06 ± 0.31 |
|  | **Preprocessed** | **FD-SNV** | **700** | **0.9640 ± 0.0009** | **0.2351 ± 0.0028** | **0.1728 ± 0.0017** | **5.27 ± 0.06** | **30.71 ± 0.35** |
|  | Preprocessed + wavelength subset | FD-SNV | 192 | 0.9640 ± 0.0007 | 0.2352 ± 0.0024 | 0.1728 ± 0.0032 | 5.27 ± 0.05 | 30.70 ± 0.31 |
| (e) Barley moisture (NIRS5000, n = 985) | Raw | none | 700 | 0.9667 ± 0.0011 | 0.2216 ± 0.0036 | 0.1711 ± 0.0028 | 5.48 ± 0.09 | 30.09 ± 0.47 |
|  | Preprocessed | SNV-FD | 700 | 0.9823 ± 0.0005 | 0.1615 ± 0.0022 | 0.1256 ± 0.0018 | 7.53 ± 0.10 | 41.30 ± 0.56 |
|  | **Preprocessed + wavelength subset** | **SNV-FD** | **196** | **0.9825 ± 0.0017** | **0.1607 ± 0.0077** | **0.1245 ± 0.0034** | **7.58 ± 0.35** | **41.59 ± 1.93** |
| (f) Maize moisture (NIRS5000, n = 585) | Raw | none | 700 | 0.7433 ± 0.0083 | 0.7818 ± 0.0122 | 0.5316 ± 0.0124 | 1.97 ± 0.03 | 9.34 ± 0.14 |
|  | **Preprocessed** | **SD-SNV** | **700** | **0.8791 ± 0.0051** | **0.5364 ± 0.0111** | **0.3681 ± 0.0028** | **2.88 ± 0.06** | **13.61 ± 0.28** |
|  | Preprocessed + wavelength subset | SD-SNV | 621 | 0.8771 ± 0.0032 | 0.5410 ± 0.0070 | 0.3706 ± 0.0014 | 2.85 ± 0.04 | 13.50 ± 0.17 |

Note: Values are mean ± standard deviation over 25 seeds (42-66), which vary only the cross-validation fold assignment within each hyperparameter search; SVR is otherwise deterministic, and the data partition (70/15/15) is held fixed. Three inputs are compared: raw spectra with column-wise standardization only; the optimal preprocessing applied to the full spectrum; and the same preprocessing combined with wavelength selection. For the preprocessed input, it is selected independently for each dataset by SVR itself, using a grid search ranked by validation-set R^2^; the retained strategy is listed in the Preprocessing column. The wavelength subset is the PLSR validation-selected subset for that dataset (Table S2) and is the same across all models. Hyperparameters are re-optimized on each input (grid search over 384 combinations, five-fold cross-validation on the training set). The highest Test R^2^ is shown in bold. R^2^, coefficient of determination; RMSEP, root mean square error of prediction; MAE, mean absolute error; RPD, the ratio of performance to bias; RER, the range error ratio.

**Table S4.** The performance of extreme gradient boosting (XGBoost) on test set across six datasets under three input conditions.

| **Dataset** | **Input** | **Preprocessing** | **Wave-**  **lengths** | **Test R²** | **Test**  **RMSEP** | **Test**  **MAE** | **Test**  **RPD** | **Test**  **RER** |
| --- | --- | --- | --- | --- | --- | --- | --- | --- |
| (a) Wheat protein (NIRS5000, n = 5046) | Raw | none | 700 | 0.6402 ± 0.0060 | 0.8506 ± 0.0071 | 0.6163 ± 0.0068 | 1.67 ± 0.01 | 16.24 ± 0.14 |
|  | Preprocessed | SD-SNV | 700 | 0.9497 ± 0.0021 | 0.3181 ± 0.0066 | 0.2412 ± 0.0051 | 4.46 ± 0.09 | 43.45 ± 0.90 |
|  | **Preprocessed + wavelength subset** | **SD-SNV** | **175** | **0.9508 ± 0.0016** | **0.3144 ± 0.0052** | **0.2369 ± 0.0045** | **4.51 ± 0.07** | **43.95 ± 0.73** |
| (b) Barley protein (NIRS5000, n = 2096) | Raw | none | 700 | 0.4131 ± 0.0150 | 1.2183 ± 0.0156 | 0.8764 ± 0.0122 | 1.31 ± 0.02 | 7.40 ± 0.10 |
|  | **Preprocessed** | **SNV-SD** | **700** | **0.9416 ± 0.0031** | **0.3841 ± 0.0103** | **0.2910 ± 0.0081** | **4.14 ± 0.11** | **23.47 ± 0.64** |
|  | Preprocessed + wavelength subset | SNV-SD | 700 | 0.9416 ± 0.0031 | 0.3841 ± 0.0103 | 0.2910 ± 0.0081 | 4.14 ± 0.11 | 23.47 ± 0.64 |
| (c) Wheat protein (XDS, n = 500) | Raw | none | 1050 | 0.2467 ± 0.0145 | 0.9312 ± 0.0089 | 0.7259 ± 0.0100 | 1.15 ± 0.01 | 4.86 ± 0.05 |
|  | **Preprocessed** | **SNV-SD** | **1050** | **0.8666 ± 0.0068** | **0.3917 ± 0.0099** | **0.3013 ± 0.0084** | **2.74 ± 0.07** | **11.57 ± 0.29** |
|  | Preprocessed + wavelength subset | SNV-SD | 272 | 0.8416 ± 0.0078 | 0.4269 ± 0.0105 | 0.3300 ± 0.0099 | 2.51 ± 0.06 | 10.62 ± 0.26 |
| (d) Wheat moisture (NIRS5000, n = 3305) | Raw | none | 700 | 0.8475 ± 0.0073 | 0.4840 ± 0.0115 | 0.3632 ± 0.0093 | 2.56 ± 0.06 | 14.93 ± 0.35 |
|  | **Preprocessed** | **SNV-SD** | **700** | **0.9552 ± 0.0015** | **0.2624 ± 0.0045** | **0.1924 ± 0.0038** | **4.73 ± 0.08** | **27.52 ± 0.47** |
|  | Preprocessed + wavelength subset | SNV-SD | 192 | 0.9539 ± 0.0014 | 0.2660 ± 0.0042 | 0.1945 ± 0.0034 | 4.66 ± 0.07 | 27.15 ± 0.43 |
| (e) Barley moisture (NIRS5000, n = 985) | Raw | none | 700 | 0.8596 ± 0.0064 | 0.4551 ± 0.0104 | 0.3371 ± 0.0087 | 2.67 ± 0.06 | 14.66 ± 0.34 |
|  | Preprocessed | FD | 700 | 0.9631 ± 0.0042 | 0.2332 ± 0.0132 | 0.1694 ± 0.0092 | 5.23 ± 0.29 | 28.68 ± 1.59 |
|  | **Preprocessed + wavelength subset** | **FD** | **196** | **0.9653 ± 0.0024** | **0.2264 ± 0.0078** | **0.1667 ± 0.0069** | **5.37 ± 0.18** | **29.49 ± 0.99** |
| (f) Maize moisture (NIRS5000, n = 585) | Raw | none | 700 | 0.5491 ± 0.0260 | 1.0358 ± 0.0301 | 0.7523 ± 0.0212 | 1.49 ± 0.04 | 7.05 ± 0.21 |
|  | Preprocessed | SD | 700 | **0.8207 ± 0.0170** | **0.6529 ± 0.0300** | **0.4366 ± 0.0246** | **2.37 ± 0.10** | **11.20 ± 0.48** |
|  | Preprocessed + wavelength subset | SD | 621 | 0.8179 ± 0.0162 | 0.6580 ± 0.0286 | 0.4368 ± 0.0210 | 2.35 ± 0.10 | 11.11 ± 0.46 |

Note: Values are mean ± standard deviation over 25 seeds (42-66), which vary the sampled hyperparameter configurations and the model’s internal row and column subsampling; the cross-validation folds and the data partition are held fixed. Three inputs are compared: raw spectra with column-wise standardization only; the optimal preprocessing applied to the full spectrum; and same preprocessing combined with wavelength selection. The preprocessing was selected independently for each dataset by XGBoost itself, using a fixed screening configuration evaluated over three seeds and ranked by validation-set R^2^. The wavelength subset is the PLSR validation-selected subset for that dataset (Table S2) and is same across all the models. Hyperparameters were re-optimized on each input (randomized search over 150 configurations, five-fold cross-validation on the training set). The highest Test R^2^ is shown in bold. R^2^, coefficient of determination; RMSEP, root mean square error of prediction; MAE, mean absolute error; RPD, the ratio of performance to bias; RER, the range error ratio.

**Table S5.** Ablation studies for the 1D-CNN architecture and training hyperparameters performed on the Wheat Protein (NIRS5000) dataset.

| **(a) Study 1.1 — Number of convolutional layers** | | | | | | |
| --- | --- | --- | --- | --- | --- | --- |
| **Configuration** | **Params** | **Train R²** | **Val R²** | **Test R²** | **Test RMSEP** | **Test RPD** |
| 1 | 404,403 | 0.9525 | 0.9512 | 0.9581 | 0.2902 | 4.89 |
| 2 | 405,731 | 0.9559 | 0.9545 | 0.9596 | 0.2851 | 4.97 |
| **3** | **407,059** | **0.9658** | **0.9566** | **0.9601** | **0.2834** | **5.00** |
| 4 | 408,387 | 0.9661 | 0.9560 | 0.9607 | 0.2811 | 5.04 |
| 5 | 409,715 | 0.9765 | 0.9527 | 0.9546 | 0.3021 | 4.69 |
| 6 | 411,043 | 0.9711 | 0.9505 | 0.9528 | 0.3081 | 4.60 |
| 7 | 412,371 | 0.9752 | 0.9464 | 0.9543 | 0.3031 | 4.68 |
| 8 | 413,699 | 0.9786 | 0.9435 | 0.9498 | 0.3178 | 4.46 |
| 9 | 415,027 | 0.9783 | 0.9463 | 0.9524 | 0.3095 | 4.58 |
| 10 | 416,355 | 0.9792 | 0.9448 | 0.9515 | 0.3122 | 4.54 |
| 11 | 417,683 | 0.9757 | 0.9453 | 0.9495 | 0.3186 | 4.45 |
| **(b) Study 1.2 — Number of filters** | | | | | | |
| **Configuration** | **Params** | **Train R²** | **Val R²** | **Test R²** | **Test RMSEP** | **Test RPD** |
| **1** | **26,283** | **0.9519** | **0.9547** | **0.9599** | **0.2840** | **4.99** |
| 8 | 202,739 | 0.9446 | 0.9508 | 0.9593 | 0.2859 | 4.96 |
| 16 | 404,403 | 0.9491 | 0.9519 | 0.9585 | 0.2887 | 4.91 |
| 32 | 807,731 | 0.9544 | 0.9531 | 0.9580 | 0.2907 | 4.88 |
| 64 | 1,614,387 | 0.9495 | 0.9540 | 0.9551 | 0.3005 | 4.72 |
| 128 | 3,227,699 | 0.9446 | 0.9499 | 0.9563 | 0.2964 | 4.78 |
| 256 | 6,454,323 | 0.9522 | 0.9495 | 0.9550 | 0.3008 | 4.71 |
| **(c) Study 1.3 — Kernel size** | | | | | | |
| **Configuration** | **Params** | **Train R²** | **Val R²** | **Test R²** | **Test RMSEP** | **Test RPD** |
| 3 | 404,371 | 0.9490 | 0.9513 | 0.9564 | 0.2960 | 4.79 |
| 5 | 404,403 | 0.9417 | 0.9474 | 0.9547 | 0.3017 | 4.70 |
| 7 | 404,435 | 0.9487 | 0.9519 | 0.9568 | 0.2947 | 4.81 |
| 9 | 404,467 | 0.9515 | 0.9532 | 0.9590 | 0.2870 | 4.94 |
| **11** | **404,499** | **0.9475** | **0.9545** | **0.9605** | **0.2820** | **5.03** |
| 13 | 404,531 | 0.9466 | 0.9496 | 0.9547 | 0.3017 | 4.70 |
| 15 | 404,563 | 0.9492 | 0.9516 | 0.9576 | 0.2919 | 4.86 |
| 19 | 404,627 | 0.9459 | 0.9504 | 0.9559 | 0.2977 | 4.76 |
| **(d) Study 2.1 — Number of FC layers** | | | | | | |
| **Configuration** | **Params** | **Train R²** | **Val R²** | **Test R²** | **Test RMSEP** | **Test RPD** |
| 1 | 717,185 | 0.9447 | 0.9497 | 0.9575 | 0.2925 | 4.85 |
| 2 | 1,442,561 | 0.9446 | 0.9527 | 0.9600 | 0.2837 | 5.00 |
| 3 | 1,444,673 | 0.9509 | 0.9543 | 0.9609 | 0.2803 | 5.06 |
| 4 | 2,911,809 | 0.9496 | 0.9535 | 0.9591 | 0.2867 | 4.95 |
| 5 DEEP | 2,912,353 | 0.9492 | 0.9526 | 0.9603 | 0.2826 | 5.02 |
| **5 WIDE** | **5,911,617** | **0.9477** | **0.9549** | **0.9609** | **0.2804** | **5.06** |
| **(e) Study 2.2 — FC layer size** | | | | | | |
| **Configuration** | **Params** | **Train R²** | **Val R²** | **Test R²** | **Test RMSEP** | **Test RPD** |
| small | 359,345 | 0.9432 | 0.9455 | 0.9526 | 0.3087 | 4.59 |
| medium | 719,841 | 0.9453 | 0.9495 | 0.9547 | 0.3018 | 4.70 |
| **large** | **1,444,673** | **0.9513** | **0.9567** | **0.9626** | **0.2742** | **5.17** |
| xlarge | 2,909,697 | 0.9441 | 0.9534 | 0.9598 | 0.2843 | 4.99 |
| original | 404,403 | 0.9541 | 0.9542 | 0.9599 | 0.2838 | 5.00 |
| **(f) Study 3.1 — Dropout rate** | | | | | | |
| **Configuration** | **Params** | **Train R²** | **Val R²** | **Test R²** | **Test RMSEP** | **Test RPD** |
| **0.0** | **1,444,673** | **0.9511** | **0.9543** | **0.9600** | **0.2837** | **5.00** |
| 0.1 | 1,444,673 | 0.6592 | 0.9422 | 0.9495 | 0.3187 | 4.45 |
| 0.2 | 1,444,673 | 0.4040 | 0.9339 | 0.9427 | 0.3394 | 4.18 |
| 0.3 | 1,444,673 | 0.1186 | 0.9210 | 0.9303 | 0.3745 | 3.79 |
| 0.4 | 1,444,673 | -0.3616 | 0.8739 | 0.8888 | 0.4728 | 3.00 |
| 0.5 | 1,444,673 | -0.7633 | 0.8764 | 0.8905 | 0.4693 | 3.02 |
| **(g) Study 3.2 — Batch normalization** | | | | | | |
| **Configuration** | **Params** | **Train R²** | **Val R²** | **Test R²** | **Test RMSEP** | **Test RPD** |
| **With BN** | **1,444,673** | **0.9536** | **0.9558** | **0.9601** | **0.2834** | **5.00** |
| Without BN | 1,444,193 | 0.7428 | 0.7310 | 0.7756 | 0.6718 | 2.11 |
| **(h) Study 3.3 — Weight decay** | | | | | | |
| **Configuration** | **Params** | **Train R²** | **Val R²** | **Test R²** | **Test RMSEP** | **Test RPD** |
| 0.0 | 1,444,673 | 0.9725 | 0.9554 | 0.9630 | 0.2728 | 5.20 |
| **0.001** | **1,444,673** | **0.9662** | **0.9591** | **0.9605** | **0.2819** | **5.03** |
| 0.005 | 1,444,673 | 0.9493 | 0.9529 | 0.9595 | 0.2855 | 4.97 |
| 0.01 | 1,444,673 | 0.9274 | 0.9425 | 0.9494 | 0.3191 | 4.44 |
| 0.05 | 1,444,673 | -0.0579 | -0.1671 | 0.0966 | 1.3478 | 1.05 |
| **(i) Study 4.1 — Learning rate** | | | | | | |
| **Configuration** | **Params** | **Train R²** | **Val R²** | **Test R²** | **Test RMSEP** | **Test RPD** |
| 0.0001 | 26,283 | 0.9282 | 0.9490 | 0.9566 | 0.2955 | 4.80 |
| **0.0005** | **26,283** | **0.9422** | **0.9512** | **0.9585** | **0.2889** | **4.91** |
| 0.001 | 26,283 | 0.9340 | 0.9314 | 0.9448 | 0.3331 | 4.26 |
| 0.0025 | 26,283 | 0.9087 | 0.9340 | 0.9461 | 0.3293 | 4.31 |
| 0.005 | 26,283 | 0.9000 | 0.9059 | 0.9282 | 0.3800 | 3.73 |
| 5e-05 | 26,283 | 0.8435 | 0.8727 | 0.8900 | 0.4702 | 3.02 |
| **(j) Study 4.2 — Optimizer** | | | | | | |
| **Configuration** | **Params** | **Train R²** | **Val R²** | **Test R²** | **Test RMSEP** | **Test RPD** |
| **ADAM** | **26,283** | **0.9458** | **0.9498** | **0.9594** | **0.2857** | **4.96** |
| SGD | 26,283 | 0.6044 | 0.6225 | 0.6933 | 0.7853 | 1.81 |
| **(k) Study 5 — Batch size** | | | | | | |
| **Configuration** | **Params** | **Train R²** | **Val R²** | **Test R²** | **Test RMSEP** | **Test RPD** |
| 16 | 26,283 | 0.7778 | 0.9385 | 0.9471 | 0.3261 | 4.35 |
| 32 | 26,283 | 0.8938 | 0.9509 | 0.9571 | 0.2938 | 4.83 |
| 64 | 26,283 | 0.9271 | 0.9536 | 0.9582 | 0.2898 | 4.89 |
| 128 | 26,283 | 0.9149 | 0.9488 | 0.9547 | 0.3017 | 4.70 |
| **256** | **26,283** | **0.9510** | **0.9545** | **0.9610** | **0.2802** | **5.06** |
| 512 | 26,283 | 0.6411 | 0.8040 | 0.8272 | 0.5894 | 2.41 |
| **(l) Study 6.1 — Activation function** | | | | | | |
| **Configuration** | **Params** | **Train R²** | **Val R²** | **Test R²** | **Test RMSEP** | **Test RPD** |
| **ELU** | **26,283** | **0.9566** | **0.9554** | **0.9625** | **0.2747** | **5.16** |
| TANH | 26,283 | 0.9146 | 0.9424 | 0.9232 | 0.3929 | 3.61 |
| RELU | 26,283 | 0.9102 | 0.9370 | 0.9450 | 0.3326 | 4.26 |
| GELU | 26,283 | 0.8982 | 0.8757 | 0.9071 | 0.4323 | 3.28 |
| RELU | 26,283 | 0.7824 | 0.7965 | 0.8497 | 0.5498 | 2.58 |
| SELU | 26,149 | 0.7880 | 0.7453 | 0.8047 | 0.6267 | 2.26 |
| **(m) Study 6.2 — Pooling strategy** | | | | | | |
| **Configuration** | **Params** | **Train R²** | **Val R²** | **Test R²** | **Test RMSEP** | **Test RPD** |
| **2 layer max pool 2** | **730,529** | **0.9607** | **0.9594** | **0.9614** | **0.2787** | **5.09** |
| no pool | 1,444,673 | 0.9530 | 0.9565 | 0.9627 | 0.2739 | 5.18 |
| avg pool 2 | 727,873 | 0.9548 | 0.9551 | 0.9622 | 0.2758 | 5.14 |
| max pool 2 | 727,873 | 0.9460 | 0.9533 | 0.9581 | 0.2903 | 4.88 |
| max pool 4 | 369,473 | 0.9386 | 0.9486 | 0.9535 | 0.3058 | 4.64 |

Note: In each study, one factor was altered at a time based on the initial working configuration and experiments on the wheat protein (NIRS5000) dataset were conducted only once. Within each study, the configuration with the highest validation R^2^ was shown in bold and highlighted. Parameters refer to the number of trainable parameters. R^2^, coefficient of determination; RMSE, root mean square error; RPD, the ratio of performance to bias. The final architecture (Table 2) was selected from these studies based on the principle of parsimony, prioritizing the simplest configuration that also achieves performance close to the optimal.

**Table S6.** The performance of CNN-Baseline model evaluated on the test set across six datasets under three input conditions.

| **Dataset** | **Input** | **Preprocessing** | **Wave-**  **lengths** | **Test R²** | **Test**  **RMSEP** | **Test**  **MAE** | **Test**  **RPD** | **Test**  **RER** |
| --- | --- | --- | --- | --- | --- | --- | --- | --- |
| (a) Wheat protein (NIRS5000, n = 5046) | Raw | none | 700 | 0.9634 ± 0.0026 | 0.2711 ± 0.0096 | 0.2126 ± 0.0088 | 5.24 ± 0.18 | 51.02 ± 1.80 |
|  | Preprocessed | snv-fd | 700 | 0.9613 ± 0.0017 | 0.2789 ± 0.0061 | 0.2172 ± 0.0046 | 5.09 ± 0.11 | 49.56 ± 1.07 |
|  | Preprocessed + wavelengths | snv-fd | 175 | 0.9632 ± 0.0017 | 0.2719 ± 0.0062 | 0.2093 ± 0.0050 | 5.22 ± 0.12 | 50.83 ± 1.14 |
| (b) Barley protein (NIRS5000, n = 2096) | Raw | none | 700 | 0.9596 ± 0.0064 | 0.3187 ± 0.0232 | 0.2474 ± 0.0168 | 5.01 ± 0.31 | 28.40 ± 1.77 |
|  | Preprocessed | fd-snv | 700 | 0.9599 ± 0.0045 | 0.3178 ± 0.0180 | 0.2443 ± 0.0140 | 5.02 ± 0.28 | 28.44 ± 1.60 |
|  | Preprocessed + wavelengths | fd-snv | 700 | 0.9604 ± 0.0041 | 0.3160 ± 0.0165 | 0.2428 ± 0.0119 | 5.05 ± 0.26 | 28.58 ± 1.49 |
| (c) Wheat protein (XDS, n = 500) | Raw | none | 1050 | 0.8464 ± 0.0438 | 0.4161 ± 0.0623 | 0.3166 ± 0.0470 | 2.64 ± 0.44 | 11.15 ± 1.84 |
|  | Preprocessed | snv-fd | 1050 | 0.8804 ± 0.0233 | 0.3694 ± 0.0356 | 0.2727 ± 0.0272 | 2.93 ± 0.28 | 12.37 ± 1.17 |
|  | Preprocessed + wavelengths | snv-fd | 272 | 0.8972 ± 0.0183 | 0.3427 ± 0.0306 | 0.2501 ± 0.0247 | 3.16 ± 0.28 | 13.32 ± 1.20 |
| (d) Wheat moisture (NIRS5000, n = 3305) | Raw | none | 700 | 0.9514 ± 0.0054 | 0.2730 ± 0.0148 | 0.2057 ± 0.0128 | 4.55 ± 0.24 | 26.52 ± 1.39 |
|  | Preprocessed | fd-snv | 700 | 0.9617 ± 0.0025 | 0.2424 ± 0.0078 | 0.1796 ± 0.0065 | 5.12 ± 0.17 | 29.82 ± 0.97 |
|  | Preprocessed + wavelengths | fd-snv | 192 | 0.9582 ± 0.0038 | 0.2533 ± 0.0115 | 0.1885 ± 0.0076 | 4.90 ± 0.22 | 28.56 ± 1.26 |
| (e) Barley moisture (NIRS5000, n = 985) | Raw | none | 700 | 0.9705 ± 0.0066 | 0.2074 ± 0.0232 | 0.1610 ± 0.0191 | 5.93 ± 0.67 | 32.54 ± 3.65 |
|  | Preprocessed | snv-fd | 700 | 0.9758 ± 0.0048 | 0.1881 ± 0.0187 | 0.1376 ± 0.0093 | 6.52 ± 0.67 | 35.80 ± 3.67 |
|  | Preprocessed + wavelengths | snv-fd | 196 | 0.9760 ± 0.0035 | 0.1878 ± 0.0137 | 0.1426 ± 0.0107 | 6.51 ± 0.49 | 35.70 ± 2.67 |
| (f) Maize moisture (NIRS5000, n = 585) | Raw | none | 700 | 0.7954 ± 0.0444 | 0.6945 ± 0.0723 | 0.4899 ± 0.0588 | 2.24 ± 0.22 | 10.61 ± 1.02 |
|  | Preprocessed | sd | 700 | 0.8424 ± 0.0324 | 0.6097 ± 0.0618 | 0.4313 ± 0.0395 | 2.56 ± 0.26 | 12.09 ± 1.22 |
|  | Preprocessed + wavelengths | sd | 621 | 0.8338 ± 0.0389 | 0.6252 ± 0.0720 | 0.4438 ± 0.0486 | 2.50 ± 0.28 | 11.82 ± 1.34 |

Note: Values are mean ± standard deviation over 25 seeds (42-66), which vary the weight initialization. The data partition is held fixed. Three inputs are compared: raw spectra with column-wise scaling only; the optimal preprocessing over the full spectrum; and preprocessing combined with the PLSR-derived wavelength subset. Unlike SVR and XGBoost, which adopted the strategy carried forward by PLSR, the preprocessing for the convolutional networks was screened independently for each dataset across the twelve strategies of Table S1, each evaluated over three seeds and ranked by validation R². On barley protein, the validation-based wavelength selection retained the full spectrum, so the third row of those blocks differs from the second only through the numerical path taken, not through the information available to the model. R^2^, coefficient of determination; RMSEP, root mean square error of prediction; MAE, mean absolute error; RPD, the ratio of performance to bias; RER, the range error ratio.

**Table S7.** Architectures selected by random search for CNN-RS under each of the three input variants.

| **Dataset** | **Input variant** | **Preprocessing** | **Wave-**  **lengths** | **Conv**  **layers** | **Filters** | **Kernels** | **Pool** | **FC layers** | **Batch**  **norm** | **Activa-**  **tion** | **Drop-**  **out** | **Learning**  **rate** | **Weight**  **decay** |
| --- | --- | --- | --- | --- | --- | --- | --- | --- | --- | --- | --- | --- | --- |
| Wheat protein (NIRS5000, n=5046) | Raw | none | 700 | **3** | 4, 8, 128 | 15, 9, 7 | max | 32, 32 | yes | SELU | 0.10 | 0.00360 | 0 |
|  | Preprocessed | SNV-FD | 700 | **3** | 4, 8, 128 | 15, 9, 7 | max | 32, 32 | yes | SELU | 0.10 | 0.00360 | 0 |
|  | Preprocessed +  wavelength selection | SNV-FD | 175 | **1** | 1 | 15 | max | 64, 32 | no | SELU | 0.00 | 0.00983 | 0 |
| Barley protein (NIRS5000, n=2096) | Raw | none | 700 | **3** | 4, 8, 128 | 15, 9, 7 | max | 32, 32 | yes | SELU | 0.10 | 0.00360 | 0 |
|  | Preprocessed | FD-SNV | 700 | **1** | 1 | 15 | max | 64, 32 | no | SELU | 0.00 | 0.00983 | 0 |
|  | Preprocessed +  wavelength selection | FD-SNV | 700 | **3** | 4, 8, 128 | 15, 9, 7 | max | 32, 32 | yes | SELU | 0.10 | 0.00360 | 0 |
| Wheat protein (XDS, n=500) | Raw | none | 1050 | **2** | 2, 16 | 21, 11 | max | 128 | yes | ELU | 0.00 | 0.00718 | 0.003 |
|  | Preprocessed | SNV-FD | 1050 | **2** | 32, 64 | 7, 5 | max | 128 | no | Leaky ReLU | 0.05 | 0.00005 | 0.002 |
|  | Preprocessed +  wavelength selection | SNV-FD | 272 | **1** | 4 | 7 | avg | 128 | no | Leaky ReLU | 0.00 | 0.00075 | 0.012 |
| Wheat moisture (NIRS5000, n=3305) | Raw | none | 700 | **3** | 4, 8, 128 | 15, 9, 7 | max | 32, 32 | yes | SELU | 0.10 | 0.00360 | 0 |
|  | Preprocessed | FD-SNV | 700 | **1** | 1 | 15 | max | 64, 32 | no | SELU | 0.00 | 0.00983 | 0 |
|  | Preprocessed +  wavelength selection | FD-SNV | 192 | **3** | 4, 8, 128 | 15, 9, 7 | max | 32, 32 | yes | SELU | 0.10 | 0.00360 | 0 |
| Barley moisture (NIRS5000, n=985) | Raw | none | 700 | **1** | 1 | 15 | max | 64, 32 | no | SELU | 0.00 | 0.00983 | 0 |
|  | Preprocessed | SNV-FD | 700 | **1** | 1 | 15 | max | 64, 32 | no | SELU | 0.00 | 0.00983 | 0 |
|  | Preprocessed +  wavelength selection | SNV-FD | 196 | **1** | 1 | 15 | max | 64, 32 | no | SELU | 0.00 | 0.00983 | 0 |
| Maize moisture (NIRS5000, n=585) | Raw | none | 700 | **3** | 4, 8, 128 | 15, 9, 7 | max | 32, 32 | yes | SELU | 0.10 | 0.00360 | 0 |
|  | Preprocessed | SD | 700 | **3** | 4, 64, 8 | 21, 11, 7 | max | 64 | no | ELU | 0.05 | 0.00052 | 0.001 |
|  | Preprocessed +  wavelength selection | SD | 621 | **3** | 4, 64, 8 | 21, 11, 7 | max | 64 | no | ELU | 0.05 | 0.00052 | 0.001 |

Note: All searches utilized 300 trails with a fixed sampling seed, so the same 300 candidate configurations were evaluated in every dataset and ranked differed. Pooling size was 2 throughout. Batch size was 32 except where the selected configurations specified 64 (wheat protein XDS, raw and preprocessed; maize moisture, preprocessed and fully optimized). Across the eighteen searches only six distinct architectures were ever selected, and two of them account for thirteen of the eighteen. The selected depth does not track the input variant: relative to the raw input, the number of convolutional layers fell in two datasets and was unchanged in four. Together with the finding that the architecture ranked best on the validation set was not the best over 25 seeds, this indicates that selection among the leading candidates is driven largely by validation-set noise rather than by a genuine input-dependent optimum. The preprocessing column gives the strategy screened independently for the CNNs, which is not in general the strategy selected by PLSR for the same dataset.

**Table S8.** The performance of CNN-RS on the test set across the six datasets under three input conditions.

| **Dataset** | **Input** | **Preprocessing** | **Wave-**  **lengths** | **Test R²** | **Test**  **RMSEP** | **Test**  **MAE** | **Test**  **RPD** | **Test**  **RER** |
| --- | --- | --- | --- | --- | --- | --- | --- | --- |
| (a) Wheat protein (NIRS5000, n = 5046) | Raw | none | 700 | 0.9662 ± 0.0011 | 0.2608 ± 0.0044 | 0.2008 ± 0.0035 | 5.44 ± 0.09 | 52.99 ± 0.90 |
|  | Preprocessed | snv-fd | 700 | 0.9640 ± 0.0013 | 0.2690 ± 0.0048 | 0.2076 ± 0.0038 | 5.27 ± 0.09 | 51.38 ± 0.92 |
|  | Preprocessed + wavelengths | snv-fd | 175 | 0.9604 ± 0.0019 | 0.2822 ± 0.0068 | 0.2175 ± 0.0053 | 5.03 ± 0.12 | 48.98 ± 1.18 |
| (b) Barley protein (NIRS5000, n = 2096) | Raw | none | 700 | 0.9662 ± 0.0023 | 0.2924 ± 0.0098 | 0.2255 ± 0.0080 | 5.44 ± 0.18 | 30.85 ± 1.04 |
|  | Preprocessed | fd-snv | 700 | 0.9633 ± 0.0020 | 0.3044 ± 0.0083 | 0.2356 ± 0.0056 | 5.23 ± 0.14 | 29.63 ± 0.81 |
|  | Preprocessed + wavelengths | fd-snv | 700 | 0.9638 ± 0.0017 | 0.3024 ± 0.0072 | 0.2301 ± 0.0060 | 5.26 ± 0.13 | 29.81 ± 0.71 |
| (c) Wheat protein (XDS, n = 500) | Raw | none | 1050 | 0.9020 ± 0.0089 | 0.3355 ± 0.0153 | 0.2447 ± 0.0164 | 3.20 ± 0.15 | 13.53 ± 0.63 |
|  | Preprocessed | snv-fd | 1050 | 0.9195 ± 0.0059 | 0.3043 ± 0.0112 | 0.2010 ± 0.0115 | 3.53 ± 0.13 | 14.91 ± 0.55 |
|  | Preprocessed + wavelengths | snv-fd | 272 | 0.9368 ± 0.0054 | 0.2695 ± 0.0115 | 0.1863 ± 0.0117 | 3.99 ± 0.17 | 16.84 ± 0.72 |
| (d) Wheat moisture (NIRS5000, n = 3305) | Raw | none | 700 | 0.9681 ± 0.0016 | 0.2213 ± 0.0058 | 0.1639 ± 0.0045 | 5.61 ± 0.15 | 32.65 ± 0.86 |
|  | Preprocessed | fd-snv | 700 | 0.9656 ± 0.0029 | 0.2296 ± 0.0095 | 0.1715 ± 0.0076 | 5.41 ± 0.22 | 31.49 ± 1.29 |
|  | Preprocessed + wavelengths | fd-snv | 192 | 0.9585 ± 0.0017 | 0.2525 ± 0.0051 | 0.1876 ± 0.0050 | 4.91 ± 0.10 | 28.61 ± 0.59 |
| (e) Barley moisture (NIRS5000, n = 985) | Raw | none | 700 | 0.9795 ± 0.0041 | 0.1734 ± 0.0166 | 0.1289 ± 0.0109 | 7.07 ± 0.64 | 38.78 ± 3.54 |
|  | Preprocessed | snv-fd | 700 | 0.9836 ± 0.0023 | 0.1552 ± 0.0107 | 0.1170 ± 0.0073 | 7.87 ± 0.54 | 43.16 ± 2.94 |
|  | Preprocessed + wavelengths | snv-fd | 196 | 0.9829 ± 0.0020 | 0.1586 ± 0.0094 | 0.1228 ± 0.0084 | 7.69 ± 0.45 | 42.18 ± 2.47 |
| (f) Maize moisture (NIRS5000, n = 585) | Raw | none | 700 | 0.8178 ± 0.0251 | 0.6573 ± 0.0448 | 0.4697 ± 0.0331 | 2.36 ± 0.16 | 11.16 ± 0.74 |
|  | Preprocessed | sd | 700 | 0.8724 ± 0.0125 | 0.5507 ± 0.0270 | 0.3900 ± 0.0193 | 2.81 ± 0.14 | 13.29 ± 0.65 |
|  | Preprocessed + wavelengths | sd | 621 | 0.8670 ± 0.0128 | 0.5621 ± 0.0273 | 0.3982 ± 0.0246 | 2.75 ± 0.14 | 13.02 ± 0.64 |

Note: Values are mean ± standard deviation over 25 seeds (42-66), which vary the weight initialization. The data partition is held fixed. Three inputs are compared: raw spectra with column-wise scaling only; the optimal preprocessing over the full spectrum; and preprocessing combined with the PLSR-derived wavelength subset. The preprocessing for the convolutional networks was screened independently for each dataset across the twelve strategies of Table S1, each evaluated over three seeds and ranked by validation R². On barley protein, the validation-based wavelength selection retained the full spectrum, so the third row of those blocks differs from the second only through the numerical path taken, not through the information available to the model. The architecture search was repeated independently within each input condition, so the three rows of a block correspond to three separately selected architectures (Table S7). R^2^, coefficient of determination; RMSEP, root mean square error of prediction; MAE, mean absolute error; RPD, the ratio of performance to bias; RER, the range error ratio.
